# Identifiability analysis for models of the translation kinetics after mRNA transfection

**DOI:** 10.1101/2021.05.18.444633

**Authors:** Susanne Pieschner, Jan Hasenauer, Christiane Fuchs

## Abstract

Mechanistic models are a powerful tool to gain insights into biological processes. The parameters of such models, e.g. kinetic rate constants, usually cannot be measured directly but need to be inferred from experimental data. In this article, we study dynamical models of the translation kinetics after mRNA transfection and analyze their parameter identifiability. Previous studies have considered ordinary differential equation (ODE) models of the process, and here we formulate a stochastic differential equation (SDE) model. For both model types, we consider structural identifiability based on the model equations and practical identifiability based on simulated as well as experimental data and find that the SDE model provides better parameter identifiability than the ODE model. Moreover, our analysis shows that even for those parameters of the ODE model that are considered to be identifiable, the obtained estimates are sometimes unreliable. Overall, our study clearly demonstrates the relevance of considering different modeling approaches and that stochastic models can provide more reliable and informative results.

## 1 Introduction

mRNA transfection is the process of introducing mRNA into a living cell. mRNA delivery has become increasingly interesting for biomedical applications because it enables treatment of diseases by means of targeted expression of proteins and it is transient, avoiding the risk of permanently integrating into the genome (see e.g. Sahin et al., 2014). One of the most prominent applications of mRNA transfection at the moment are the mRNA-based vaccine candidates that are already in use or currently under investigation to prevent COVID-19 infections (Borah et al., 2021, DeFrancesco, 2020). In such a context, it is, of course, very import to have a precise understanding of the dynamics of the underlying processes in order to be able to control them. Yet, many aspects and the determinants of the mRNA delivery process and the translation kinetics are difficult to measure and therefore poorly understood.

We aim at facilitating insights into these aspects through the use of mechanistic modeling and parameter inference for such models from experimental data. The data comes from an mRNA transfection experiment using fluorescence reporters and fluorescence microscopy which is one of the few ways to measure quantities within a living cell over time (i.e. keeping it alive is necessary). Due to the discrete nature of the molecular species within a cell and due to the fact that random fluctuations play a key role (Elowitz et al., 2002, Raj & van Oudenaarden, 2008), a continuous-time, discrete-space Markov process, also called a *Markov jump process (MJP)*, for which the dynamics are described by the so-called chemical master equation (CME), is widely accepted to be an appropriate stochastic description of the biochemical processes within a cell (Gillespie, 1992, Schnoerr et al., 2017). However, parameter inference for MJPs is computationally very demanding and often infeasible (see e. g. Warne et al., 2019). Therefore, several other representations of the biochemical kinetics have been developed. To some extent those can be considered as approximations to the corresponding MJP. The most commonly used representation is the *reaction rate equation (RRE)* which is a system of *ordinary differential equations (ODEs)* and thus provides a deterministic and state-continuous description of the kinetics. One approach that preserves the stochastic nature of the underlying process is the approximation by *Itô diffusion processes*. These are continuous-time, continuous-space stochastic processes described by *Itô-type stochastic differential equations (SDEs)*.

Single-cell fluorescence data from transfection experiments has been analyzed based on ODE modeling in several previous studies e. g. Ligon et al. (2014), Leonhardt et al. (2014), Fröhlich et al. (2018), and Reiser et al. (2019). Yet, several parameters of the considered ODE models for the translation kinetics after mRNA transfection are not identifiable from the experimental data. Moreover, the quality of the parameter estimates, i. e. whether the true kinetic rates are adequately captured, is unclear. Using models that explicitly account for stochasticity can help improve our ability to determine kinetic parameters from experimental data (Munsky et al., 2009). Browning et al. (2020) have recently compared parameter identifiability for ODE and SDE modeling approaches for example models from different contexts based on simulated data and showed that SDE modeling does improve the identifiability for the studied models.

Here, we formulate an SDE model of the translation kinetics after mRNA transfection and compare its parameter identifiability to that of the corresponding ODE model. Inference from fluorescence data for SDE models has also been conducted e. g. in Heron et al. (2007), Finkenstädt et al. (2008), and Komorowski et al. (2009), however for an experimental setup that also included the transcription process. Finkenstädt et al. (2008) even considered an SDE and an ODE model in one of their case studies, but their results did not directly show any differences in the parameter identifiability and the study was not focused on this aspect.

This article is structured as follows: In Section 2, we describe the experimental data that motivated the study. In Section 3, we present the biochemical reaction network which we want to consider for the translation kinetics after mRNA transfection and formulate its ODE and SDE representation. After stating the assumed model of the observations and summarizing the parameters that we would like to infer from these observations in Section 4, we consider several approaches to assess structural identifiability of the parameters for both model types in Section 5. We define the parameter posteriors for both modeling approaches and study the practical identifiability of the parameters based on simulated and experimental data in Section 6, and conclude with a summary and discussion of our findings in Section 7. All relevant code to perform the presented analysis is available on Github: https://github.com/fuchslab/Translation_kinetics_after_transfection.

## 2 Experimental data

We consider data that has previously been analyzed (based on ODE modeling) and published in Fröhlich et al. (2018). The data was generated in an experiment where human hepatoma epithelial cells from the cell line HuH7 were transfected with mRNA encoding a green fluorescent protein (GFP). The cells were fixed on micro patterned protein arrays and time lapse microscopy images of the cells were taken every 10 minutes over the course of at least 30 hours (i.e. there are at least 180 measurements per cell). For the first hour, mRNA lipoplexes were added. Afterwards, the cells were washed with cell culture medium such that no further lipoplex uptake occurred. The time point at which the lipoplexes were taken up, dissolved and released the mRNA as well as the number of mRNA molecules released are unknown.

The released mRNA was translated into a fluorescent protein which caused the cells to fluoresce. For each image taken during the experiment, the fluorescence intensity is integrated over squares occupied by one cell in order to obtain one value for the mean fluorescence intensity per cell and time point (see Fröhlich et al., 2018, for further details about the image analysis).

The experiment was conducted with two different types of GFP that differ in their protein lifetime: enhanced GFP (eGFP) and a destabilized enhanced GFP (d2eGFP). The data set contains measurements for more than 800 cells for each type of GFP.

Some trajectories of the mean fluorescence intensity are displayed in Figure 1.

**Figure 1:**
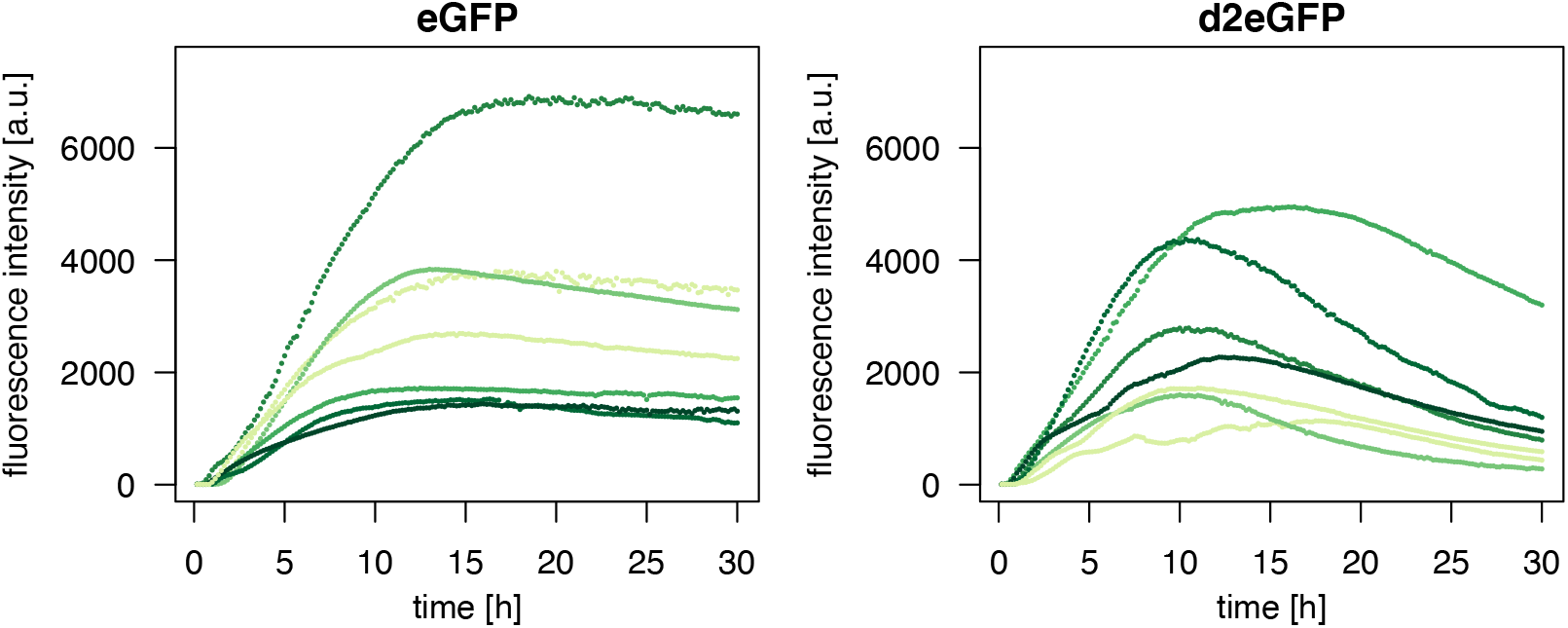
Trajectories of the mean fluorescence intensity for seven cells from the mRNA transfection experiment in Fröhlich et al. (2018) for eGFP and d2eGFP (April 27, 2016), respectively.

It was shown before that ODE models of the translation kinetics of an individual cell are not globally identifiable with the available experimental data as described above. Several of the ODE model parameters cannot be uniquely determined based on one observed fluorescence trajectory. Fröhlich et al. (2018) use a mixed-effect ODE model in order to incorporate the translation kinetics of several cells and data for both different types of GFP (eGFP and d2eGFP). Through this approach, they are able to improve parameter identifiability (by breaking the symmetry between the degradation rate constants); however, their approach is computationally very intense, necessitates distribution assumptions about the random effects, required conducting the experiment with two types of GFP, and still leaves several parameters non-identifiable. Here, we are interested in the question whether the use of an SDE model can improve the parameter identifiability even when only one fluorescence trajectory is observed.

## 3 Mathematical models of the translation kinetics

We will focus on modeling the translation kinetics of one cell in order to study parameter identifiability based on one observed fluorescence trajectory. Therefore, we only consider the (released) mRNA and the GFP molecules explicitly:

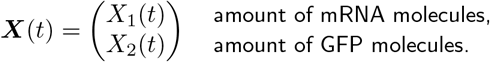

We assume that all mRNA molecules (within one cell) are released at once from the lipoplexes and denote this initial time point by *t*_0_. Before *t*_0_, there are neither mRNA nor GFP molecules, and at *t*_0_, an amount of *m*_0_ mRNA molecules is released, i.e.

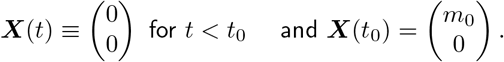

Conceivable extensions of this basic model are e. g. to include enzymatic degradation of the mRNA and/or the protein, ribosomal binding to the mRNA for translation, and a maturation step of the protein. However, we will only consider the basic configuration as described above.

### 3.1 Markov Jump Process

Assuming that the matter within the cell is well-stirred and in thermal equilibrium, an MJP is regarded to be the most adequate representation of this system after *t*_0_. In our model, there are three possible reactions:

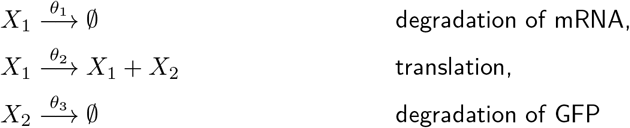

with kinetic rate parameters *θ*_1_, *θ*_2_, and *θ*_3_. Given the state vector ***X***(*t*) = (*X*_1_(*t*),*X*_2_(*t*))^*T*^, the propensity functions of the three reactions are

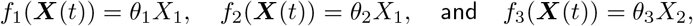

i.e. the probability that reaction *i* occurs within the next time interval of length Δ*t* is approximately *f_i_*(***X***(*t*))Δ*t* for *i* = 1, 2, 3.

If we denote the probability distribution of the random variable ***X***(*t*) by

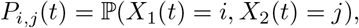

the corresponding CME reads

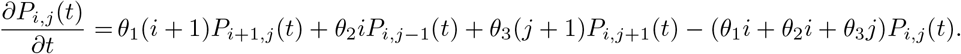

Although the system contains only first-order reactions, there is no known closed-form solution to the CME. Thus, there is no explicit formula for the transition probability distribution *p*(***X***(*t*)|***X***(*s*),*θ*) for *s* < *t*. This fact impedes efficient inference for this MJP and leads to the consideration of other modeling approaches.

### 3.2 ODE model

The RRE is a system of ODEs that provides a deterministic and state-continuous description of biochemical kinetics and can be derived as the large-volume limit of the corresponding MJP (see e. g. Kurtz, 1972). The following system of ODEs is a deterministic approximation of the MJP modeling the dynamics as described above:

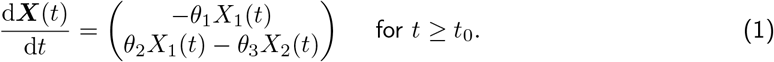

This system has previously been formulated and analyzed e. g. in Leonhardt et al. (2014). It admits the solution

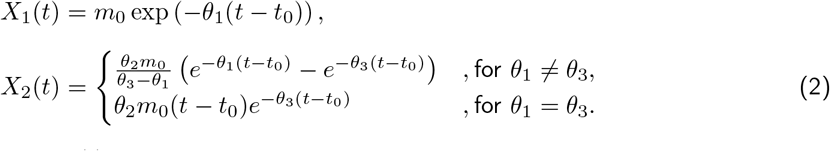

Note that the solution for *X*_2_(*t*) is symmetric in the parameters *θ*_1_ and *θ*_3_. Moreover, note that while it is common for the RRE to describe molecule concentrations; here, we let ***X***(*t*) denote molecule numbers. However, since the model that we consider here contains only first-order reactions, this does not affect the interpretation of the kinetic rate parameters (i. e. Equation (1) would look the same for concentrations).

### 3.3 SDE model

While the ODE model of the translation kinetics after mRNA transfection had been considered before, we are the first to formulate the stochastic but state-continuous approximation to the MJP in Section 3.1 based on an (Itô) diffusion process. The approximating diffusion process is described by an SDE which is known as the chemical Langevin equation (CLE) in the context of biochemical kinetics (Gillespie, 2000). The CLE is derived based on the changes in molecule numbers caused by the three possible reactions and on the corresponding propensities and reads:

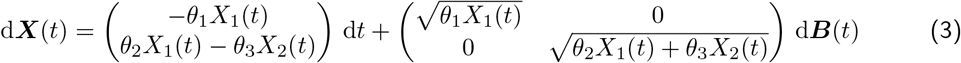

for *t* ≥ *t*_0_ and where ***B***(*t*) is a 2-dimensional standard Brownian motion. The solution of Equation (3) is not explicitly known and will be approximated based on the Euler-Maruyama scheme as explained in Section A.1 of the supplementary material. See Fuchs (2013) for a concise introduction to diffusion processes and several approaches to derive general diffusion approximations.

## 4 Model parameters and observations

### 4.1 Model of the observations

In the experiment described in Section 2, neither the amount of mRNA molecules nor that of GFP molecules can be measured directly. Instead, a fluorescence signal is observed which is composed of a background fluorescence and a signal which is proportional to the amount of GFP molecules. Moreover, previous studies assumed a constant offset and multiplicative measurement noise in the recorded fluorescence trajectories (Fröhlich et al., 2018). Therefore, denoting a trajectory of mean fluorescence intensity observed at time points *t_k_*, for *k* = 1,…,*K*, by {*y_k_*}_*k*=1,…,*K*_, we assume that

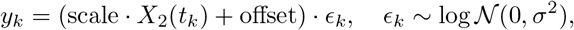

where the random variables ϵ*_k_* are independent.

Note that the observations depend only on the amount *X*_2_ of GFP molecules, but not directly on the amount *X*_1_ of mRNA molecules.

### 4.2 Model parameters

Based on the observations {*y_k_*}_*k*=1,…,*K*_, we aim to infer the following unknown parameters:

- the three kinetic parameters ***θ*** = (*θ*_1_,*θ*_2_,*θ*_3_) that denote the rate constants for mRNA degradation, translation, and GFP degradation,
- the initial amount *m*_0_ of mRNA molecules and the time point *t*_0_ at which it is released,
- the scaling factor scale and the offset for the fluorescence signal,
- and the standard deviation *σ* of the measurement errors.

## 5 Structural identifiability analysis

Our main interest lies in the question which of the model parameters for our two model types (ODE and SDE) can be inferred from the experimental data as described in Sections 2 and 4. Here, we first focus on the parameters **θ**, *m*_0_, scale, and offset that drive the dynamics of the process and the fluorescence signal. We analyze the structural identifiability which only considers the model equations of the process dynamics and the observation equation (not the actual data) and assumes that we are in a perfect data situation, i. e. we have an infinite amount of data observed without measurement error (Raue et al., 2009). Plainly speaking, structural identifiability analysis answers the question whether different parameter combinations can lead to the same model output. While for ODE models, there are analytical methods to assess structural identifiability, no such methods exit for SDE models. Therefore, we consider several approaches that do not directly assess the structural identifiability for our SDE model, but answer related questions. In the following subsections, we consider a transformed version of both model types, we make use of a surrogate model and the open source software DAISY as has recently been suggested by Browning et al. (2020), and finally we also study simulations of both model types.

### 5.1 Transformed models

We can reformulate the differential equations for both model types by setting

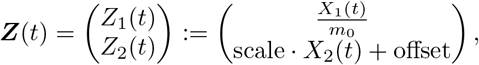

which means that

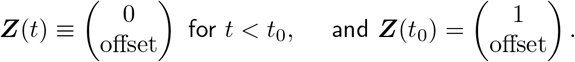

Hence, the second component of the transformed process models the fluorescence signal which we assume to be observed, meaning *y* = *Z*_2_.

#### Transformed ODE model

For the ODE model in Equation (1), the transformed model reads

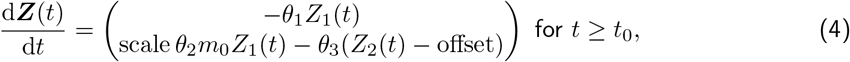

and has the solution

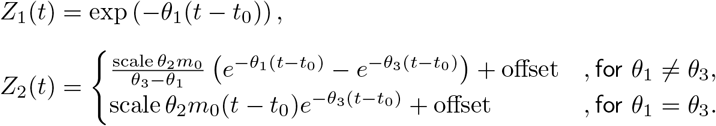

The parameters scale, *m*_0_, and *θ*_2_ appear only as a product. Thus, it can already be deduced from this equation that at most the product of the three parameters will be identifiable but not the three parameters individually (Fröhlich et al., 2018). Moreover, since only *Z*_2_(*t*) is observed and it is symmetric in the parameters *θ*_1_ and *θ*_3_ (i. e. switching their values will lead to the same model output), these two parameters can at most be locally identifiable if no further constraints are imposed on them as done in Leonhardt et al. (2014).

#### Transformed SDE model

For the SDE model in Equation (3), we apply the Itô formula (as stated in Section A.1 of the supplementary material) to obtain the transformed model

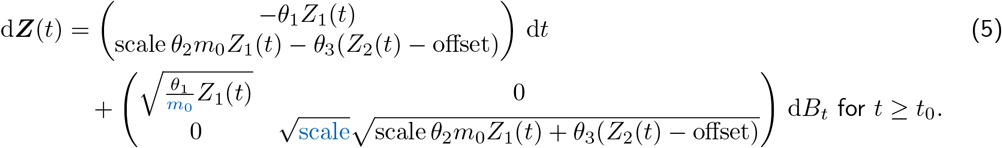

Note that here, the parameters scale and *m*_0_ also appear outside the product scale*θ*_2_*m*_0_. Furthermore, there is no apparent symmetry between *θ*_1_ and *θ*_3_. Therefore, it might be possible to gain more information about the individual parameters from data for the SDE model than for the ODE model.

### 5.2 Using a surrogate model

Next, we want to use the open source software DAISY (Differential Algebra for Identifiability of SYstems, Bellu et al., 2007) to assess the structural identifiability of the parameters in the two models of the translation kinetics. DAISY is a software tool that implements a differential algebra algorithm to perform structural identifiability analysis for systems of polynomial or rational ODEs and that also allows to include unknown initial conditions. We use the transformed models from the previous subsection for the identifiability analysis. For the ODE model in Equation (4), the analysis with DAISY is straight forward since it is intended for the use for ODE models. After applying DAISY, the obtained output shows that when considering the set of parameters {***θ**, m*_0_, scale, offset}, the model is non-identifiable. The DAISY output also reveals that this non-identifiability is due to the fact that the parameters *θ*_1_ and *θ*_3_ are only locally identifiable and the parameters *θ*_2_, *m*_0_, and scale are not individually identifiable, but only their product is. This confirms the assertions from the previous subsection. Moreover, we obtain that the remaining parameter offset is structurally identifiable.

For SDE models, Browning et al. (2020) suggest to formulate a surrogate model based on the moment equations of the diffusion process. The moment equations are a system of ODEs, and thus, DAISY can be applied to this system. For the SDE (5), let 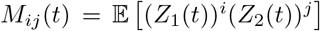 be the (mixed) moment of the diffusion process of order *i* and *j*. The moments are obtained by applying the Itô formula (as stated in Section A.1 of the supplementary material) to (*Z*_1_ (*t*))^i^(*Z*_2_(*t*))^j^ and then taking the expectation. Considering the first and the second moments of the process states results in the following system of ODEs:

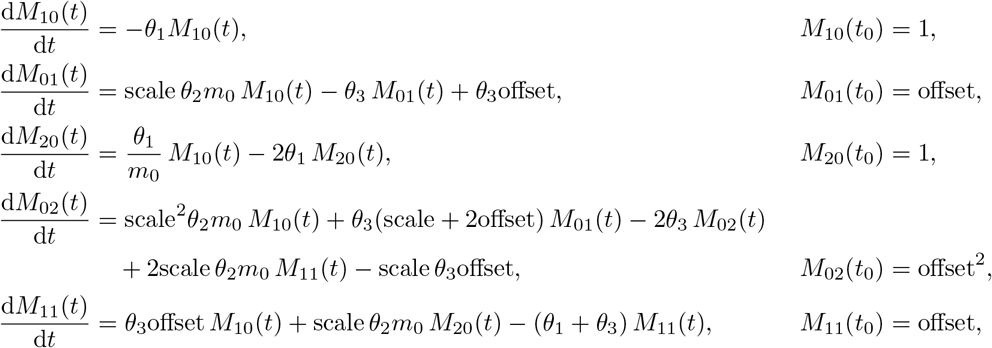

where the equations for the two first moments *M*_10_ and *M*_01_ coincide with the ODE model in Equation (4). Since in the experiment, only the fluorescence signal is observed, we consider the moments that only depend on the second component of the process, i.e. *M*_01_ and *M*_02_, as output states for the identifiability analysis. Using DAISY, we obtain that the surrogate model is globally identifiable, i. e. all six parameter values could be uniquely determined if we were able to observe the moments *M*_01_ and *M*_02_ directly, infinitely long over time, and without measurement error. While this implies that the parameters are structurally identifiable from moment measurements, it does not directly answer our original question of whether the parameters of the SDE model can be identified from perfect observations of a single fluorescence trajectory.

Moreover, as has already been pointed out, structural identifiability analysis assumes a perfect data situation. From its results, we cannot conclude that the parameters will be identifiable from the actual experimental data (which is always finite and usually subject to measurement error). In contrast, the notion of practical identifiability is concerned with the question whether the parameters can be determined from a specific data set (Raue et al., 2009) and we will address this question in Section 6.

### 5.3 Simulation-based assessment of parameter influence

Another attempt to assess parameter identifiability is to simulate from both model types for different parameter settings and compare whether we see differences in the simulation output. To obtain simulations from the ODE model, we use its solution in Equation (2). Since the ODE model is deterministic, each parameter setting yields one unique output trajectory; whereas for the SDE model, we simulate several trajectories for each parameter setting using the Euler-Maruyama scheme with a time step of 0.01 hours.

#### **Keeping the product** scale *θ*_2_*m*_0_ **constant**

As already pointed out in Subsection 5.1, the trajectories of the fluorescence intensity for the ODE model are identical if the product scale *θ*_2_*m*_0_ and the remaining parameters are fixed, even when the individual factors scale, *θ*_2_, and *m*_0_ vary. To assess whether this is (at least approximately) also the case for the SDE model, we simulate several trajectories with different values for scale, *θ*_2_, and *m*_0_ while keeping their product constant. For each parameter setting, we set the same random seed at the beginning of the simulation. To ensure that the analysis is performed in a relevant regime, we chose the parameters similar to the parameters estimated by Fröhlich et al. (2018): scale*θ*_2_*m*_0_ = 350, *θ*_1_ = 0.11, *θ*_3_ = 0.03, offset = 8.9, and *t*_0_ = 0. Figure 2 displays the simulated trajectories.

**Figure 2:**
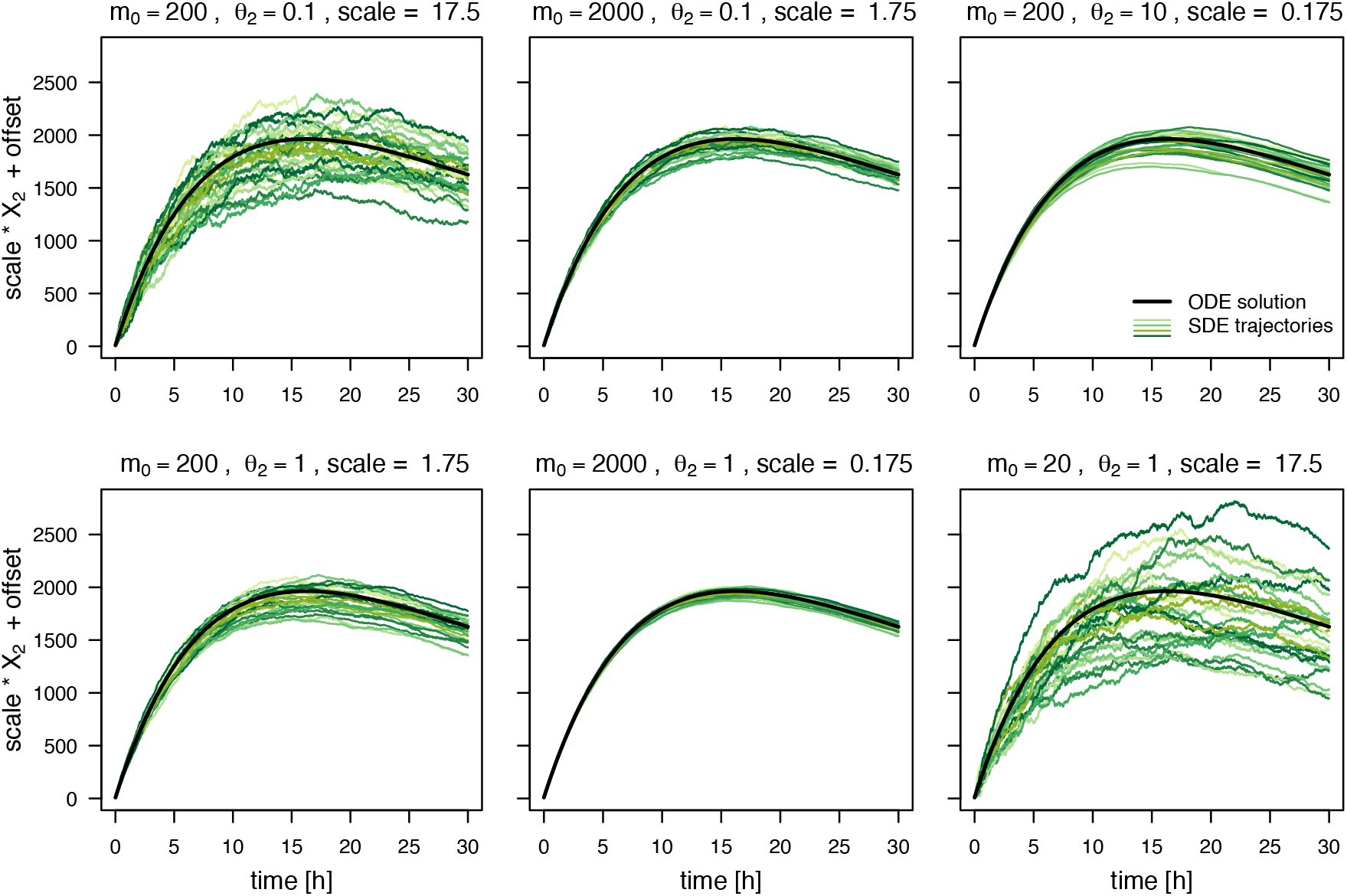
The ODE trajectory and 20 SDE trajectories of the fluorescence intensity simulated for different values of *m*_0_, *θ*_2_, and scale, while keeping their product constant at scale *θ*_2_*m*_0_ = 350. The remaining parameters are set to *θ*_1_ = 0.11, *θ*_3_ = 0.03, offset = 8.9, and *t*_0_ = 0.

It is evident that the SDE trajectories behave differently for different combinations of scale, *θ*_2_, and *m*_0_ which yield the same values of the product scale*θ*_2_*m*_0_. For example, when we keep *m*_0_ fixed while increasing scale and decreasing *θ*_2_, the variation between but also within the trajectories increases. When we keep scale fixed while decreasing *m*_0_ and increasing *θ*_2_, especially the variation between trajectories seems to increase. And finally, when we keep *θ*_2_ fixed while decreasing *m*_0_ and increasing scale, the variation between and within the trajectories increases. Our focus is on estimating the parameters from individual observed trajectories. In this context, especially the difference in the variation within the trajectories is relevant.

#### **Swapping the degradation rate constants** *θ*_1_ **and** *θ*_3_

The trajectories of the fluorescence intensity for the ODE model are also identical if the values for *θ*_1_ and *θ*_3_ are swapped while the remaining parameters are fixed. We simulate trajectories for the parameter combinations (*θ*_1_,*θ*_3_) = (0.11, 0.03) and 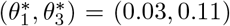, respectively, while setting the remaining parameters to scale = 17.5, *θ*_2_ = 0.1, *m*_0_ = 200, offset = 8.9, and *t*_0_ = 0. For the SDE model, we again simulate several trajectories for both parameter settings and set the same random seed at the beginning of the simulation.

Figure 3 shows the ODE trajectory and several SDE trajectories in one panel for each of the two parameter combinations separately. Whereas, Figure 4 presents one SDE trajectory for each of the two parameter combinations together in one panel. Again, the SDE trajectories do behave differently for the different parameter combinations. While there seems to be only little difference in the variation between the trajectories, the variation within the trajectories is clearly higher for lower *θ*_1_ and higher *θ*_3_. This indicates that it may be possible to uniquely determine the values of *θ*_1_ and *θ*_3_ even when estimating from only one observed trajectory.

**Figure 3:**
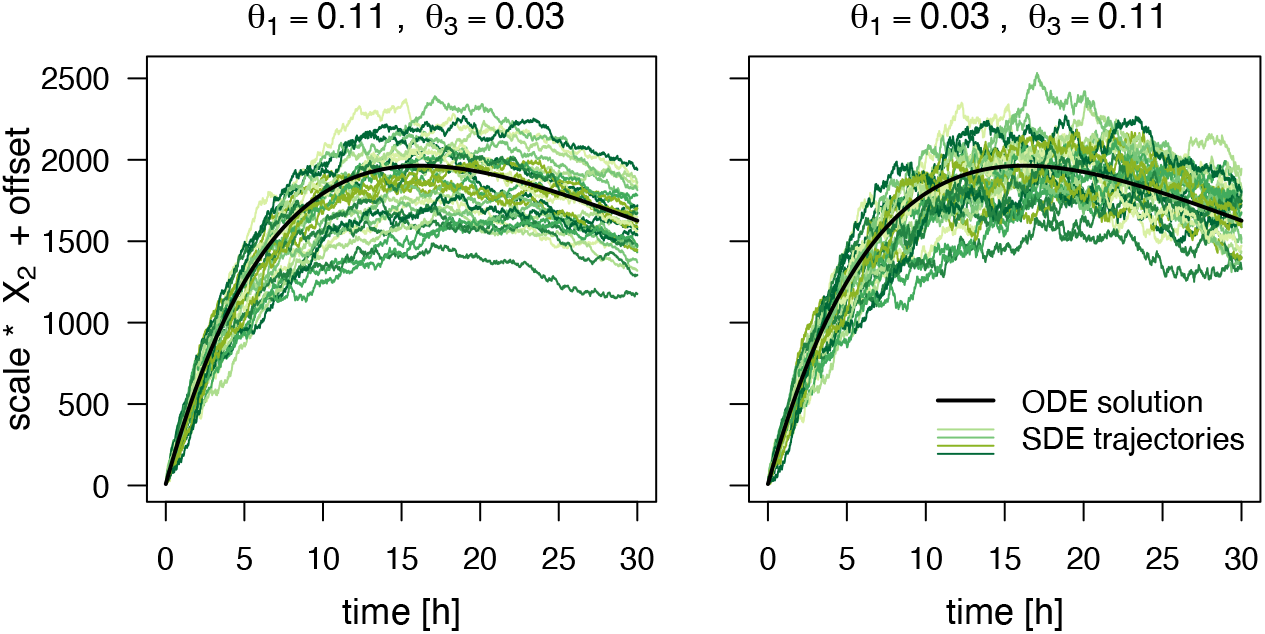
The ODE trajectory and 20 SDE trajectories of the fluorescence intensity simulated for two parameter combinations where the values of *θ*_1_ and *θ*_3_ are swapped. The remaining parameters are set to scale = 17.5, *θ*_2_ = 0.1, *m*_0_ = 200, offset = 8.9, and *t*_0_ = 0.

**Figure 4:**
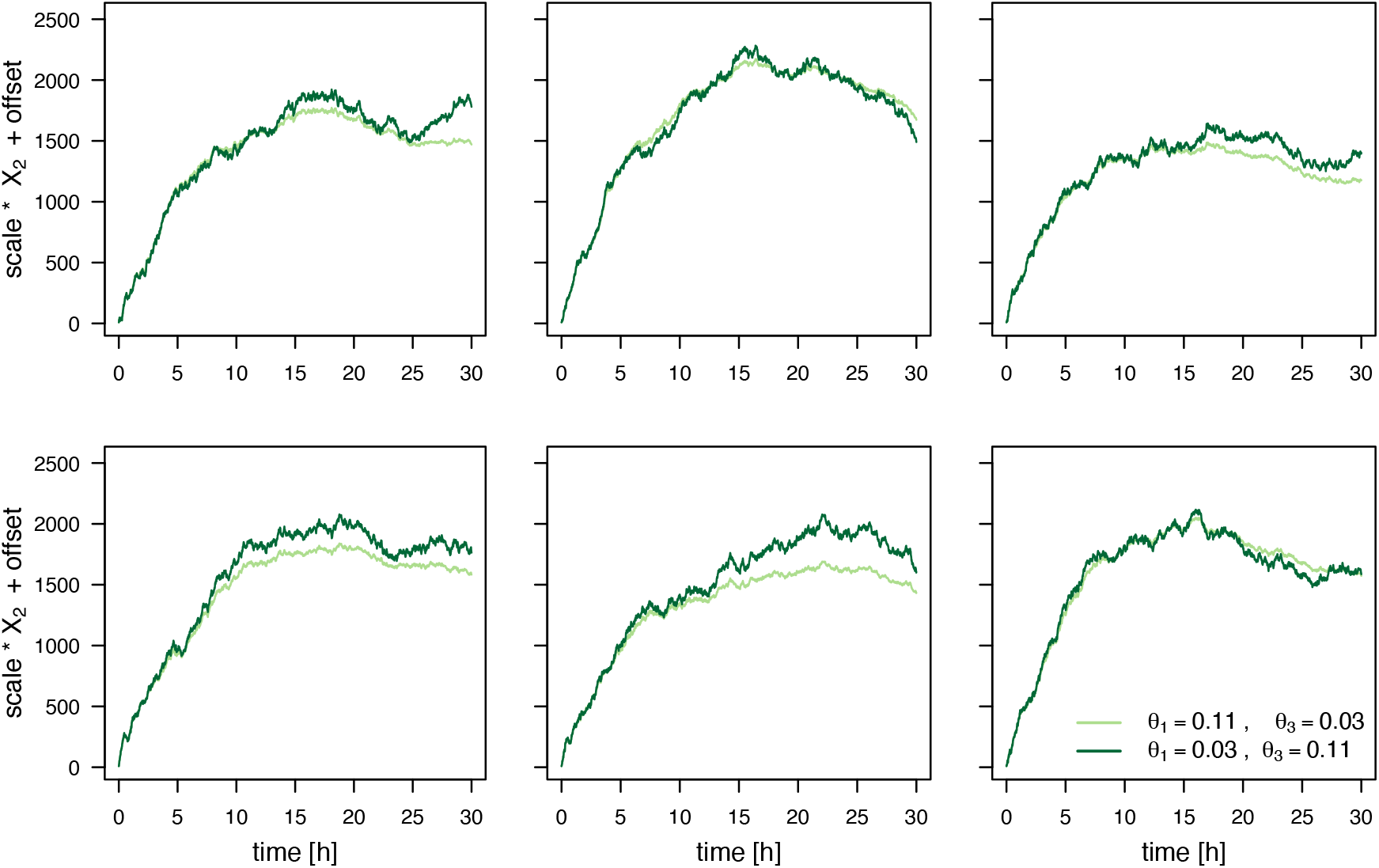
One trajectory of the fluorescence intensity for the SDE model simulated for each of the two parameter combinations where the values of *θ*_1_ and *θ*_3_ are swapped. The remaining parameters are set to scale = 17.5, *θ*_2_ = 0.1, *m*_0_ = 200, offset = 8.9, and *t*_0_ = 0.

## 6 Posterior properties and credible intervals

After having studied the structural properties of the ODE and SDE model in the previous section; next, we would like to assess the practical parameter identifiability by trying to estimate the parameters from data as described in Section 4. We take a Bayesian approach to parameter estimation because it allows for uncertainty assessment of the parameter estimates and also for handling unobserved process components and measurement error by using Markov chain Monte Carlo (MCMC) methods to sample from the parameter posterior distribution. Therefore, in this section, we define the parameter posterior densities for the two model types and study their properties based on MCMC sampling results for simulated data generated without and with (simulated) measurement error as well as for experimental data.

### 6.1 Formulation of the inference problem

In Bayesian statistics, we can formulate our assumptions and general knowledge about the model parameters 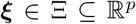 in terms of a prior distribution with probability density *p*(*ξ*). After having observed data 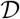 about the phenomenon which we are trying to model, we update our knowledge about the parameter and describe it by the posterior distribution with density 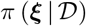. In our case, we consider data 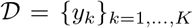 and the vector of all unknown parameters is *ξ* = (***θ**, m*_0_, scale, offset, *t*_0_, *σ*). The relation between the prior and the posterior density resulting from Bayes’ theorem is given by:

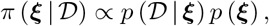

where 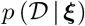 denotes the density of the distribution of 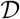 conditioned on *ξ* and is determined by the considered model. Viewed as a function of the parameter, 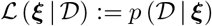 is called the likelihood (function). Comprehensive introductions to Bayesian statistics can be found e. g. in Lee (2012) and Gelman et al. (2013).

To sample from the subsequently formulated posterior densities for the two model types, we use the open source software Stan (Carpenter et al., 2017) which provides an efficient C++ implementation of the Hamiltonian Monte Carlo (HMC) based No-U-Turn Sampler (NUTS). See Section A.3 of the supplementary material for a brief introduction to this topic. We use Stan though its R interface rstan (Stan Development Team, 2019).

#### Posterior of the ODE model

For the ODE model, there is a deterministic relationship between the process values ***X***(*t*) and the parameters ***θ**, m*_0_ and *t*_0_ (or between the fluorescence signal and the parameters additionally including scale and offset, respectively).

Define the index *k*\*:= min{*k* ∈ {1, …, *K*}|*t_k_* ≥ *t*_0_} of the first observation time point after the mRNA molecules are released, then the posterior density π from which we would like to sample is proportional to

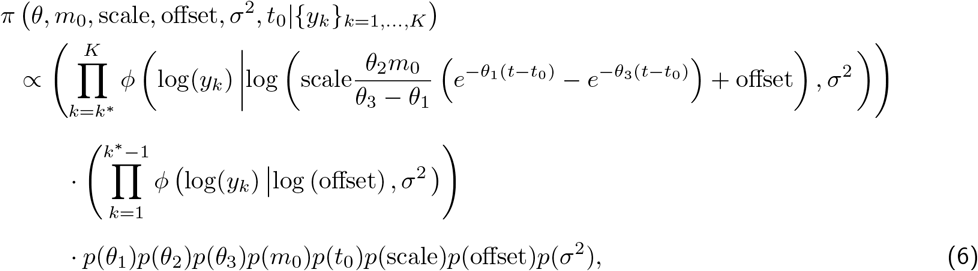

where *ϕ*(·|*μ, η*^2^) denotes the density of the normal distribution with mean *μ* and variance *η*^2^ and the *p*(·) denote the parameter prior densities which we assume to be independent.

If the priors *p*(*θ*_1_) and *p*(*θ*_3_) are symmetric to each other, then the posterior is also symmetric with respect to the two degradation rate constants.

The scaling factor scale, the translation rate constant *θ*_2_, and the initial amount of mRNA *m*_0_ appear only as a product in the likelihood function; therefore, as pointed out before, at most their product scale*θ*_2_*m*_0_ is identifiable.

#### Posterior of the SDE model

For the SDE model, the states ***X***(*t_k_*), for *k* = 1, …, *K*, of the process conditioned on the parameters ***θ**, m*_0_ and *t*_0_ are random numbers (for *t_k_* ≥ *t*_0_). Hence, we have to marginalize over the process states to obtain the posterior density of the parameters which we want to infer:

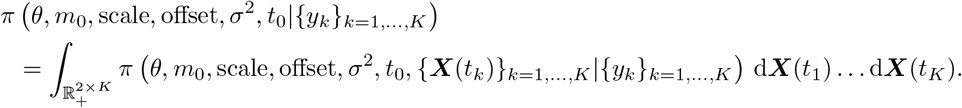

Therefore, again defining *k*\*:= min{*k* ∈ {1, …, *K*}|*t_k_* ≥ *t*_0_}, we would need to sample from

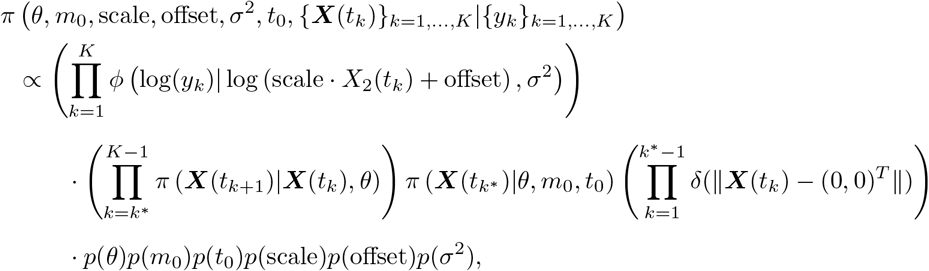

where *ϕ*(·|*μ, η*^2^) denotes the density of the normal distribution with mean *μ* and variance *η*^2^, *δ*(·) denotes the Dirac delta function, || · || denotes a norm (e. g. the *l*_2_-norm), and the factors *π* (***X***(*t*_*k*+1_)|***X***(*t_k_*), *θ*), *k* = *k\**, …, *K* — 1, denote the transition density of the process. However, the fact that the process ***X*** switches from a deterministic regime before *t*_0_ to a stochastic one after *t*_0_ complicates the estimation of *t*_0_ together with the remaining parameters. Therefore, we will assume that *t*_0_ is determined beforehand, e. g. based on the estimates for the ODE model. Consequently, we sample from

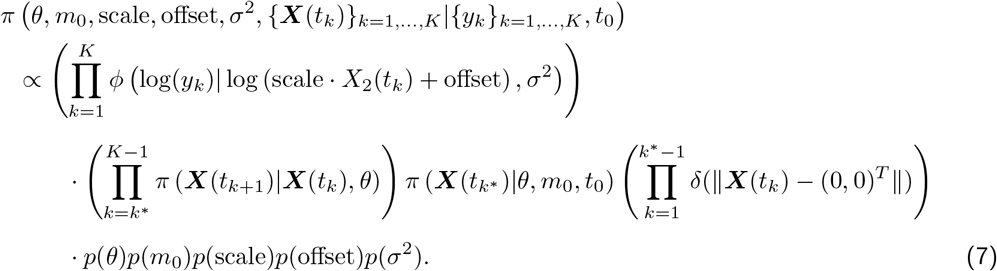

While for the ODE model, the posterior distribution is only 8-dimensional and can be sampled from directly; for the SDE model, we need to sample from a (*7* + *2K*)-dimensional distribution and then marginalize over the *2K* dimensions of the process states to obtain the posterior distribution of the parameters of interest. Moreover, there is no explicit exact expression for the transition density *π* (***X***(*t*_*k*+1_)|***X***(*t_k_*), *θ*); wherefore, it will be approximated by a normal density based on the Euler-Maruyama scheme as explained in Section A.1 of the supplementary material. For this approximation to be appropriate, we have to ensure that the time steps between observations are small enough. We do so in Section A.2 of the supplementary material.

### 6.2 Estimation based on simulated data

In order to assess how well we can recover the model parameters for both model types from individually observed trajectories, we first work with simulated data that is generated with Gillespie’s algorithm (Gillespie, 1976, 1977).

#### 6.2.1 Sampling results for simulated data without measurement error

For now, we assume the fluorescence intensity to be observed without measurement error. The data was simulated with Gillespie’s algorithm with parameters *θ* = (0.2, 0.32, 0.01), *m*_0_ = 240, *t*_0_ = 0.96, scale = 1.8, and offset = 8.5 over a time interval of 30 hours which yields trajectories similar to the experimental data, and we use the fluorescence intensity at 181 equidistant time points as observations to mirror the structure of the experimental data. The simulated fluorescence intensity (without measurement error) is depicted by the blue dotted line on the right hand side of Figure 5. We use Stan to sample from the posterior distributions of the ODE model and the SDE model given the simulated data. Since we assume the data to be observed without measurement error, the parameter offset can be determined directly from the first observation. Therefore, we do not include measurement error (and thus the parameter *σ*) and the parameter offset in the posterior distribution of the SDE model. Whereas for the ODE model, deviations of the observed data from the deterministic ODE trajectory have to be attributed to measurement error; therefore, the parameter *σ* has to be included in the posterior distribution of the ODE model. We also include the parameter offset for the ODE model in order to avoid degeneracy of the posterior. For the SDE model, we use the mean estimate of *t*_0_ obtained from the sample for the ODE model as the fixed value of *t*_0_. We assume the following prior distributions: 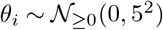 for *i* = 1, 2, 3, 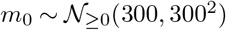, scale 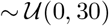, where 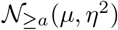 denotes the normal distribution truncated from below by a, and additionally for the ODE model, offset 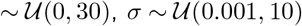, and 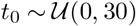.

**Figure 5:**
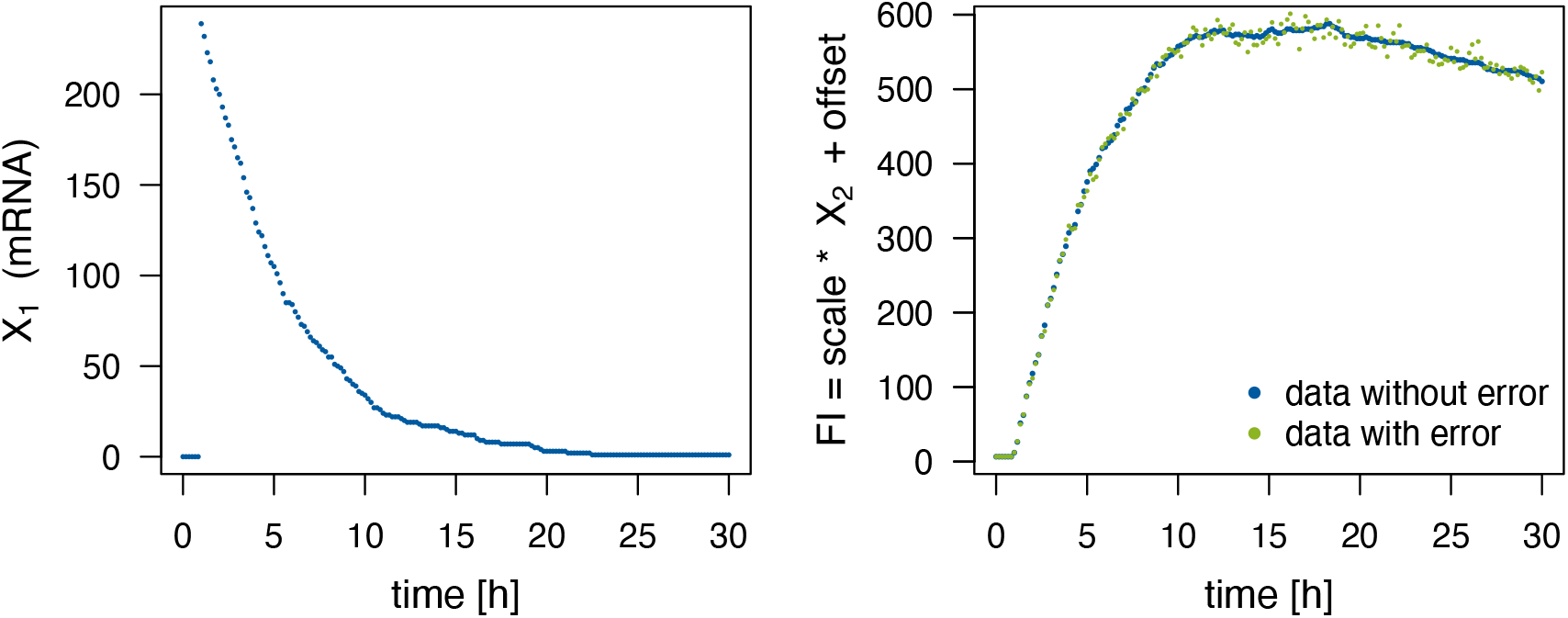
One trajectory used in the simulation study that was simulated with Gillespie’s algorithm with parameters *θ* = (0.2, 0.32, 0.01), *m*_0_ = 240, *t*_0_ = 0.96, scale = 1.8, and offset = 8.5, and for the green dotted line, multiplicative measurement error with *σ* = 0.02 was added to the fluorescence intensity (FI).

For both model types, we generate 8 HMC chains of 5000 iterations and discard the first half of the iterations as warm-up. Thus, we use a posterior sample of size 20,000 for each model type in the subsequent analysis. Tables 1 and 2 summarize the Stan output of the posterior samples for the ODE and the SDE model, respectively, and also include the true parameter values that were used to simulate the data for comparison. The tables also contain the 2.5%-, 50%-, and 97.5%-quantiles of the samples. We use the interval between the 2.5%- and the 97.5%-quantile as an estimate of the 95%-credible interval (CI). For the ODE model, we see that the parameters offset and *t*_0_ are well estimated since mean and median of the sample correspond to the true value, the CIs are very narrow, the effective sample size (ESS) *n*_eff_ is high and 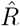 is equal to 1. As expected, the measurement error parameter *σ* is estimated to be higher than the true value of zero. Of greater interest are the remaining parameters as we can compare the results for them between the two model types.

**Table 1:** Summary of the Stan output for the ODE model given simulated data without measurement error and the true parameter values that were used to simulate the data. c.v. denotes the coefficient of variation and the columns headed by percentages contain the quantiles of the respective percentage value.

| | true value | mean | c.v. | 2.5% | 50% | 97.5% | $n_{\text{eff}}$ | $\hat{R}$ |
| --- | --- | --- | --- | --- | --- | --- | --- | --- |
| $\theta_1$ | 0.20 | 0.11 | 0.634 | 0.02 | 0.16 | 0.17 | 4 | 26.65 |
| $\theta_2$ | 0.32 | 1.52 | 1.370 | 0.02 | 0.64 | 7.56 | 12168 | 1.00 |
| $\theta_3$ | 0.01 | 0.07 | 0.943 | 0.02 | 0.02 | 0.16 | 4 | 33.00 |
| $m_0$ | 240.00 | 204.62 | 1.017 | 2.26 | 135.79 | 724.38 | 10984 | 1.00 |
| scale | 1.80 | 7.02 | 1.137 | 0.07 | 3.46 | 27.28 | 9806 | 1.00 |
| offset | 6.50 | 6.50 | 0.011 | 6.37 | 6.50 | 6.64 | 17113 | 1.00 |
| $t_0$ | 0.96 | 0.96 | 0.002 | 0.96 | 0.96 | 0.97 | 18718 | 1.00 |
| $\sigma$ | 0.00 | 0.03 | 0.054 | 0.02 | 0.03 | 0.03 | 16091 | 1.00 |
| $\theta_2 m_0$ | 76.80 | 213.08 | 2.487 | 4.57 | 35.99 | 1668.06 | 11087 | 1.00 |
| $\theta_2 \text{scale}$ | 0.58 | 6.46 | 2.875 | 0.17 | 0.92 | 55.00 | 7704 | 1.00 |
| $m_0 \text{scale}$ | 432.00 | 1033.30 | 2.181 | 16.48 | 195.96 | 7975.22 | 7899 | 1.00 |
| $\theta_2 m_0 \text{scale}$ | 138.24 | 124.67 | 0.007 | 122.96 | 124.66 | 126.38 | 13299 | 1.00 |

**Table 2:** Summary of the Stan output for the SDE model given simulated data without measurement error and the true parameter values that were used to simulate the data. The parameters *σ*, offset, and *t*_0_ are not estimated here. For the simulated data without measurement error, no measurement error is assumed for the SDE model. Thus, there is no *σ* to be estimated, and the offset is determined directly from the data and not estimated. Moreover, the initial time point *t*_0_ is predetermined based on the mean estimate of the sample for the ODE model.

| | true value | mean | c.v. | 2.5% | 50% | 97.5% | $n_{\text{eff}}$ | $\hat{R}$ |
| --- | --- | --- | --- | --- | --- | --- | --- | --- |
| $\theta_1$ | 0.20 | 0.19 | 0.108 | 0.15 | 0.19 | 0.23 | 3120 | 1.00 |
| $\theta_2$ | 0.32 | 0.39 | 0.999 | 0.09 | 0.26 | 1.48 | 206 | 1.04 |
| $\theta_3$ | 0.01 | 0.01 | 0.167 | 0.01 | 0.01 | 0.02 | 2514 | 1.00 |
| $m_0$ | 240.00 | 344.37 | 0.589 | 57.21 | 313.25 | 800.67 | 184 | 1.05 |
| scale | 1.80 | 1.66 | 0.172 | 1.19 | 1.62 | 2.30 | 3062 | 1.00 |
| $\theta_2 m_0$ | 76.80 | 82.60 | 0.178 | 55.68 | 81.69 | 113.53 | 5296 | 1.00 |
| $\theta_2 \text{scale}$ | 0.58 | 0.61 | 0.923 | 0.17 | 0.43 | 2.21 | 178 | 1.05 |
| $m_0 \text{scale}$ | 432.00 | 576.69 | 0.634 | 86.08 | 511.66 | 1440.41 | 232 | 1.04 |
| $\theta_2 m_0 \text{scale}$ | 138.24 | 133.18 | 0.083 | 112.47 | 132.74 | 156.40 | 5829 | 1.00 |

We first focus on the two degradation rate constants *θ*_1_ and *θ*_3_. Our analysis in Section 5 already showed that for the ODE model, these two parameters are only locally identifiable and the posterior distribution is symmetric with respect to them in the case of identical priors for both parameters. This is also apparent in the density plots in Figure 6. The density estimates of the posterior sample for the ODE model are clearly bimodal. The reason that the two modes are not exactly symmetric here is that HMC chains usually are only able to explore one mode and in our example 5 out of the 8 chains happen to end up in the mode where *θ*_1_ is higher than *θ*_3_ while only 3 chains converge to the other mode. The fact that each chain only samples from one of the modes is also the reason for the extremely low ESSs and the very high values of 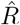 for *θ*_1_ and *θ*_3_ in Table 1. Moreover, note that neither of the modes and not even the ranges of all values in the posterior sample cover the true parameter values of *θ*_1_ and *θ*_3_. For the SDE model on the other hand, Figure 6 and Table 2 show that the posterior density is clearly unimodal with respect to *θ*_1_ and *θ*_3_, the 95% CI are narrow and cover the true parameter values, mean and median of the sample are close or equal to the true values, and high ESSs and 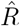 values equal to 1 are achieved. Thus, we can conclude that the parameters *θ*_1_ and *θ*_3_ are identifiable for the SDE model here.

**Figure 6:**
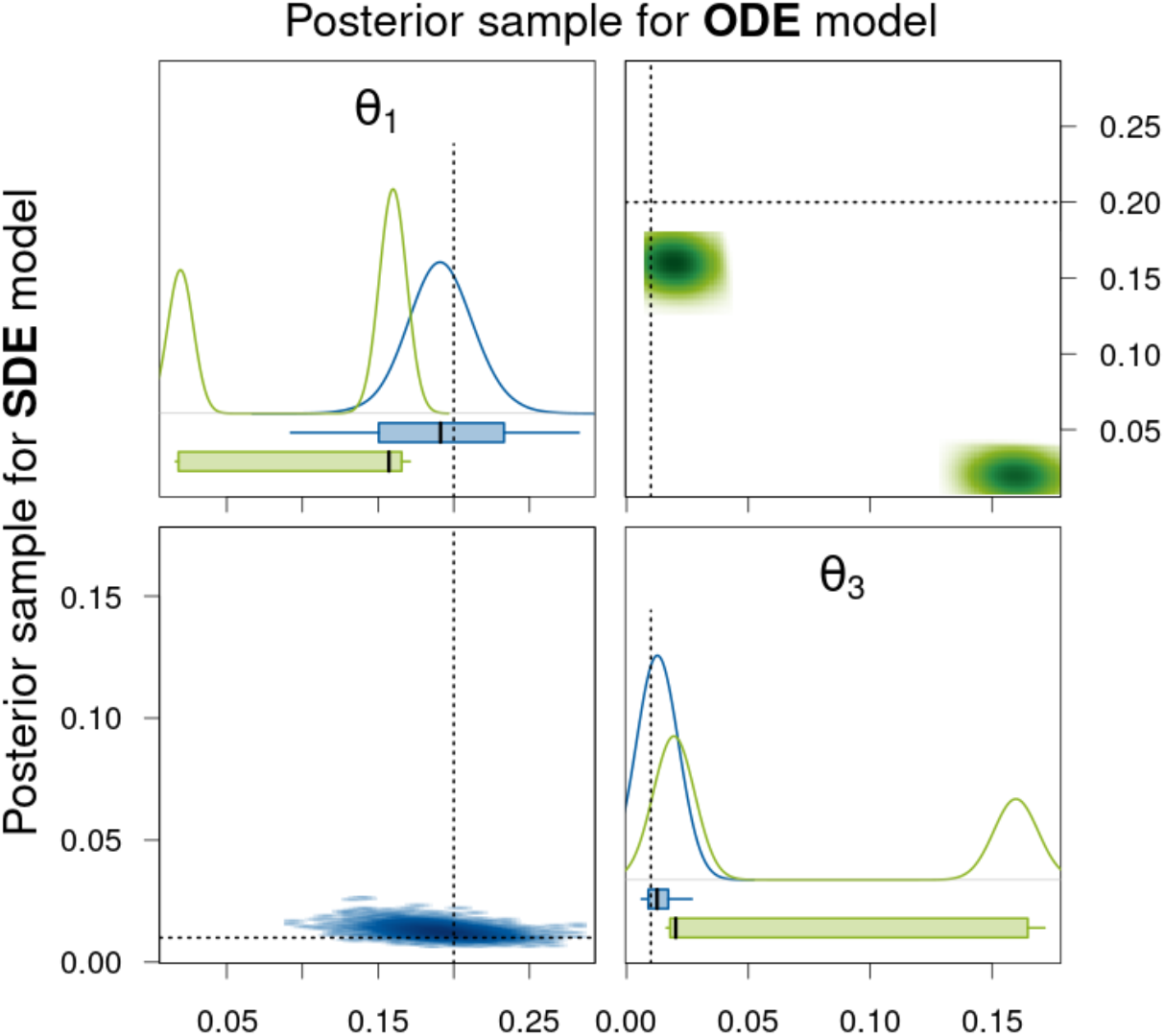
Density estimates of the posterior samples for parameters *θ*_1_ and *θ*_3_ for the SDE (blue, lower triangle) and ODE (green, upper triangle) model given simulated data without measurement error. *Diagonal panels:* Marginal densities for the respective parameter and boxplots showing the 95% CI as box, the range of the sample as whiskers, and the median as thick black line. *Off-diagonal panels:* Smoothed scatter plots of the two-dimensional projections of the samples where darker hues signify higher density values. The dotted lines represent the true parameter values that were used to simulate the data.

Next, we consider the translation rate constant *θ*_2_, the initial amount *m*_0_ of mRNA molecules, and the factor scale. For the ODE, at most the product *θ*_2_*m*_0_scale is identifiable. This is also apparent from the results presented in Table 1 and Figure 7. For the individual parameters and also for all products of two out of the three parameters, the 95% CIs are extremely broad and the mean and median as point estimates are not at all close to the true values. The reason why there are nevertheless quite high ESSs and 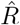 values equal to 1 achieved is that the variation within each of the HMC chains is very high and thus does not differ substantially from the variation between the chains for these parameters as can be seen in the traceplots of the chains in Section A.4.1 of the supplementary material. For the product *θ*_2_*m*_0_scale of all three parameters, the 95% CI is very narrow for the posterior sample of the ODE model and also the ESS is high and 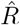 equal to 1. Without knowing the true parameter values, one would assume that this product is well estimated. However, the 95% CI and even the whole range of the sample do not cover the true value. For the SDE model, the 95% CI for the product *θ*_2_*m*_0_scale is broader, but it covers the true parameter value and also the mean and the median as point estimates are closer to the true value than the mean and the median for the ODE model. Moreover, the ESS is quite high and 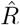 is equal to 1 for the SDE model. We therefore conclude that the product *θ*_2_*m*_0_scale is identifiable. The generally lower ESSs are due to the fact that for the SDE model, we sample from a distribution of much larger dimension as explained in Section 6.1. Additionally, for the SDE model, the parameters scale and *θ*_2_*m*_0_ have narrow 95% CIs (especially compared to those for the ODE model) that include the true parameter values, high ESSs, and 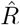 values of 1 and can therefore be considered identifiable. The remaining parameters *θ*_2_, *m*_0_, *θ*_2_scale, and *m*_0_scale have rather broad 95% CIs and only achieve low ESSs and 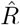 values higher than 1.02. Hence, they seem to be non-identifiable. Notice, however, that at least for the parameters *θ*_2_ and *θ*_2_scale, the 95% CIs are substantially more narrow for the SDE model compared to the ODE model.

**Figure 7:**
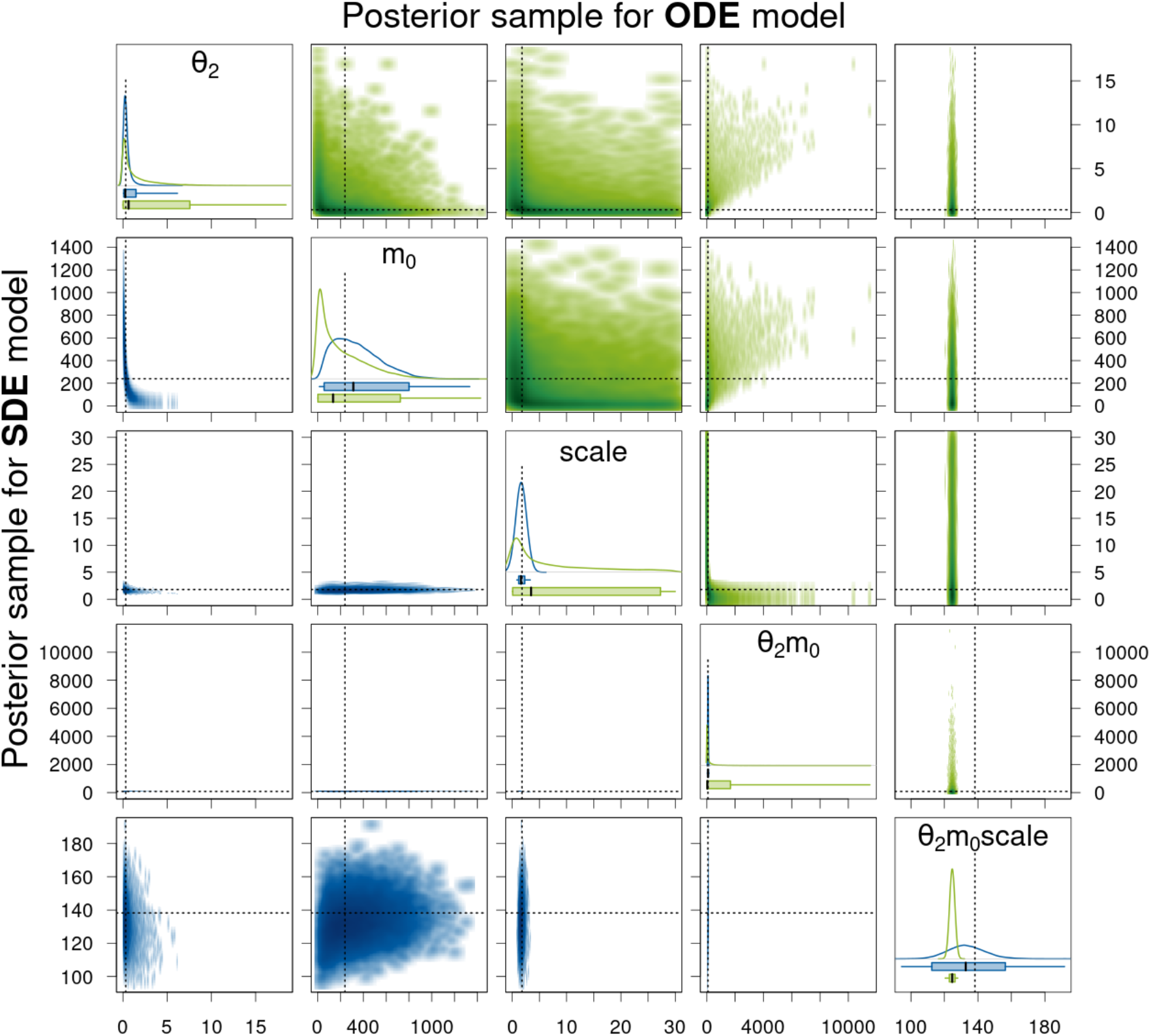
Density estimates of the posterior samples for parameters *θ*_2_, *m*_0_, scale, and their products for the SDE (blue, lower triangle) and ODE (green, upper triangle) model given simulated data without measurement error. For a detailed description of the figure’s elements, see Figure 6.

We have simulated another 99 trajectories with the same parameters and performed Stan sampling in the same way as described in the beginning of this subsection. For each model type and each posterior sample of the different simulated trajectories, we calculate the length of the 95% CI and determine the median and the coefficient of variation (c.v.) over these lengths for each model type. Also, we rescale the lengths of the 95% CI by dividing by the true parameter value and again determine the median of the normalized quantities. The rescaling is done to transfer the values to a more similar scale. Note, however, that the values are nevertheless not directly comparable between different parameters. Moreover, we check whether the true parameter value that was used to simulate the data is included in the 95% CI. Table 3 shows the aggregated results for the posterior samples of all 100 trajectories and also includes the length of the interval between the 2.5%- and the 97.5%-quantile of the prior distributions. Except for the parameters *m*_0_ and *θ*_2_*m*_0_scale, the median length of the 95% CIs for the SDE model is always smaller than for the ODE model. For parameter *θ*_2_*m*_0_scale, the CI lengths are a lot smaller for the ODE model; however, the CIs cover the true parameter value only 13 out of 100 times while for the SDE model, the true value is covered 93 times. For the other parameters that we classified as identifiable for the SDE model in the analysis of the individual trajectory (i. e. *θ*_1_, *θ*_3_, scale, and *θ*_2_*m*_0_), the median length of the 95% CIs is clearly smaller for the SDE model than for the ODE model and the true parameter value is covered at least 91 out of 100 times for the SDE model. For parameter *m*_0_, the CI lengths are high for both model types because the parameter is not identifiable for either model type. For the other parameters that we classified as not identifiable for both model types in the analysis of the individual trajectory (i.e. *θ*_2_, *θ*_2_scale, and *m*_0_scale), the median length of 95% CIs is clearly smaller for the SDE model than for the ODE model, at least by a factor of 4.

**Table 3:** Statistics of posterior samples for the two model types aggregated over 100 simulated trajectories without measurement error. We also include the length of the interval between the 2.5%- and the 97.5%-quantile of the prior distribution.

|  |  | length of<br>prior 95%<br>center<br>interval | median<br>length of<br>95% CIs | c.v. of<br>lengths of<br>95% CIs | median of<br>length of CIs<br>rescaled by<br>true value | number<br>of CIs<br>covering<br>true value |
| --- | --- | --- | --- | --- | --- | --- |
| $\theta_1$ | ODE | 11.05 | 0.20 | 0.009 | 1.01 | 58 |
|  | SDE | 11.05 | 0.09 | 0.002 | 0.45 | 93 |
| $\theta_2$ | ODE | 11.05 | 7.56 | 0.002 | 23.63 | 100 |
|  | SDE | 11.05 | 1.88 | 0.712 | 5.87 | 100 |
| $\theta_3$ | ODE | 11.05 | 0.20 | 0.010 | 20.00 | 61 |
|  | SDE | 11.05 | 0.01 | 0.000 | 0.80 | 91 |
| $m_0$ | ODE | 884.82 | 730.41 | 0.057 | 3.04 | 100 |
|  | SDE | 884.82 | 735.98 | 9.868 | 3.07 | 100 |
| scale | ODE | 28.50 | 27.27 | 0.001 | 15.15 | 100 |
|  | SDE | 28.50 | 1.20 | 0.158 | 0.67 | 95 |
| $\theta_2 m_0$ | ODE | 6056.48 | 1701.92 | 2.524 | 22.16 | 100 |
|  | SDE | 6056.48 | 63.60 | 2.768 | 0.83 | 96 |
| $\theta_2 \text{scale}$ | ODE | 228.08 | 55.47 | 0.164 | 96.29 | 100 |
|  | SDE | 228.08 | 2.89 | 0.762 | 5.01 | 100 |
| $m_0 \text{scale}$ | ODE | 19271.13 | 7923.38 | 8.603 | 18.34 | 100 |
|  | SDE | 19271.13 | 1315.63 | 86.806 | 3.05 | 99 |
| $\theta_2 m_0 \text{scale}$ | ODE | 113232.70 | 3.90 | 41.595 | 0.03 | 13 |
|  | SDE | 113232.70 | 45.28 | 0.624 | 0.33 | 93 |

The last two columns of Table 3 are visualized in Figure 8 where we plot the median of the rescaled CI lengths against the number of CIs that cover the true parameter value. The desirable region of value combinations is in the bottom right corner of the graph where the number of CIs covering the true value is high and the median rescaled CI length is small. Note that, clearly, more importance should be given to high numbers of CIs covering the true value as it is useless to be very certain about a parameter estimate (indicated by a short CI) while the correct value is not included in the CI. However, even for parameters that are identifiable, we do not expect to obtain a coverage of the true value of 100% since we are considering 95% CIs. Therefore, values of 100 rather tend to hint at non-identifiability. In Figure 8, we can see that for the majority of the parameters, the triangles representing the value combinations for the SDE model are closer to the desirable region. Only for parameter *m*_0_ (which is not identifiable for either model type), the value combinations are almost the same for both model types. And as we already pointed out for the product *θ*_2_*m*_0_scale, the median length of the 95% CIs is smaller for the ODE model; however, a lot fewer CIs cover the true parameter value for the ODE model than for the SDE model. Thus, the result obtained for the SDE model is to be preferred.

**Figure 8:**
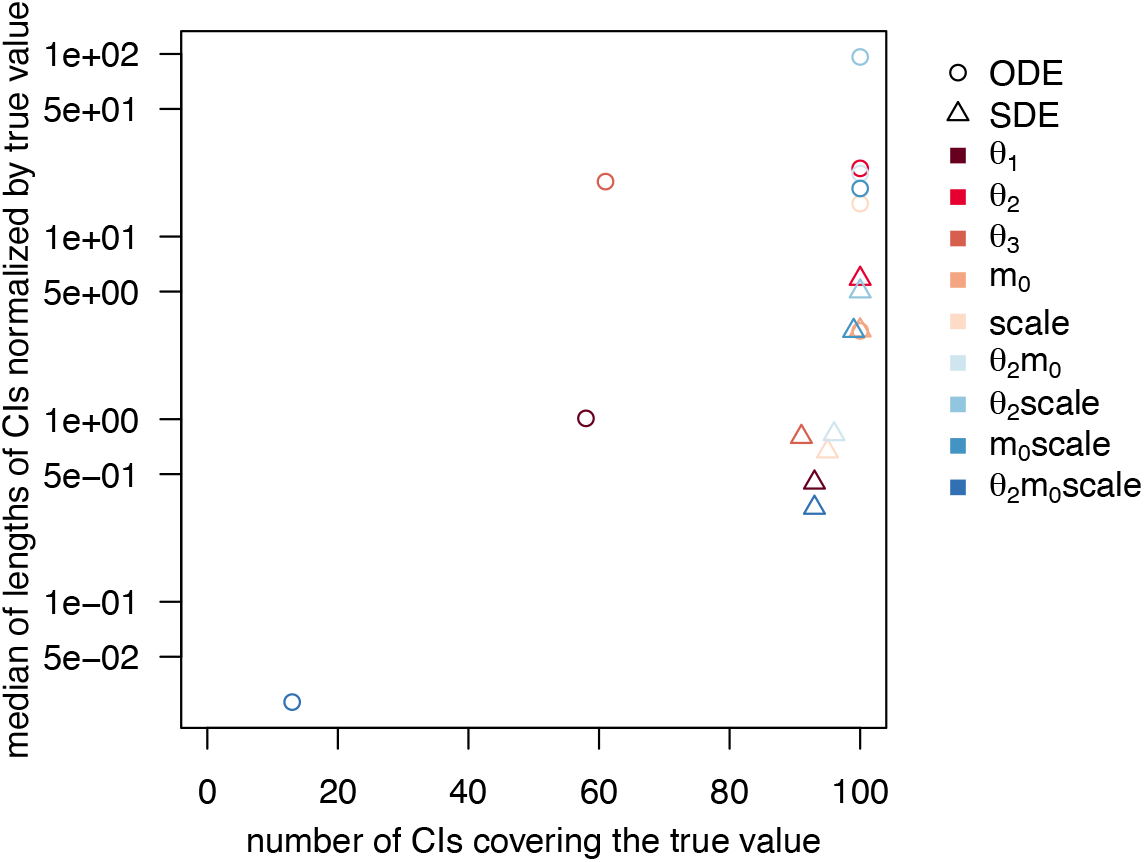
Statistics of posterior samples for the two model types aggregated over 100 simulated trajectories without measurement error. The desirable region of value combinations is in the bottom right corner of the graph.

We provide further Stan-specific diagnostics in Appendix A.5. Those mostly show poorer values for the sampling output for the SDE model than for the ODE model. This is not surprising as we sample from a much higher-dimensional distribution for the SDE model. We do not consider the poor diagnostics as a disadvantage of the procedure as they provide information that we do not even have for other MCMC algorithms and thus cannot compare to them.

#### 6.2.2 Sampling results for simulated data with measurement error

In this section, we use the same simulated data as in the previous section, but for each of the 100 trajectories, we add multiplicative measurement error with parameter *σ* = 0.02. Again, we use Stan to sample from the posterior distributions of the ODE model (6) and the SDE model (7) for each of the simulated trajectories and use the same priors as stated in the previous section. We generate 8 HMC chains of 5000 iterations, discard the first half of the iterations as warm-up, and thus use a posterior samples of size 20,000 in the subsequent analysis.

At first, we again focus on the results for one of the trajectories, namely the trajectory represented by the green dotted line in Figure 5. Tables 4 and 5 summarize the Stan output of the posterior samples for the ODE and the SDE model, respectively. The parameter *t*_0_ is estimated very accurately based on the posterior sample for the ODE model. Also, the parameter offset is well estimated for both model types but with a more narrow 95% CI for the SDE model. The parameter *σ* is accurately determined for the SDE model as well. For the ODE model, *σ* is again overestimated. Figure 9 visualizes the components of the posterior samples for parameters *θ*_1_, *θ*_3_, offset, and *σ*. Again, the bimodality of the posterior with respect to *θ*_1_ and *θ*_3_ is apparent for the ODE model and neither the 95% CIs nor the ranges of the sample cover the true parameter values. For the SDE model on the other hand, the distribution is unimodal and the 95% CIs do cover the true parameter values for *θ*_1_ and *θ*_3_. However, their 2-dimensional smoothed scatter plot in Figure 9 does not show a simple elliptic shape (as for the simulated data without measurement error) but almost a banana-like shape. This may also be the reason for the deteriorated sampling efficiency discernible from the low ESSs and higher 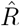-values in Table 5.

**Table 4:** Summary of the Stan output for the ODE model given simulated data with measurement error and the true parameter values that were used to simulate the data. c.v. denotes the coefficient of variation and the columns headed by percentages contain the quantiles of the respective percentage value.

| | true value | mean | c.v. | 2.5% | 50% | 97.5% | $n_{\text{eff}}$ | $\hat{R}$ |
| --- | --- | --- | --- | --- | --- | --- | --- | --- |
| $\theta_1$ | 0.20 | 0.11 | 0.632 | 0.02 | 0.15 | 0.17 | 4 | 20.94 |
| $\theta_2$ | 0.32 | 1.54 | 1.364 | 0.02 | 0.65 | 7.63 | 11619 | 1.00 |
| $\theta_3$ | 0.01 | 0.07 | 0.938 | 0.02 | 0.02 | 0.16 | 4 | 26.81 |
| $m_0$ | 240.00 | 205.41 | 1.024 | 2.25 | 133.96 | 738.49 | 10649 | 1.00 |
| scale | 1.80 | 6.84 | 1.152 | 0.07 | 3.29 | 27.06 | 8778 | 1.00 |
| offset | 6.50 | 6.50 | 0.013 | 6.34 | 6.50 | 6.67 | 14496 | 1.00 |
| $t_0$ | 0.96 | 0.96 | 0.003 | 0.96 | 0.96 | 0.97 | 17129 | 1.00 |
| $\sigma$ | 0.02 | 0.03 | 0.053 | 0.03 | 0.03 | 0.04 | 13581 | 1.00 |
| $\theta_2 m_0$ | 76.80 | 217.59 | 2.441 | 4.56 | 37.40 | 1683.06 | 9479 | 1.00 |
| $\theta_2 \text{scale}$ | 0.58 | 6.31 | 2.841 | 0.17 | 0.92 | 54.94 | 7209 | 1.00 |
| $m_0 \text{scale}$ | 432.00 | 983.07 | 2.151 | 16.20 | 189.75 | 7514.28 | 8054 | 1.00 |
| $\theta_2 m_0 \text{scale}$ | 138.24 | 123.47 | 0.009 | 121.43 | 123.46 | 125.58 | 12035 | 1.00 |

**Table 5:**
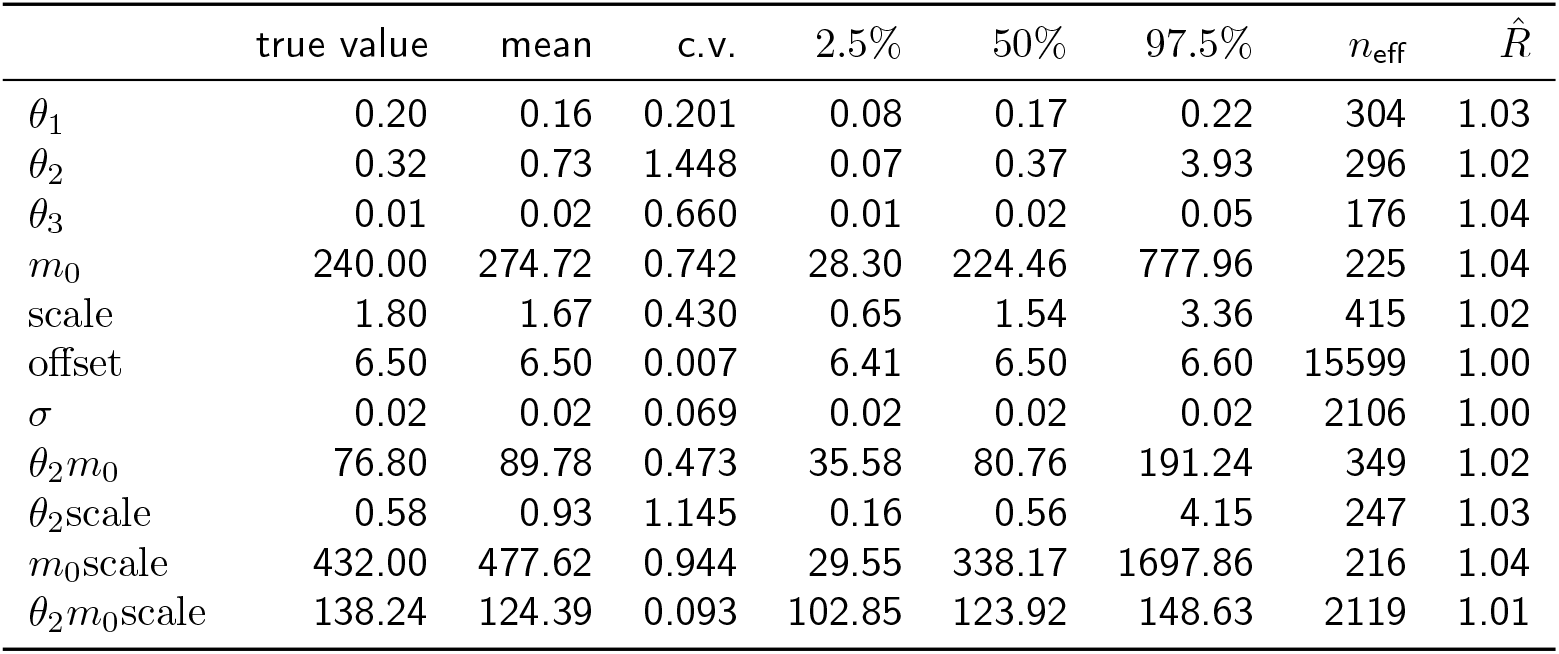
Summary of the Stan output for the SDE model given simulated data with measurement error and the true parameter values that were used to simulate the data. The initial time point *t*_0_ is not estimated here, but predetermined based on the mean estimate of the sample for the ODE model.

| | true value | mean | c.v. | 2.5% | 50% | 97.5% | $n_{\text{eff}}$ | $\hat{R}$ |
| --- | --- | --- | --- | --- | --- | --- | --- | --- |
| $\theta_1$ | 0.20 | 0.16 | 0.201 | 0.08 | 0.17 | 0.22 | 304 | 1.03 |
| $\theta_2$ | 0.32 | 0.73 | 1.448 | 0.07 | 0.37 | 3.93 | 296 | 1.02 |
| $\theta_3$ | 0.01 | 0.02 | 0.660 | 0.01 | 0.02 | 0.05 | 176 | 1.04 |
| $m_0$ | 240.00 | 274.72 | 0.742 | 28.30 | 224.46 | 777.96 | 225 | 1.04 |
| scale | 1.80 | 1.67 | 0.430 | 0.65 | 1.54 | 3.36 | 415 | 1.02 |
| offset | 6.50 | 6.50 | 0.007 | 6.41 | 6.50 | 6.60 | 15599 | 1.00 |
| $\sigma$ | 0.02 | 0.02 | 0.069 | 0.02 | 0.02 | 0.02 | 2106 | 1.00 |
| $\theta_2 m_0$ | 76.80 | 89.78 | 0.473 | 35.58 | 80.76 | 191.24 | 349 | 1.02 |
| $\theta_2 \text{scale}$ | 0.58 | 0.93 | 1.145 | 0.16 | 0.56 | 4.15 | 247 | 1.03 |
| $m_0 \text{scale}$ | 432.00 | 477.62 | 0.944 | 29.55 | 338.17 | 1697.86 | 216 | 1.04 |
| $\theta_2 m_0 \text{scale}$ | 138.24 | 124.39 | 0.093 | 102.85 | 123.92 | 148.63 | 2119 | 1.01 |

**Figure 9:**
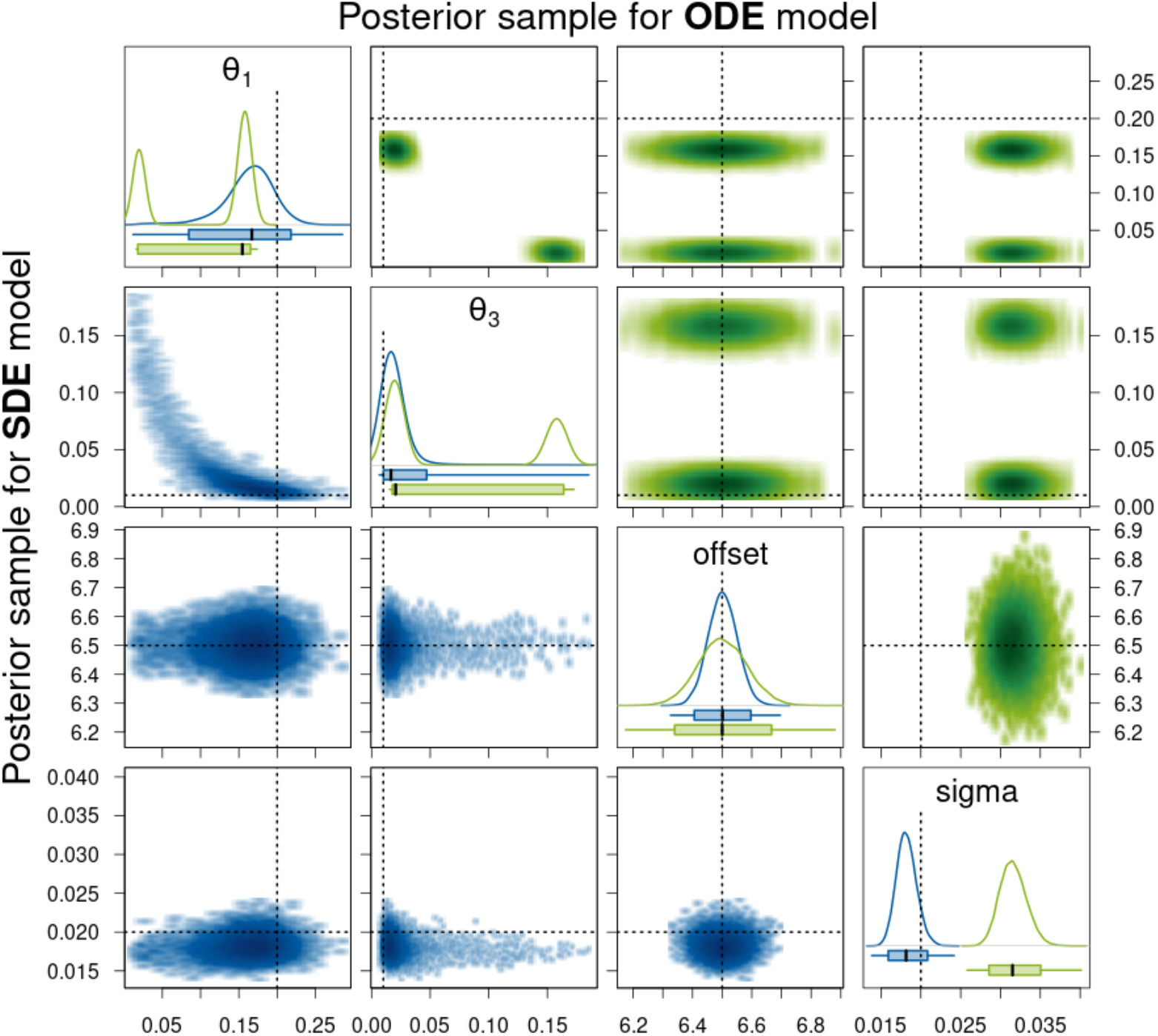
Density estimates of the posterior samples for parameters *θ*_1_, *θ*_3_, offset, and *σ* for the SDE (blue, lower triangle) and ODE (green, upper triangle) model given simulated data with measurement error. For a detailed description of the figure’s elements, see Figure 6.

Figure 9 visualizes the components of the posterior samples for parameters *θ*_2_, *m*_0_, scale, and their products. For the ODE model, again only the product *θ*_2_*m*_0_scale is identifiable in the sense that the corresponding 95% CI is very narrow, the ESS is high, and the 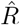-value is equal to 1. However, the 95% CI again does not cover the true parameter value. For the SDE model, the 95% CI for *θ*_2_*m*_0_scale is broader but it does contain the true value. Also, the ESS is high and the 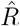-value is close to 1. Moreover, the parameters scale and *θ*_2_*m*_0_ have narrow 95% CIs, high ESSs, and 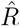-values close to 1 for the SDE model, and thus, we conclude that they are identifiable. Note that also for *θ*_2_, *m*_0_scale, and *θ*_2_scale, the 95% CIs are much narrower for the SDE model than for the ODE model.

In Section A.4.1 of the supplementary material, we also include trace plots and further figures of the sampling output for the trajectory displayed in Figure 5. There, we present the same posterior samples as used in this and the previous subsection. But instead of comparing the posterior samples between the two model types, the posterior samples are compared between the simulated data without and with measurement error for each model type separately. In summary, we find that for the SDE model, the 95% CIs increase for almost all parameters except *m*_0_ for data with measurement error. Whereas for the ODE model, there is hardly any difference for most of the parameters between the posterior samples for the data without and with measurement error since the majority of the parameters is not identifiable anyway. The marginal posterior samples for the parameters offset, *t*_0_, and *θ*_2_*m*_0_scale are only slightly affected by the measurement error. Only the marginal posterior sample of the measurement error parameter *σ* is substantially affected and, as expected, consists of higher values for data with measurement error.

Table 6 and Figure 11 display the statistics of the posterior samples aggregated over the 100 simulated trajectories. Similar to the results for the simulated data without measurement error, the median length of the 95% CIs for the SDE model is always smaller than for the ODE model, except for the parameters *m*_0_ and *θ*_2_*m*_0_scale and additionally *σ* (which was not included for the SDE model in the previous subsection). Again, for the majority of the parameters, the results for the SDE model represented by triangles in Figure 11 are closer to the desirable region of value combinations in the bottom right corner of the graph, except for the parameters *m*_0_, *θ*_2_*m*_0_scale, and offset. For the parameter offset, the median CI length is slightly higher for the ODE model than for the SDE model, however, the CIs for the ODE model also contain the true parameter value more often. So for this parameter, the ODE model, for once, shows the preferable result.

**Table 6:** Statistics of posterior samples for the two model types aggregated over 100 simulated trajectories with measurement error. We also include the length of the interval between the 2.5%- and the 97.5%-quantile of the prior distribution.

|  |  | length of<br>prior 95%<br>center<br>interval | median<br>length of<br>95% CIs | c.v. of<br>lengths of<br>95% CIs | median of<br>length of CIs<br>rescaled by<br>true value | number<br>of CIs<br>covering<br>true value |
| --- | --- | --- | --- | --- | --- | --- |
| $\theta_1$ | ODE | 11.05 | 0.20 | 0.008 | 1.01 | 60 |
|  | SDE | 11.05 | 0.11 | 0.016 | 0.55 | 90 |
| $\theta_2$ | ODE | 11.05 | 7.55 | 0.002 | 23.59 | 100 |
|  | SDE | 11.05 | 3.69 | 0.996 | 11.52 | 99 |
| $\theta_3$ | ODE | 11.05 | 0.20 | 0.008 | 20.35 | 63 |
|  | SDE | 11.05 | 0.02 | 0.094 | 1.65 | 89 |
| $m_0$ | ODE | 884.82 | 733.85 | 3.584 | 3.06 | 100 |
|  | SDE | 884.82 | 746.74 | 5.909 | 3.11 | 100 |
| scale | ODE | 28.50 | 27.25 | 0.002 | 15.14 | 100 |
|  | SDE | 28.50 | 2.54 | 0.255 | 1.41 | 91 |
| $\theta_2 m_0$ | ODE | 6056.48 | 1702.37 | 2.714 | 22.17 | 100 |
|  | SDE | 6056.48 | 223.22 | 428.350 | 2.91 | 92 |
| $\theta_2 \text{scale}$ | ODE | 228.08 | 55.70 | 0.125 | 96.70 | 100 |
|  | SDE | 228.08 | 3.55 | 0.639 | 6.16 | 100 |
| $m_0 \text{scale}$ | ODE | 19271.13 | 7896.07 | 667.494 | 18.28 | 100 |
|  | SDE | 19271.13 | 1479.13 | 253.530 | 3.42 | 98 |
| $\theta_2 m_0 \text{scale}$ | ODE | 113232.70 | 4.96 | 38.038 | 0.04 | 15 |
|  | SDE | 113232.70 | 45.56 | 1.492 | 0.33 | 92 |
| offset | ODE | 28.50 | 0.33 | 0.987 | 0.05 | 96 |
|  | SDE | 28.50 | 0.21 | 0.016 | 0.03 | 84 |
| $\sigma$ | ODE | 9.50 | 0.01 | 0.325 | 0.35 | 0 |
|  | SDE | 9.50 | 0.01 | 0.000 | 0.25 | 87 |

**Figure 10:**
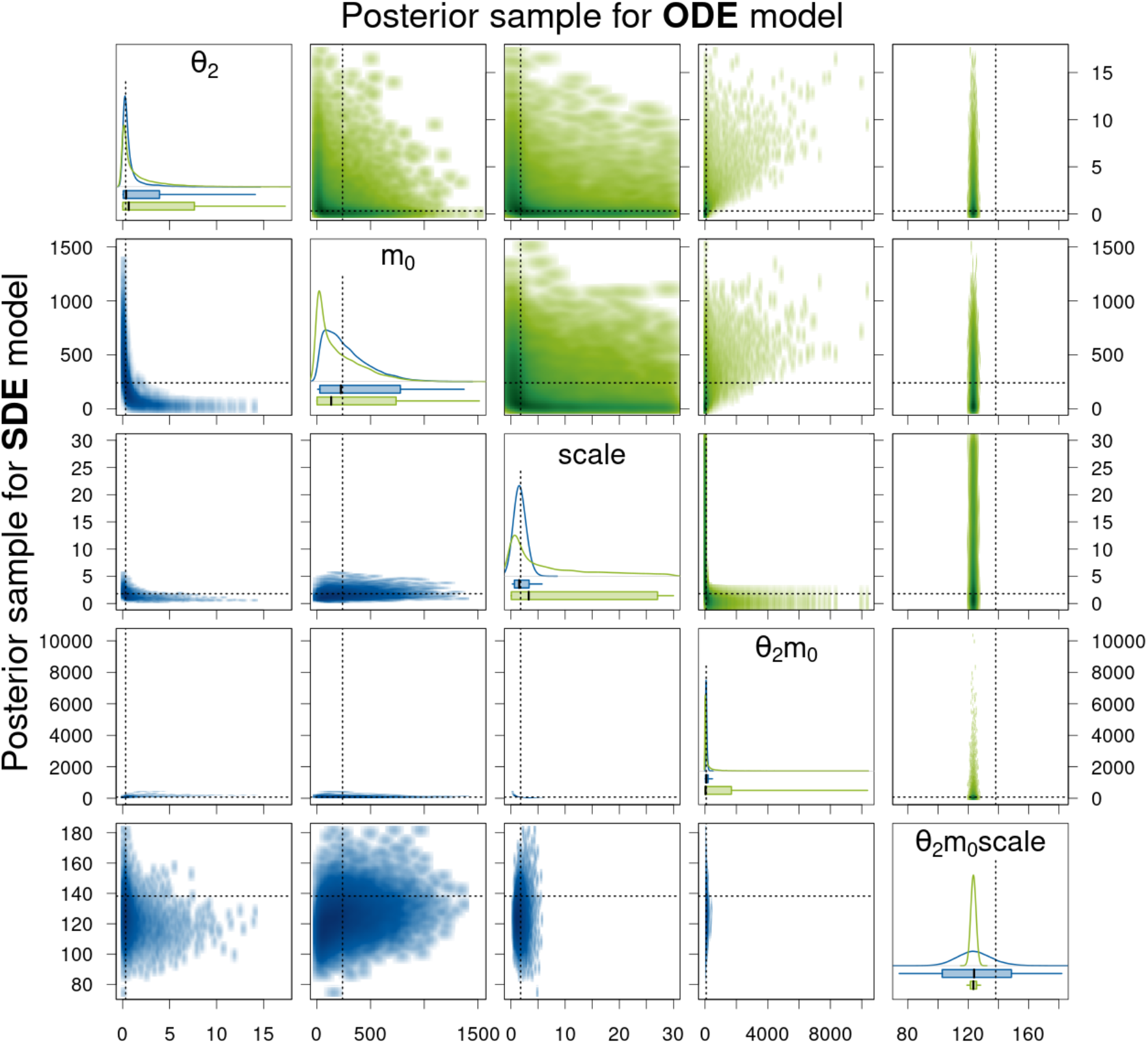
Density estimates of the posterior samples for parameters *θ*_2_, *m*_0_, scale, and their products for the SDE (blue, lower triangle) and ODE (green, upper triangle) model given simulated data with measurement error. For a detailed description of the figure’s elements, see Figure 6.

**Figure 11:**
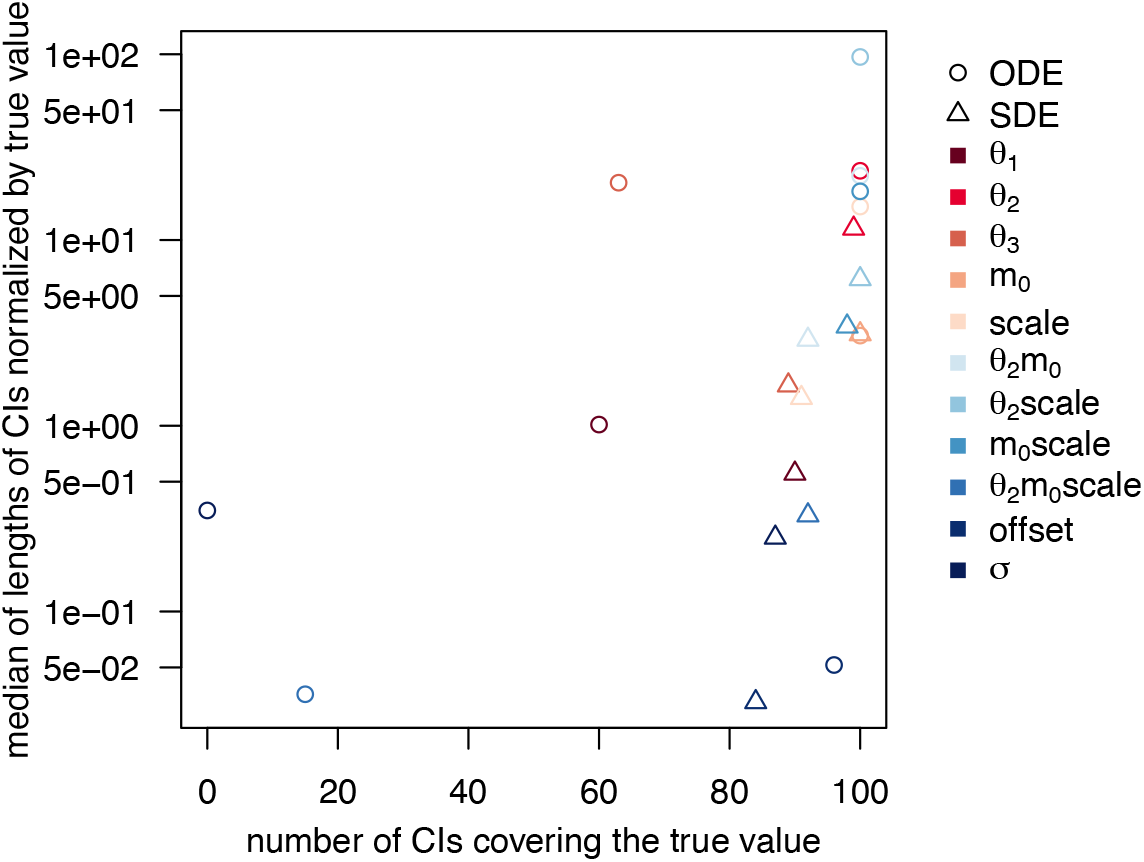
Statistics of posterior samples for the two model types aggregated over 100 simulated trajectories with measurement error. The desirable region of value combinations is in the bottom right corner of the graph.

### 6.3 Estimation based on experimental data

Since our results in the previous section have shown that the SDE model yields more reliable parameter estimates (assuming an MJP is an adequate description for the translation kinetics after mRNA transfection) than the ODE model even for those parameters that are identifiable for both model types, we reanalyze the experimental data published in Fröhlich et al. (2018) and described in Section 2. For each type of GFP (eGFP and d2eGFP), we randomly select 100 observed trajectories for our analysis and again use Stan to sample from the posterior distributions of the ODE model (6) and the SDE model (7) for each of the trajectories using the same priors as stated in Section 6.2.1. We generate 8 HMC chains of 5000 iterations, discard the first half of the iterations as warm-up, and thus use a posterior samples of size 20,000 in the subsequent analysis. For each type of GFP, we first analyze the sampling output for one observed trajectory in detail and then summarize results for all 100 observed trajectories. Here in the main part of the article, we only show the results for the eGFP data and provide the results for the d2eGFP data in Appendix A.4.3. Moreover, we provide further Stan-specific diagnostics in Appendix A.5.

#### Sampling results for experimental dataset 1 (for eGFP)

Tables 7 and 8 present a summary of the Stan output for the posterior sample of one observed trajectory for the ODE and the SDE model, respectively, and Figures 12 and 13 compare the density estimates of these two posterior samples. Moreover, we provide trace plots for these two posterior samples in Section A.4.2 in the supplementary material. The results look qualitatively *very* similar (almost identical) to those obtained for the simulated data with measurement error in Section 6.2.2. Therefore, we do not repeat the detailed description but only point out that the range of values sampled for the parameters *θ*_1_ and *θ*_3_ for the SDE model is slightly smaller for the experimental trajectory here. Thus, we do not see the banana-like shape in the two-dimensional smoothed scatter plot of the two parameters for the SDE model in Figure 12 as for the simulated trajectory in Figure 9 and the sampling efficiency increases as indicated by higher ESSs and lower 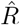 values for the two parameters in Table 8.

**Table 7:**
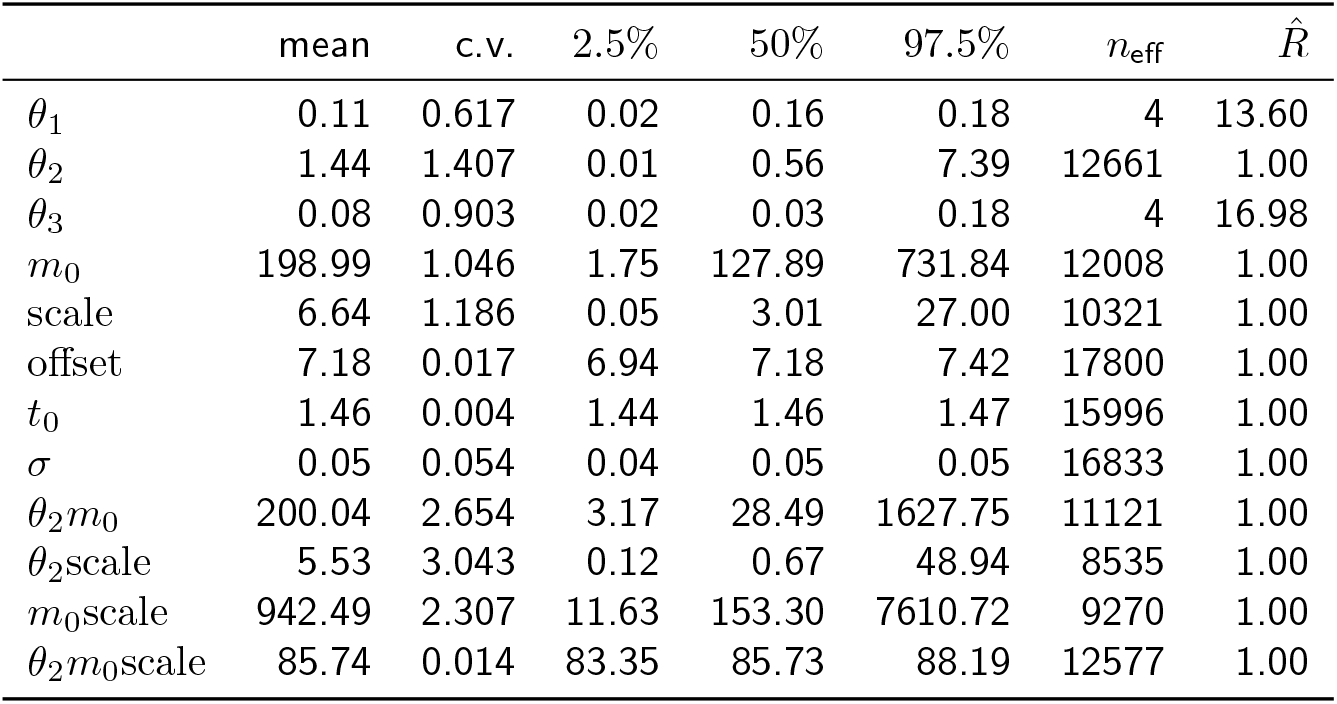
Summary of the Stan output for the ODE model given experimental data for eGFP. c.v. denotes the coefficient of variation and the columns headed by percentages contain the quantiles of the respective percentage value.

| | mean | c.v. | 2.5% | 50% | 97.5% | $n_{\text{eff}}$ | $\hat{R}$ |
| --- | --- | --- | --- | --- | --- | --- | --- |
| $\theta_1$ | 0.11 | 0.617 | 0.02 | 0.16 | 0.18 | 4 | 13.60 |
| $\theta_2$ | 1.44 | 1.407 | 0.01 | 0.56 | 7.39 | 12661 | 1.00 |
| $\theta_3$ | 0.08 | 0.903 | 0.02 | 0.03 | 0.18 | 4 | 16.98 |
| $m_0$ | 198.99 | 1.046 | 1.75 | 127.89 | 731.84 | 12008 | 1.00 |
| scale | 6.64 | 1.186 | 0.05 | 3.01 | 27.00 | 10321 | 1.00 |
| offset | 7.18 | 0.017 | 6.94 | 7.18 | 7.42 | 17800 | 1.00 |
| $t_0$ | 1.46 | 0.004 | 1.44 | 1.46 | 1.47 | 15996 | 1.00 |
| $\sigma$ | 0.05 | 0.054 | 0.04 | 0.05 | 0.05 | 16833 | 1.00 |
| $\theta_2 m_0$ | 200.04 | 2.654 | 3.17 | 28.49 | 1627.75 | 11121 | 1.00 |
| $\theta_2 \text{scale}$ | 5.53 | 3.043 | 0.12 | 0.67 | 48.94 | 8535 | 1.00 |
| $m_0 \text{scale}$ | 942.49 | 2.307 | 11.63 | 153.30 | 7610.72 | 9270 | 1.00 |
| $\theta_2 m_0 \text{scale}$ | 85.74 | 0.014 | 83.35 | 85.73 | 88.19 | 12577 | 1.00 |

**Table 8:** Summary of the Stan output for the SDE model given experimental data for eGFP. The initial time point *t*_0_ is not estimated here, but predetermined based on the mean estimate of the sample for the ODE model.

| | mean | c.v. | 2.5% | 50% | 97.5% | $n_{\text{eff}}$ | $\hat{R}$ |
| --- | --- | --- | --- | --- | --- | --- | --- |
| $\theta_1$ | 0.20 | 0.152 | 0.14 | 0.20 | 0.26 | 946 | 1.01 |
| $\theta_2$ | 0.33 | 1.366 | 0.04 | 0.19 | 1.52 | 305 | 1.02 |
| $\theta_3$ | 0.02 | 0.269 | 0.01 | 0.02 | 0.03 | 894 | 1.01 |
| $m_0$ | 298.35 | 0.705 | 34.46 | 250.62 | 809.42 | 218 | 1.03 |
| scale | 2.12 | 0.309 | 1.14 | 2.02 | 3.69 | 772 | 1.01 |
| offset | 7.18 | 0.012 | 7.01 | 7.18 | 7.35 | 20589 | 1.00 |
| $\sigma$ | 0.03 | 0.058 | 0.03 | 0.03 | 0.04 | 17224 | 1.00 |
| $\theta_2 m_0$ | 47.92 | 0.327 | 23.41 | 46.00 | 83.31 | 857 | 1.01 |
| $\theta_2 \text{scale}$ | 0.60 | 1.182 | 0.11 | 0.37 | 2.54 | 256 | 1.02 |
| $m_0 \text{scale}$ | 650.64 | 0.831 | 57.98 | 498.32 | 2049.93 | 283 | 1.02 |
| $\theta_2 m_0 \text{scale}$ | 92.71 | 0.115 | 73.32 | 92.15 | 115.16 | 2711 | 1.00 |

**Figure 12:**
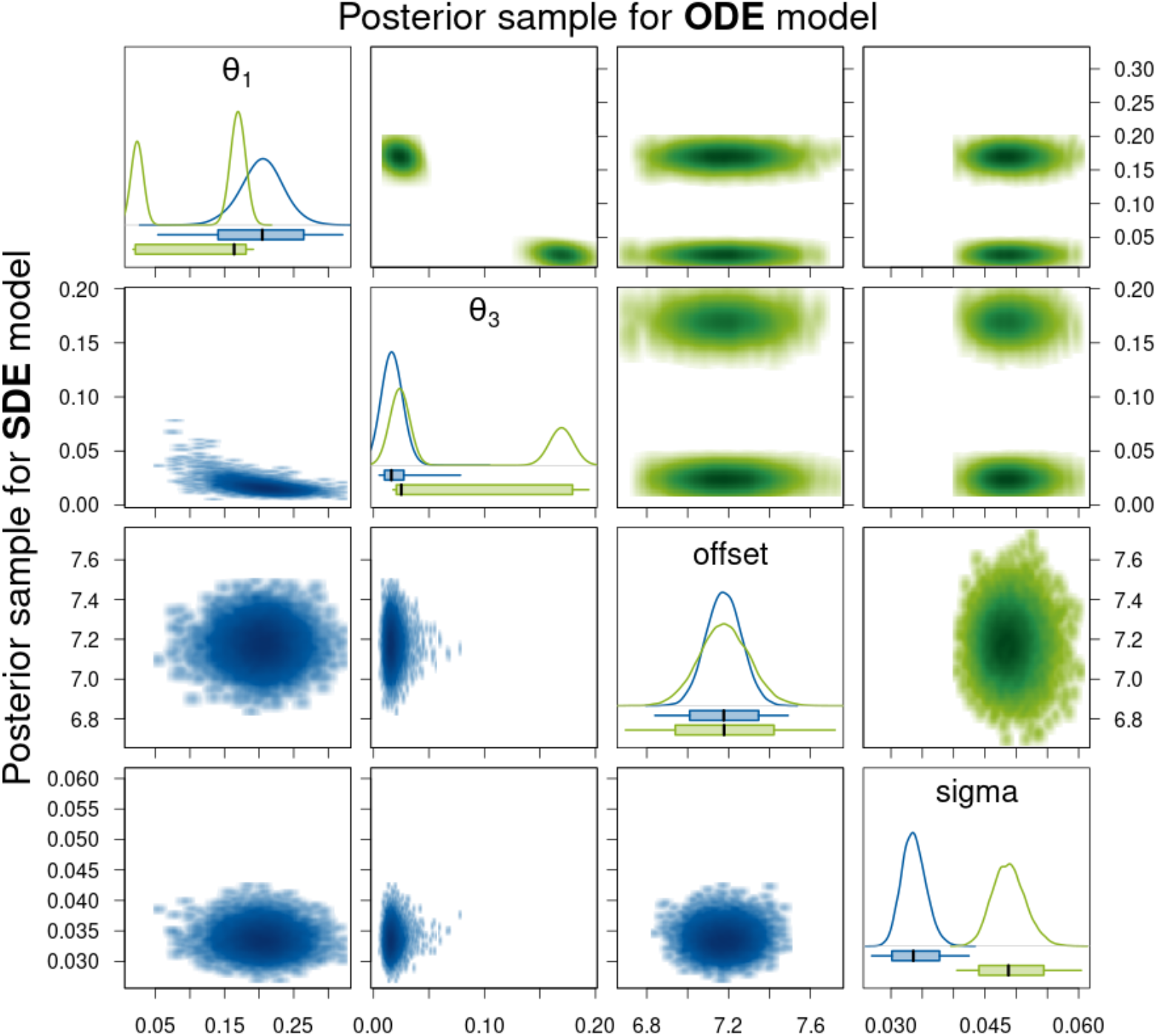
Density estimates of the posterior samples for parameters *θ*_1_, *θ*_3_, offset, and *σ* for the SDE (blue, lower triangle) and ODE (green, upper triangle) model given experimental data for eGFP. *Diagonal panels:* Marginal densities for the respective parameter and boxplots showing the 95% CI as box, the range of the sample as whiskers, and the median as thick black line. *Off-diagonal panels:* Smoothed scatter plots of the two-dimensional projections of the samples where darker hues signify higher density values.

**Figure 13:**
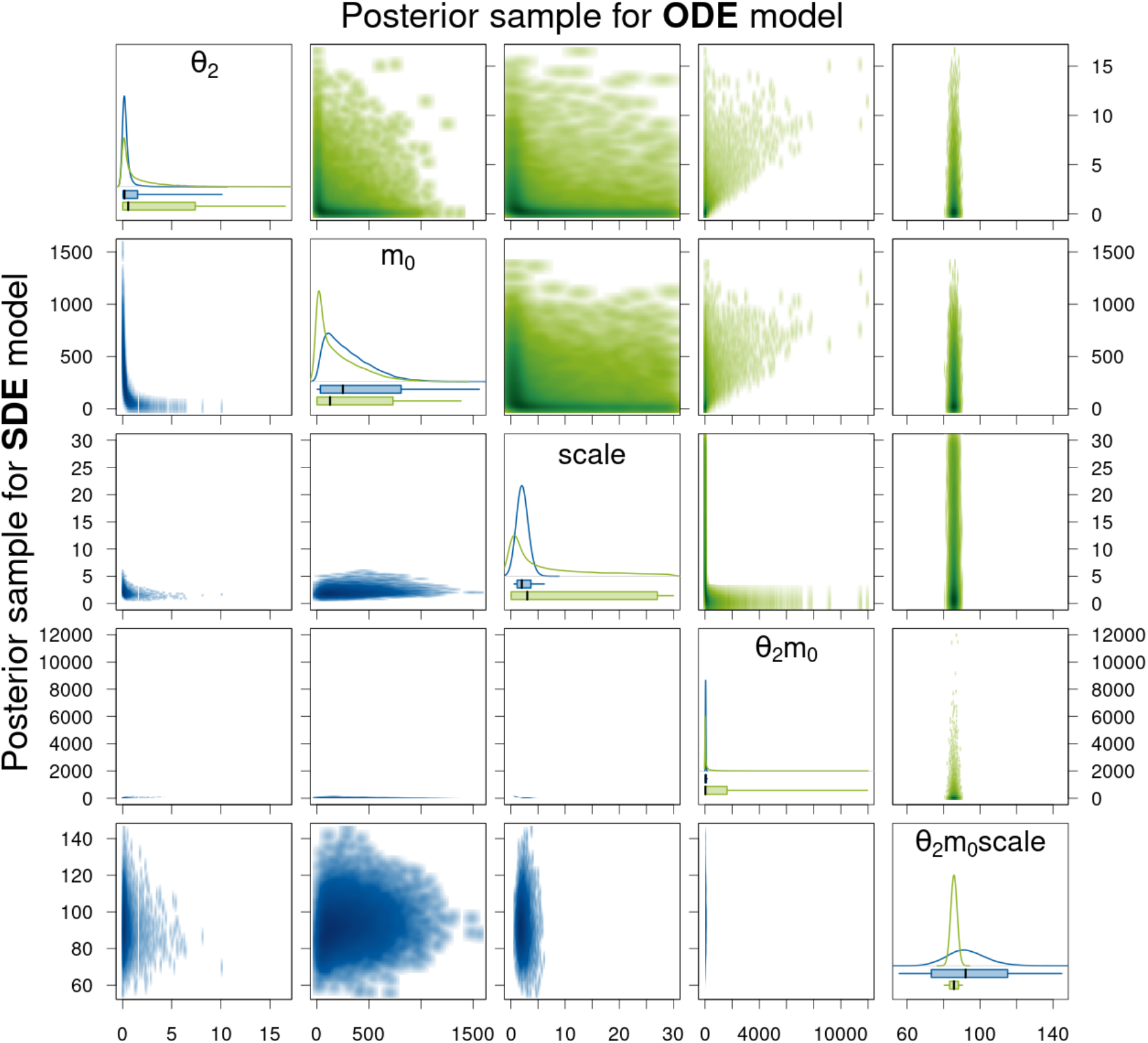
Density estimates of the posterior samples for parameters *θ*_2_, *m*_0_, scale, and their products for the SDE (blue, lower triangle) and ODE (green, upper triangle) model given experimental data for eGFP. For a detailed description of the figure’s elements, see Figure 12.

The statistics of posterior samples aggregated for 100 experimental trajectories for eGFP in Table 9 are also qualitatively similar to those for the simulated trajectories in Table 6. For the majority of the parameters, the median length of the 95% CI is smaller for the posterior samples for the SDE model than for those for the ODE model. Only for parameters *θ*_1_, *θ*_2_, and *θ*_2_*m*_0_scale, this is not the case. Note in particular that for the parameters *θ*_2_*m*_0_ and scale, which are non-identifiable for the ODE model (also apparent from the very long CIs here), the median length of the 95% CI for the SDE model is again much smaller compared to that of the ODE and to that of the prior. This indicates that these two parameters are identifiable for the SDE model also for the experimental data. That the uncertainty of the parameter estimate for *θ*_2_*m*_0_scale is greater for the SDE than for the ODE model is consistent with our results for the simulated data. The parameter *θ*_2_ is considered to be non-identifiable for both model types and the difference between the median CI lengths is relatively small. Finally, for parameter *θ*_1_, we see that the result is more or less the same as for *θ*_3_ for the ODE model due to the symmetry of the posterior distribution with respect to these two parameters. Whereas for the SDE model there is no symmetry and there is more variance in the posterior samples with respect to *θ*_1_ than to *θ*_3_ (indicated by a greater median CI length). The smaller median CI length of *θ*_1_ for the ODE model compared to the SDE model is due to the fact that for many of the observed trajectories the values of *θ*_1_ and *θ*_3_ seem to be very close together. In this case, the posterior distribution of the ODE model appears to be unimodal (as the two modes overlap) and the posterior variance with respect to the two parameters is small (and equal due to the symmetry). Thus, overall this variance is smaller than the posterior variance with respect to *θ*_1_ for the SDE model.

**Table 9:**
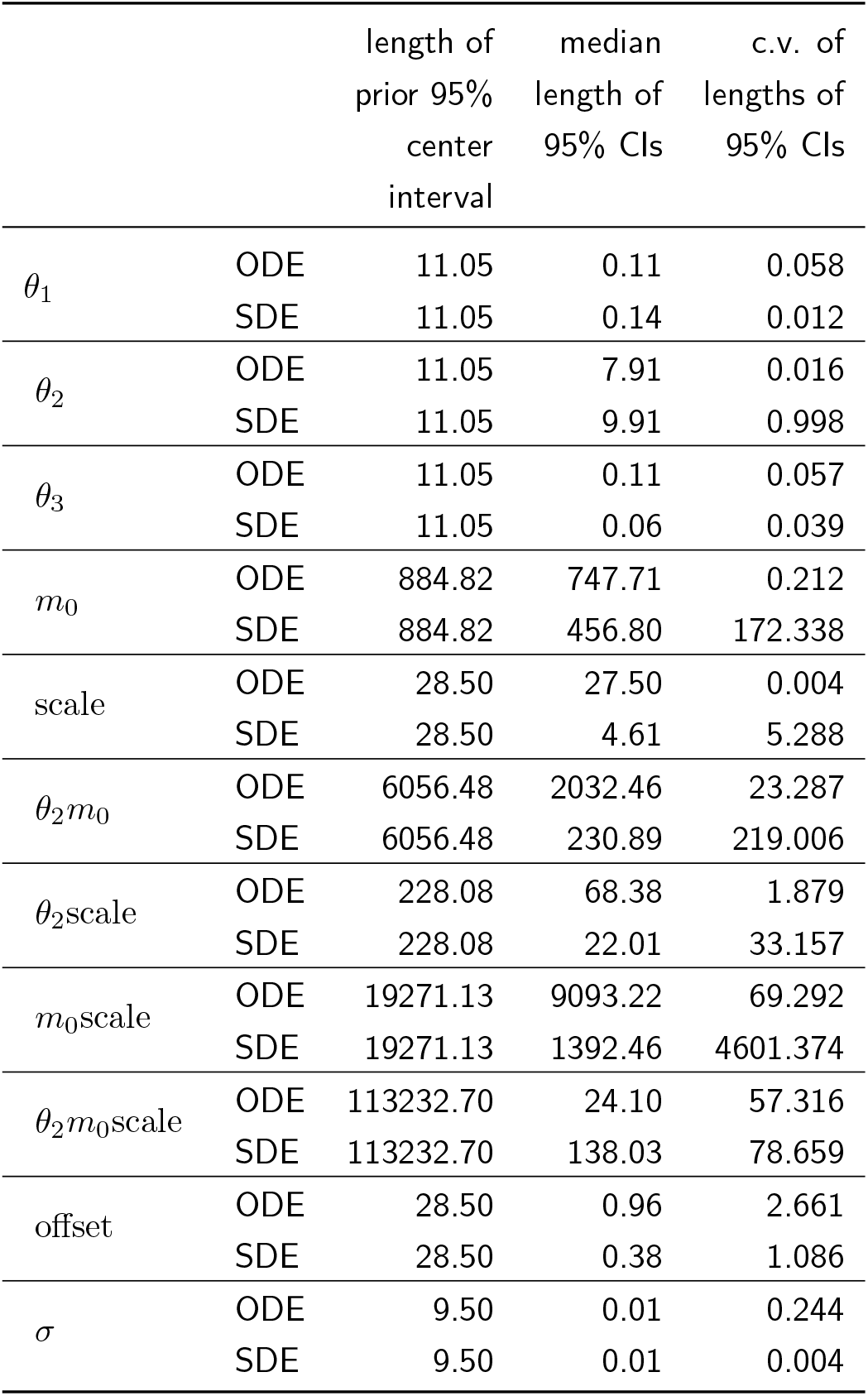
Statistics of posterior samples aggregated for 100 experimental trajectories for eGFP.

## 7 Discussion and conclusion

We have modeled the translation kinetics after mRNA transfection using a two-dimensional Itô diffusion process described by an SDE and compared this modeling approach to one using ODEs. We have studied the parameter identifiability for both modeling approaches for the case that we observe a fluorescence signal which we assume to be a linear transformation of the amount of protein molecules (corrupted by multiplicative measurement error). Our analysis using the structural identifiability of the moment-equations-based surrogate model and a simulation based assessment suggest that the SDE model might lead to better parameter identifiability. The most systematic approach is the one based on the surrogate model and DAISY as suggested by Browning et al. (2020); however, it cannot help us *confirm* a difference in the parameter identifiability between the SDE and the ODE model. Especially because we are interested in the parameter identifiability based on one observed trajectory and the moment-equations-based approach assumes that we were able to observe the first and the second moment of the fluorescence signal. Even when we take into account that we have several observed trajectories available from the experiment, these do not provide information about the moments because the initial time point *t*_0_ of mRNA release is different for every trajectory and also for the other parameters, in particular for *m*_0_, assuming that they are equal for all observed cells does not seem reasonable. By simulating from the SDE model, we were able to assess the differences in the variation within individual trajectories for different parameter combinations. Again our results suggest that the SDE model provides better parameter identifiability. Unlike for the ODE model, the degradation rates *θ*_1_ and *θ*_3_ for the SDE model appear to be structurally globally identifiable. Also, scale and the product *θ*_2_*m*_0_ seem to be structurally identifiable, but the individual parameters *θ*_2_ and *m*_0_ do not. While this simple simulation approach worked out well for the model considered here, one of its weak points is, of course, the somewhat subjective visual assessment of the variation within trajectories. A more quantitative approach to this would be to simulate a large number of trajectories (with very small time step) for every considered parameter combination, to approximate the quadratic variation for each trajectory, and then, to compare these values between individual trajectories started with the same seed for different parameter combinations and to compare also the distributions of these values for different parameter combinations. Another drawback of both simulation-based approaches is the fact that the analysis is based on a finite set of parameter combinations that can be considered; and thus, drawing general conclusions for the entire parameter space may be problematic.

Moreover, we have assessed the practical parameter identifiability for both model types by sampling from the parameter posterior distribution given simulated data with and without measurement error and the experimental data published in Fröhlich et al. (2018). While our focus was on inference from individual fluorescence trajectories, using SDE mixed-effect modeling as done in Wiqvist et al. (2021) would be a meaningful extension, especially e. g. to obtain a common estimate for the parameter scale. We found that the parameters *θ*_1_ and *θ*_3_ are indeed globally identifiable for the SDE model given individual trajectories, unlike for the ODE model. And not only the product *θ*_2_*m*_0_scale but also the parameter scale and the product *θ*_2_*m*_0_ are globally identifiable for the SDE model. Moreover, for the simulated data, the 95% CIs for the identifiable parameters for the SDE model covered the true parameter value adequately many times. Whereas for the ODE model, the true parameter values for the parameters *θ*_1_, *θ*_3_, and *θ*_2_*m*_0_scale were not covered by the 95% CIs for many of the posterior samples and were sometimes not even included in the range of values in the sample. The fact that the parameters *θ*_1_ and *θ*_3_ can be adequately determined using the SDE modeling approach given an individual trajectory renders the multi-experiment approach with different mRNA constructs and the computationally intense hierarchical optimization algorithm used in Fröhlich et al. (2018) unnecessary in the case that the determination of these parameters is the main objective. Besides, assuming that an MJP is the most appropriate description of the underlying dynamics, we saw that the estimated parameter values for a single cell trajectory based on the ODE model cannot be trusted even when narrow 95% CIs suggest low uncertainty. While the SDE model is clearly superior in terms of the information that we are able to extract from a single trajectory about the parameters that determine the dynamics of the underlying process, it has nevertheless several disadvantages. First of all, we were not able to include the estimation of the initial time point *t*_0_ of mRNA release into the Stan sampling procedure due to technical/methodological limitations. Other sampling approaches such as particle MCMC (Golightly & Wilkinson, 2011) might alleviate this problem, but to our knowledge, no examples of inferring an unknown time point for SDE models have been investigated so far and would thus require further work. Another drawback of the SDE model are the higher computational costs as we need to sample from a higher-dimensional distribution (due to the random process values) than for the ODE model. For the SDE model, the sampling in our study takes on average almost 5.5 hours while for the ODE model, it averages at about 20 minutes. Note, however, that for the ODE model, the NUTS method implemented in Stan generated only chains that sample only from one of the posterior modes. Therefore, one might argue that sampling did not properly converge in this example. In general, estimation procedures for SDE models are more complex and unlike for ODE models, publicly available software tools are rare and usually not generally applicable. There is a clear need to further develop such tools for SDE models in order to harness their full potential, especially with regard to better identifiability of kinetic parameters. On the other hand, combining both modeling approaches as we have done here by first determining *t*_0_ based on the ODE model and then estimating the kinetic parameters based on the SDE model is certainly also meaningful. Overall, our study clearly demonstrates the relevance of considering different modeling approaches and of selecting the appropriate one. We expect to see stochastic models being used more frequently as increasing computing power becomes available and facilitates inference for such models.

## Declarations

### Availability of data and material

The experimental data used in this study had previously been published with the article Fröhlich et al. (2018). The simulated data can be generated with the code referenced below.

### Code availability

The source code used to produce the computational results is available on GitHub at https://github.com/fuchslab/Translation_kinetics_after_transfection.

### Authors’ contributions

All authors designed the study. SP implemented the used methods, carried out the analysis and drafted the manuscript. CF and JH provided supervision. All authors contributed to the final version of the manuscript and gave final approval for publication.

