## Supplementary material for "Identifiability analysis for models of the translation kinetics after mRNA transfection"

### A.1 General Itô diffusion processes

In the main article, we consider an example of a parametric time-homogeneous Itô diffusion process. In general, a $d$-dimensional *time-homogeneous Itô diffusion process* $(\boldsymbol{X}_t)_{t\geq 0}$ is a stochastic process that fulfills the following SDE:

$$\mathrm{d}\boldsymbol{X}_t = \boldsymbol{\mu}\left(\boldsymbol{X}_t, \boldsymbol{\theta}\right)\, \mathrm{d}t + \boldsymbol{\sigma}\left(\boldsymbol{X}_t, \boldsymbol{\theta}\right)\, \mathrm{d}\boldsymbol{B}_t, \qquad \boldsymbol{X}_0 = \boldsymbol{x}_0, \tag{8}$$

with state space $\mathcal{X} \subseteq \mathbb{R}^d$, starting value $\boldsymbol{x}_0 \in \mathcal{X}$, and an $r$-dimensional Brownian motion $(\boldsymbol{B}_t)_{t\geq 0}$. The model parameter $\boldsymbol{\theta} \in \Theta$ is from an open set $\Theta \subseteq \mathbb{R}^p$. The function $\boldsymbol{\mu} : \mathbb{R}^d \times \Theta \to \mathbb{R}^d$ is usually called the drift coefficient and $\boldsymbol{\sigma} : \mathbb{R}^d \times \Theta \to \mathbb{R}^{d\times r}$ the diffusion coefficient. Equation (8) is a symbolic way of writing the stochastic integral equation

$$\boldsymbol{X}_t = \boldsymbol{x}_0 + \int_0^t \boldsymbol{\mu}\left(X_s, \boldsymbol{\theta}\right)\, \mathrm{d}s + \int_0^t \boldsymbol{\sigma}\left(\boldsymbol{X}_s, \boldsymbol{\theta}\right)\, \mathrm{d}\boldsymbol{B}_s \quad \text{for all } t \geq 0 \quad \mathbb{P}\text{-almost surely},$$

where the first integral is an ordinary Riemann integral and the second integral is a stochastic integral in the Itô sense. In the remainder of this section, we omit the dependence of $\boldsymbol{\mu}$ and $\boldsymbol{\sigma}$ on the parameter $\boldsymbol{\theta}$ and briefly state two important tools for handling SDE models of this type. Elaborate and general introductions to SDEs can be found e. g. in Øksendal (2003), Fuchs (2013), and Braumann (2019).

The Itô integral and thus also Itô diffusion processes do not adhere to the rules of classical calculus. Instead, the following theorem states the stochastic counterpart of the chain rule from classical calculus which is known as *Itô formula*. The formulation of the Itô formula specific for Itô diffusion processes as we state it here follows directly from the general Itô formula as stated in Øksendal (2003, Chapter 4.2).

**Theorem A.1** (Itô formula). *Let $\boldsymbol{X}_t$ be a $d$-dimensional Itô diffusion process described by an SDE as in (8). Let $g(t, \boldsymbol{x}) = (g_1(t, \boldsymbol{x}), \ldots, g_q(t, \boldsymbol{x}))$ be a map from $[0, T] \times \mathbb{R}^d$ into $\mathbb{R}^q$ with continuous first-order partial derivatives in $t$ and continuous first- and second-order partial derivatives in $\boldsymbol{x}$. Then the process*

$$\boldsymbol{Y}(t, \omega) = g(t, \boldsymbol{X}_t)$$

*is an Itô process whose $k^{th}$ component $\boldsymbol{Y}^{(k)}$ is given by*

$$\begin{aligned}\mathrm{d}\boldsymbol{Y}^{(k)} &= \frac{\partial g_k}{\partial t}(t, \boldsymbol{X})\, \mathrm{d}t + \sum_{i=1}^{q} \frac{\partial g_k}{\partial x^{(i)}}(t, \boldsymbol{X})\, \mathrm{d}\boldsymbol{X}^{(i)} + \frac{1}{2}\sum_{i=1}^{q}\sum_{j=1}^{q} \frac{\partial^2 g_k}{\partial x^{(i)} \partial x^{(j)}}(t, \boldsymbol{X})\, \mathrm{d}\boldsymbol{X}^{(i)} \cdot \mathrm{d}\boldsymbol{X}^{(j)}, \\ &= \left( \frac{\partial g_k}{\partial t}(t, \boldsymbol{X}) + \boldsymbol{\mu}(\boldsymbol{X})^{Tr} \nabla g_k(t, \boldsymbol{X}) + \frac{1}{2}\mathrm{trace}\left(\boldsymbol{\sigma}(\boldsymbol{X})\boldsymbol{\sigma}(\boldsymbol{X})^{Tr} \nabla\left(\nabla g_k(t, \boldsymbol{X})\right)\right)\right) \mathrm{d}t \\ &\quad + \left(\nabla g_k(t, \boldsymbol{X})\right)^{Tr} \boldsymbol{\sigma}(\boldsymbol{X})\, \mathrm{d}\boldsymbol{B}_t,\end{aligned} \tag{9}$$

*where $\nabla g_k$ denotes the gradient of $g_k$ with respect to the components of $\boldsymbol{x}$ and $\mathrm{d}\boldsymbol{X}^{(i)} \cdot \mathrm{d}\boldsymbol{X}^{(j)}$ is computed according to the rules $\mathrm{d}\boldsymbol{B}^{(i)} \cdot \mathrm{d}t = \mathrm{d}t \cdot \mathrm{d}\boldsymbol{B}^{(j)} = (\mathrm{d}t)^2 = 0$ and $\mathrm{d}\boldsymbol{B}^{(i)} \cdot \mathrm{d}\boldsymbol{B}^{(j)} = \delta_{ij}\, \mathrm{d}t$ with $\delta_{ij}$ denoting the Kronecker delta.*

Most SDEs do not have an analytical solution and their transition densities are not explicitly known. Instead, numerical approximation schemes are used for the solution of the SDEs. Kloeden and Platen (1992) provide a detailed description of these methods. The most commonly used approximation is the *Euler(-Maruyama) scheme*. It can be conveniently written in vector notation and approximates the $d$-dimensional solution $(\boldsymbol{X}_t)_{t\geq 0}$ of an SDE by setting $\boldsymbol{Y}_0 = \boldsymbol{x}_0$ and, then, successively calculating the following:

$$\boldsymbol{Y}_{k+1} = \boldsymbol{Y}_k + \boldsymbol{\mu}\left(\boldsymbol{Y}_k\right) \Delta t_k + \boldsymbol{\sigma}\left(\boldsymbol{Y}_k\right) \Delta \boldsymbol{B}_k, \tag{10}$$

where $\Delta t_k = t_{k+1} - t_k$, $\Delta \boldsymbol{B}_k = \boldsymbol{B}_{t_{k+1}} - \boldsymbol{B}_{t_k}$, and $\boldsymbol{Y}_k$ is the approximation of $\boldsymbol{X}_{t_k}$ for $k = 0, 1, 2, \ldots$. Since the Euler scheme is a linear transformation of the normally-distributed increments $\Delta \boldsymbol{B}_k \sim \mathcal{N}(\boldsymbol{0}, \Delta t_k \boldsymbol{I}_r)$ of the Brownian motion, where $\boldsymbol{I}_r$ denotes the $r$-dimensional identity matrix, the process state $\boldsymbol{Y}_{k+1}$ conditioned on $\boldsymbol{Y}_k$ is also normally-distributed with

$$\boldsymbol{Y}_{k+1} \,|\, \boldsymbol{Y}_k \;\sim\; \mathcal{N}\left(\boldsymbol{Y}_k + \boldsymbol{\mu}(\boldsymbol{Y}_k)\,\Delta t_k\,,\, \boldsymbol{\sigma}(\boldsymbol{Y}_k)\,\boldsymbol{\sigma}^{\mathrm{Tr}}(\boldsymbol{Y}_k)\,\Delta t_k\right),$$

where $\mathcal{N}(\boldsymbol{a}, \boldsymbol{b})$ denotes the multivariate normal distribution with mean vector $\boldsymbol{a} \in \mathbb{R}^d$ and covariance matrix $\boldsymbol{b} \in \mathbb{R}^{d \times d}$.

### A.2 Investigating the need for data augmentation for the SDE model

In this section, we focus on the inference problem for the SDE model and investigate whether the amount of data that we have available ($K = 181$ observations per cell with time step $\Delta t = 1/6$ hours) is sufficient for the Euler approximation to be appropriate, i. e. whether the step size between observations is small enough. We simulate one trajectory of the MJP described in Section 3.1 with parameters $\theta_1 = 0.11$, $\theta_2 = 0.3$, $\theta_3 = 0.09$, and $m_0 = 200$ on the time interval $[0, 30]$ using Gillespie's algorithm and use observations at 181 equidistant time points. We assume for now that the amount $X_2$ of GFP is directly observed without error and that for the amount $X_1$ of mRNA, we only observe the initial value $m_0 = 200$. All observations are without measurement error and we assume $t_0 = 0$ to be known. Thus, we only estimate the kinetic parameters $\boldsymbol{\theta}$ for the SDE model, and to this end, use Stan and Bayesian data augmentation with different numbers of inter-observation intervals which means that we impute additional (artificial) data points between every two observations and these points are treated as additional parameters in the estimation procedure (for a detailed description of Bayesian data augmentation see Fuchs, 2013). A number of inter-observation intervals of 1 means that we do not impute any points between observations. A number of 2 inter-observation intervals means that we impute one point between every two observations and so on. We generated 4 HMC chains with 1000 iterations after warm-up each. Figure 14 shows the median of the obtained posterior sample as the point estimates and the CIs for the three kinetic parameters and for different numbers of inter-observation intervals. Evidently, the estimation results do not improve when increasing the number of inter-observation intervals. Therefore, we conclude that data augmentation is not necessary and do not make use of data augmentation in the main part of the article.

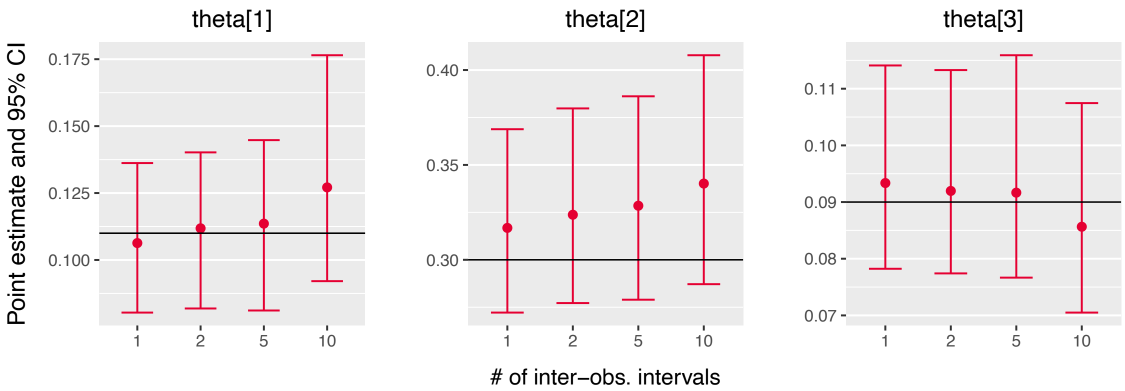


Figure 14: Point estimates (median of the posterior sample) and 95% CIs for the kinetic parameters estimated with Stan and Bayesian data augmentation for different numbers of inter-observations intervals. The black line represents the true parameter values with which the data was generated.

### A.3 Hamiltonian Monte Carlo (HMC) methods and Stan

#### A.3.1 Brief introduction to the algorithm

To sample from the posterior densities of the two model types (ODE and SDE) as formulated in Section 6 of the main article, we use the open source software Stan (Carpenter et al., 2017). Stan provides an implementation of the Hamiltonian Monte Carlo (HMC) based No-U-Turn Sampler (NUTS) to which we give a very brief introduction here which mainly draws from the description in Gelman et al. (2013). Neal (2011) gives a more detailed account. HMC methods (originally called *hybrid* Monte Carlo methods by Duane et al. (1987)) are a class of Markov chain Monte Carlo (MCMC) methods. The computational cost in each iteration for HMC methods is higher than for other MCMC methods such as Gibbs sampling or Metropolis-Hastings algorithms because HMC makes use of the derivative of the target distribution. But by that, transitions between the chain states can be generated that efficiently span the (with respect to the target distribution) important regions of the state space. By taking into account the information of the gradient, HMC avoids the random walk behavior and difficulties caused by distributions with high correlations that other MCMC methods exhibit.

Assume we want to sample from the $p$-dimensional distribution $\pi(\boldsymbol{\theta})$ for parameter $\boldsymbol{\theta} \in \mathbb{R}^p$. Motivated by the physical concept of Hamiltonian dynamics, HMC introduces an auxiliary momentum variables $\boldsymbol{\rho} \in \mathbb{R}^p$ and draws from a joint density $p(\boldsymbol{\theta}, \boldsymbol{\rho}) = p(\boldsymbol{\rho} \,|\, \boldsymbol{\theta})\pi(\boldsymbol{\theta})$. The joint density defines the so-called Hamiltonian

$$H(\boldsymbol{\theta}, \boldsymbol{\rho}) = -\log p(\boldsymbol{\theta}, \boldsymbol{\rho}) = -\log p(\boldsymbol{\rho} \,|\, \boldsymbol{\theta}) - \log \pi(\boldsymbol{\theta}) = K(\boldsymbol{\theta}, \boldsymbol{\rho}) + V(\boldsymbol{\theta}) \tag{11}$$

that describes the total energy of the system and is equal to the sum of the kinetic energy $K$ and the potential energy $V$. In HMC, the distribution of $\boldsymbol{\rho}$ is usually chosen to be independent of $\boldsymbol{\theta}$. A common choice is $\boldsymbol{\rho} \sim \mathcal{N}(\mathbf{0}_p, \boldsymbol{M})$, where $\mathcal{N}(\mathbf{0}_p, \boldsymbol{M})$ denotes the multivariate normal distribution with mean vector $\mathbf{0}_p$ and covariance matrix $\boldsymbol{M} \in \mathbb{R}^{p \times p}$, and $\boldsymbol{M}$ is called the *design* or (by analogy to the physical model) *mass matrix* and often chosen to be a diagonal matrix. Thus, the kinetic energy becomes

$$K(\boldsymbol{\rho}) = \boldsymbol{\rho}^{\mathsf{Tr}} \boldsymbol{M}^{-1} \boldsymbol{\rho}/2, \tag{12}$$

where $\boldsymbol{M}^{-1}$ denotes the inverse matrix of $\boldsymbol{M}$.

In each iteration of the HMC algorithm, a momentum $\boldsymbol{\rho}$ is sampled (e. g. from $\mathcal{N}(\mathbf{0}_p, \boldsymbol{M})$) and then by analogy to the physical model of the frictionless movement of a marble with position $\boldsymbol{\theta}$ and momentum $\boldsymbol{\rho}$ (describing the marble's mass and velocity) across a surface, the dynamics, i. e. the changes in position and momentum, that preserve the total energy are described by the Hamiltonian equations

$$\begin{aligned} \frac{d\rho_i}{dt} &= -\frac{\partial H}{\partial \theta_i}, \\ \frac{d\theta_i}{dt} &= \frac{\partial H}{\partial \rho_i} \end{aligned}$$

for $i = 1, \ldots, p$. With the choice of $H$, $K$, and $V$ as in Equations (11) and (12), we have

$$\begin{aligned} \frac{d\boldsymbol{\rho}}{dt} &= -\nabla_{\boldsymbol{\theta}} V(\boldsymbol{\theta}) &&= \nabla_{\boldsymbol{\theta}} \log \pi(\boldsymbol{\theta}), \\ \frac{d\boldsymbol{\theta}}{dt} &= \nabla_{\boldsymbol{\rho}} K(\boldsymbol{\rho}) &&= \boldsymbol{M}^{-1}\boldsymbol{\rho}, \end{aligned} \tag{13}$$

where $\nabla_{\boldsymbol{x}}$ denotes the gradient with respect to $\boldsymbol{x}$. In each iteration, Equations (13) are numerically integrated to obtain proposals $\boldsymbol{\theta}^*$ and $\boldsymbol{\rho}^*$. A common choice of the numerical integrator is the leap-frog method. Then, an accept-reject step is performed analogously to the Metropolis-Hastings algorithm.

We summarize the HMC steps in Algorithm 1.

---

**Algorithm 1:** Hamiltonian Monte Carlo algorithm (with leap-frog integrator)

---

**Input:** *A target density* $\pi(\cdot)$*, an initial state* $\boldsymbol{\theta}^{(0)}$*, number of iterations* $n$*, mass matrix* $\boldsymbol{M}$*, and step size* $\epsilon$ *and number* $L$ *of steps for numerical integration.*

In each iteration $i = 1, \ldots, n$:

**Step 1** Generate $\boldsymbol{\rho} \sim \mathcal{N}(\mathbf{0}_p, \boldsymbol{M})$ and set $\boldsymbol{\theta}^* \leftarrow \boldsymbol{\theta}^{(i-1)}$ and $\boldsymbol{\rho}^* \leftarrow \boldsymbol{\rho}$.

**Step 2** Repeat $L$ leap-frog steps by setting:

$$
\begin{aligned}
\boldsymbol{\rho}_{\frac{1}{2}} &\leftarrow \boldsymbol{\rho}^* + \frac{1}{2}\epsilon \nabla_{\boldsymbol{\theta}} \log \pi(\boldsymbol{\theta}^*) \\
\boldsymbol{\theta}^* &\leftarrow \boldsymbol{\theta}^* + \epsilon M^{-1} \boldsymbol{\rho}_{\frac{1}{2}} \\
\boldsymbol{\rho}^* &\leftarrow \boldsymbol{\rho}_{\frac{1}{2}} + \frac{1}{2}\epsilon \nabla_{\boldsymbol{\theta}} \log \pi(\boldsymbol{\theta}^*)
\end{aligned}
$$

**Step 3** Accept $\boldsymbol{\theta}^*$ as $\boldsymbol{\theta}^{(i)}$ with probability

$$
\alpha(\boldsymbol{\theta}^{(i-1)}, \boldsymbol{\rho}, \boldsymbol{\theta}^*, \boldsymbol{\rho}^*) = \min\left[1, \exp\left(H(\boldsymbol{\theta}^{(i-1)}, \boldsymbol{\rho}) - H(\boldsymbol{\theta}^*, \boldsymbol{\rho}^*)\right)\right],
$$

if $\boldsymbol{\theta}^*$ is rejected $\boldsymbol{\theta}^{(i)} := \boldsymbol{\theta}^{(i-1)}$.

**Output:** *A sample* $\{\boldsymbol{\theta}^{(1)}, \ldots, \boldsymbol{\theta}^{(n)}\}$ *approximately distributed according to* $\pi(\cdot)$*.*

---

Two of the limitations of this general HMC algorithm are on the one hand that due to the use of the derivative with respect parameter, it is only suitable for continuous distributions, and on the other hand, the choice of the tuning parameters is of crucial importance to the performance of the algorithm and can be cumbersome. The tuning parameters include the mass matrix $M$, and step size $\epsilon$ and number $L$ of steps for numerical integration.

An extension of HMC, the *No-U-Turn Sampler (NUTS)*, introduced by Hoffman and Gelman (2014) includes a way to automatically determine the number $L$ of steps for numerical integration using an recursive algorithm that grows a binary tree representing leap-frog steps forward and backward in time which is stopped as soon as further steps do no longer increase the distance between a newly explored point and the original starting point (i. e. as soon as the steps start to make a U-turn).

The open-source Bayesian inference package *Stan* which we make use of through its R interface `rstan` (Stan Development Team, 2019) provides an efficient C++ implementation of NUTS. In Stan, the gradient of the log-posterior distribution is calculated (exactly) by reverse-mode automatic differentiation (Carpenter et al., 2015). Moreover, Stan can automatically optimize the step size $\epsilon$ to match a (user-defined) acceptance-rate target based on dual averaging as proposed by Nesterov (2009) and it also estimates the mass matrix $M$ during a warm-up phase consisting of several stages.

#### A.3.2 Evaluating (general) MCMC output

While in theory, any MCMC method (for which convergence of the transition kernel is ensured) will give a sample from the target distribution if infinitely many iterations are executed; in practice, the sample size can only be finite which makes it necessary to carefully evaluate the MCMC output.

A quantity that can be used to quantify the degree of convergence when several chains have been simulated is the $\hat{R}$ value. The $\hat{R}$ convergence (or rather stationarity) diagnostic compares the between- and within-chain variance for individual model parameters and other univariate quantities of interest. Assume we are considering the scalar parameter $\psi$ for which we have simulations $\psi_{i,j}$ for $i = 1, \ldots, n$ and $j = 1, \ldots, m$ and for $m$ chains (after discarding the warm-up iterations and then splitting each simulated chain in half) of length $n$. Let

$$
\widehat{\text{var}}^+(\psi \mid \mathcal{D}) = \frac{n-1}{n} W + \frac{1}{n} B \tag{14}
$$

be an estimate for the marginal posterior variance of $\psi$, where the *within-sequence variance* $W$ is defined by

$$
W = \frac{1}{m} \sum_{j=1}^{m} s_j^2 \qquad \text{with} \qquad s_j^2 = \frac{1}{n-1} \sum_{i=1}^{n} (\psi_{ij} - \bar{\psi_{\cdot j}})^2,
$$

and the *between-sequence variance* $B$ is defined by

$$B = \frac{n}{m-1}\sum_{j=1}^{m}(\bar{\psi}_{\cdot j} - \bar{\psi}_{\cdot\cdot})^2 \qquad \text{with} \quad \bar{\psi}_{\cdot j} = \frac{1}{n}\sum_{i=1}^{n}\psi_{ij} \quad \text{and} \quad \bar{\psi}_{\cdot\cdot} = \frac{1}{m}\sum_{j=1}^{m}\bar{\psi}_{\cdot j}.$$

Then, $\hat{R}$ is defined as

$$\hat{R} = \sqrt{\frac{\widehat{\mathrm{var}}^{+}(\psi \mid \mathcal{D})}{W}}.$$

Due to the splitting of chains in half, $\hat{R}$ calculated in this way is also known as split-$\hat{R}$ and was suggested in Gelman et al. (2013). The value can be interpreted as the factor by which the scale of the distribution of the current simulations for $\psi$ can be reduced by continuing the number of iterations to infinity. If chains have mixed well, $\hat{R}$ is close to 1. Gelman et al. (2013) state that values up to $1.1$ are acceptable. The $\hat{R}$ reported by Stan is calculated as the maximum of a so-called rank-normalized split-$\hat{R}$ and a rank-normalized folded-split-$\hat{R}$ which was recently suggested by Vehtari et al. (2021).

Another issue in MCMC sampling is the fact that the draws are not independent but may even be highly correlated. It is important to keep in mind that such a correlated sample from the parameter posterior distribution does not contain the same amount of information as an independent and identically distributed sample. This issue is addressed by the notion of the effective sample size (ESS). The ESS of a sample of correlated draws quantifies the size of a corresponding independent and identically distributed sample that contains the same amount information.

The ESS for a sample of scalar parameter $\psi$ consisting of $m$ chains each of length $n$ (again after discarding warm-up iterations but without splitting of the chains) can be defined as

$$n_{\mathrm{eff}} = \frac{mn}{1 + 2\sum_{t=1}^{\infty}\rho_t},$$

where $\rho_t$ is the autocorrelation of the sequence $\psi$ at lag $t$. This quantity can be approximated in different ways. Here, we give the approximation that is presented in Gelman et al. (2013) and implemented in `rstan`. The estimated autocorrelations $\hat{\rho}_t$ are computed as

$$\hat{\rho}_t = 1 - \frac{V_t}{2\widehat{\mathrm{var}}^{+}(\psi \mid \mathcal{D})}$$

for $t = 1, \ldots, T$ and, where the estimate $\widehat{\mathrm{var}}^{+}$ for the marginal posterior variance is calculated as in (14) and the variogram $V_t$ at lag $t$ is calculated as

$$V_t = \frac{1}{m(n-t)}\sum_{j=1}^{m}\sum_{i=t+1}^{n}(\psi_{i,j} - \psi_{i-t,j})^2.$$

The maximal considered lag $T$ is chosen to be the first odd positive integer for which $\hat{\rho}_{T+1} + \hat{\rho}_{T+2}$ is negative and finally, the ESS is approximated by

$$\hat{n}_{\mathrm{eff}} = \frac{mn}{1 + 2\sum_{t=1}^{T}\hat{\rho}_t}.$$

Gelman et al. (2013) recommend that a minimum ESS of 10 per simulated chain is achieved. The between-chain information is taken into account in the calculation of $\hat{n}_{\mathrm{eff}}$ by including the term $\widehat{\mathrm{var}}^{+}(\psi \mid \mathcal{D})$. Thus, the ESS is affected when we try to sample from multi-modal distributions. In fact, in the case of well-separated modes and each chain sampling only from one of these modes, the ESS roughly equals to the number chains divided by the number of modes.

#### A.3.3 Further diagnostics of MCMC output specific to HMC and NUTS

In addition to the quality indicators for MCMC output mentioned in the previous section, Stan reports further quantities that are specific to HMC and NUTS and are of interest to assess sampling efficiency. These include the number of divergent transitions, the tree depth, and the (energy) Bayesian fraction of missing (BFMI) which we briefly describe below. See the Stan reference manual for more detailed explanations (Stan Development Team, 2019).

Integrating the Hamiltonian equations (13) in Section A.3.1 analytically would preserve the value of the Hamiltonian $H(\boldsymbol{\theta}, \boldsymbol{\rho})$; however, since analytical integration is not possible for most problems of interest, the equations are numerically integrated which leads to numerical errors. If the difference between $H(\boldsymbol{\theta}, \boldsymbol{\rho})$ of the starting point and $H(\boldsymbol{\theta}^*, \boldsymbol{\rho}^*)$ of the proposed point at the end of the simulated Hamiltonian trajectory becomes too large (where the default threshold is $10^3$), Stan will classify the starting point as one of a *divergent transition*. If many of such starting points of divergent transitions are concentrated within a region of parameter space, this may be an indication that the curvature of the posterior is very high in this region and that the step size $\epsilon$ is too large to adequately explore this region.

As briefly mentioned in Section A.3.1, NUTS builds up a binary tree when determining the number $L$ of leapfrog steps to take before a U-turn would occur. Stan records the depth of this tree for each iteration and thus also the corresponding starting point. Moreover, the user can specify a maximum tree depth $d$ to avoid long execution times due too many steps; as at most $2^{d-1}$ leapfrog steps are taken in each iteration. The default value is $d = 10$. Hitting this maximum means that NUTS is terminated prematurely (i. e. more steps would have been possible before a U-turn) and Stan counts how many times this occurs. Reasons for having to take many steps may be a too small step size due to poor adaptation to a posterior of varying curvature or targeting a very high acceptance rate.

According to Betancourt et al. (2015), the *BFMI* indicates how well the energy sets of the Hamiltonian are explored. Let $E = H(\boldsymbol{\theta}, \boldsymbol{\rho})$ be the total energy, $\pi(E|\rho)$ the energy transition distribution, and $\pi(E)$ the marginal energy distribution. If $\pi(E|\rho)$ is substantially more narrow than $\pi(E)$, then a HMC chain may not be able to completely explore the tails of the target distribution. The BFMI quantifies the mismatch between the two distributions and is defined and approximated by

$$BFMI := \frac{\mathbb{E}_\pi \left[ Var_{\pi_{E|\rho}}[E|\rho] \right]}{Var_{\pi_E}[E]} \approx \frac{\sum_{n=1}^{N}(E_n - E_{n-1})^2}{\sum_{n=0}^{N}(E_n - \bar{E})^2} =: \widehat{BFMI}.$$

The Stan development team recommends to ensure that the value of $\widehat{BFMI}$ is greater than 0.2.

### A.4 Additional results for the posterior sampling

#### A.4.1 Additional sampling results for simulated data

Figures 15, 16, 17, and 18 show the same sampling output (the four posterior samples for the two simulated data sets depicted in Figure 5) as Figures 6, 7, 9, and 10 in Section 6.2; however here, the results are not compared between the ODE and the SDE model but between simulated data with and without measurement error.

For the SDE, we see in Figure 15 that the occurrence of measurement error substantially impacts the distribution of the posterior sample with respect to the parameters $\theta_1$ and $\theta_3$. The shape of the two dimensional projection changes from an elliptic shape to a banana-like shape. Especially for $\theta_3$, the 95% CI and the range of values in the posterior sample increase a lot and the true parameter value is only barely covered by the 95% CI for simulated data with measurement error.

Similarly for the parameters $\theta_2$, $m_0$, $\mathrm{scale}$ and their products, Figure 16 shows that there is quite a difference between the distributions of the posterior samples for the simulated data without and with measurement error. In particular for the parameters $\mathrm{scale}$ and $\theta_2 m_0$ which we consider to be identifiable, the 95% CIs increase substantially for data with measurement error, and also the appearance of the two-dimensional projections with respect to these two parameters changes a lot, from a slightly bent ellipse to a clear banana shape. For the product $\theta_2 m_0 \mathrm{scale}$, the dispersion of the posterior samples changes only slightly which is apparent from the similar lengths of the 95% CIs in Figure 16 and also from the similar c.v. in Tables 2 and 5 ($0.083$ for data without measurement error and $0.093$ for data with measurement error). The location of the sample measured e. g. by the median slightly shifts away from the true parameter value for the data with measurement error; however, the true value is still included in the 95% CIs. Only for parameter $m_0$ for which we also did not see much difference in the posterior samples for the ODE vs. SDE model, the occurrence of measurement error does not seem to affect the posterior sample much. For the remaining parameters $\theta_2$, $\theta_2 \mathrm{scale}$, and $m_0 \mathrm{scale}$ which we do not consider to be identifiable but for which the 95% CIs of the posterior samples for the SDE model were clearly more narrow than the 95% CIs of the corresponding posterior sample for the ODE model, the 95% CIs and ranges of values of the posterior sample for the SDE model for data with measurement error are broader than for data without measurement error.

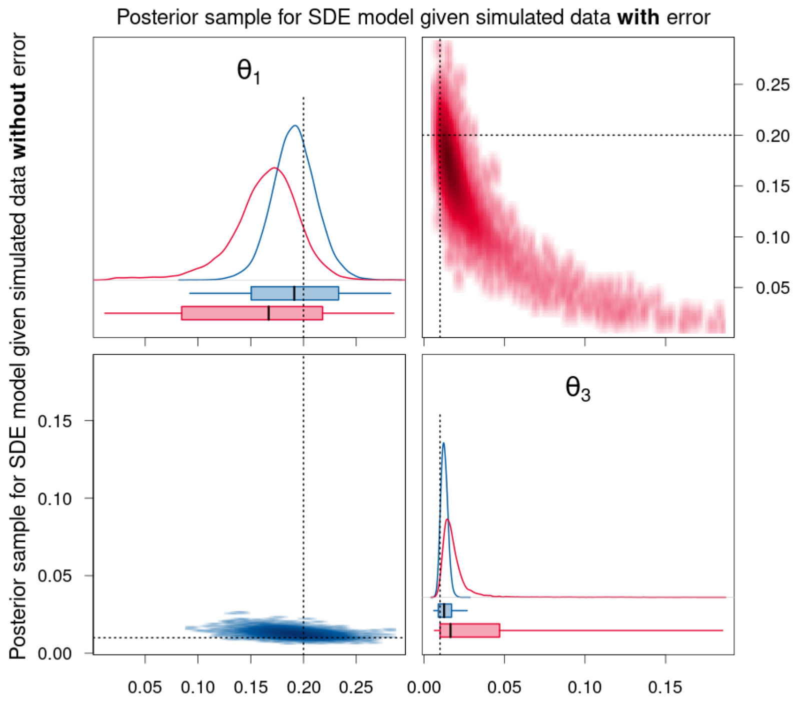


Figure 15: Density estimates of the posterior samples for parameters $\theta_1$ and $\theta_3$ for the SDE model given simulated data without (blue, lower triangle) and with (red, upper triangle) measurement error. *Diagonal panels:* Marginal densities for the respective parameter and boxplots showing the 95% CI as box, the range of the sample as whiskers, and the median as thick black line. *Off-diagonal panels:* Smoothed scatter plots of the two-dimensional projections of the samples where darker hues signify higher density values. The dotted lines represent the true parameter values that were used to simulate the data.

For the ODE model, Figures 17 and 18 show that there is hardly any difference for most of the parameters between the posterior sample for the data without and with measurement error since the majority of the parameters are not identifiable anyway. For the parameters $\mathrm{offset}$ and $t_0$, there is a slight difference. For the measurement error parameter $\sigma$, the posterior sample consists of higher values for data with measurement error as expected. Note that for both simulated datasets, the range of the posterior sample does not include the true parameter value for $\sigma$. Finally for the product $\theta_2 m_0 \mathrm{scale}$, the dispersion of the posterior sample increases only slightly for data with measurement error and the location of the sample shifts away from the true parameter value. Also for this parameter, the range of the posterior sample does not include the true parameter value for both simulated datasets.

Figure 19 shows the statistics of the posterior samples for the simulated data without and with measurement error aggregated over 100 simulated trajectories. It visualizes the last two columns of Tables 3 and 6 and compares the results of the posterior samples for the simulated data without to those with measurement error separately for the SDE and the ODE model within each plot, instead of comparing the two model types separately for each kind of data as in Figures 8 and 11.

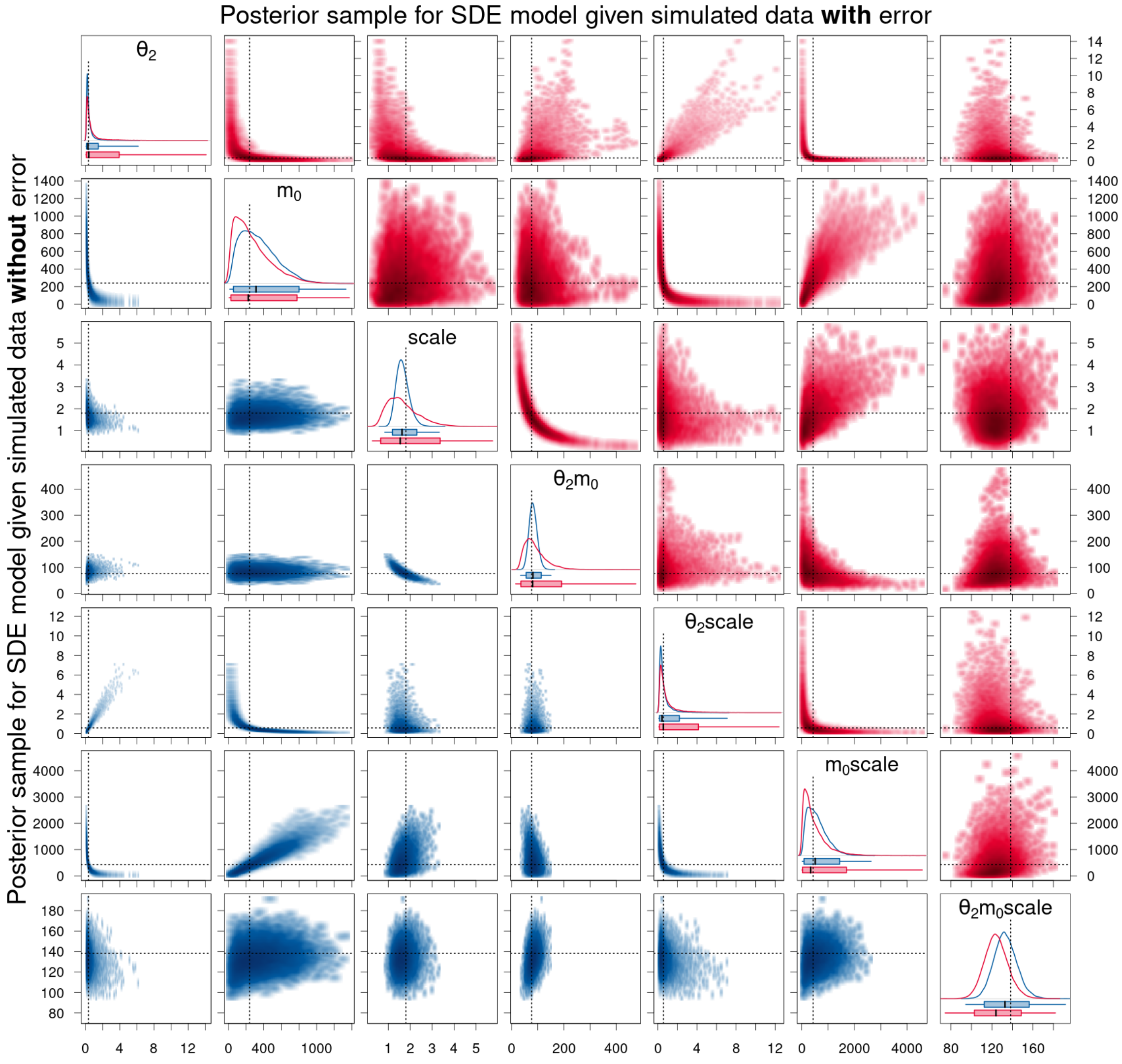


Figure 16: Density estimates of the posterior samples for parameters $\theta_2$, $m_0$, $\mathrm{scale}$, and their products for the SDE model given simulated data without (blue, lower triangle) and with (red, upper triangle) measurement error. For a detailed description of the figure's elements, see Figure 15.

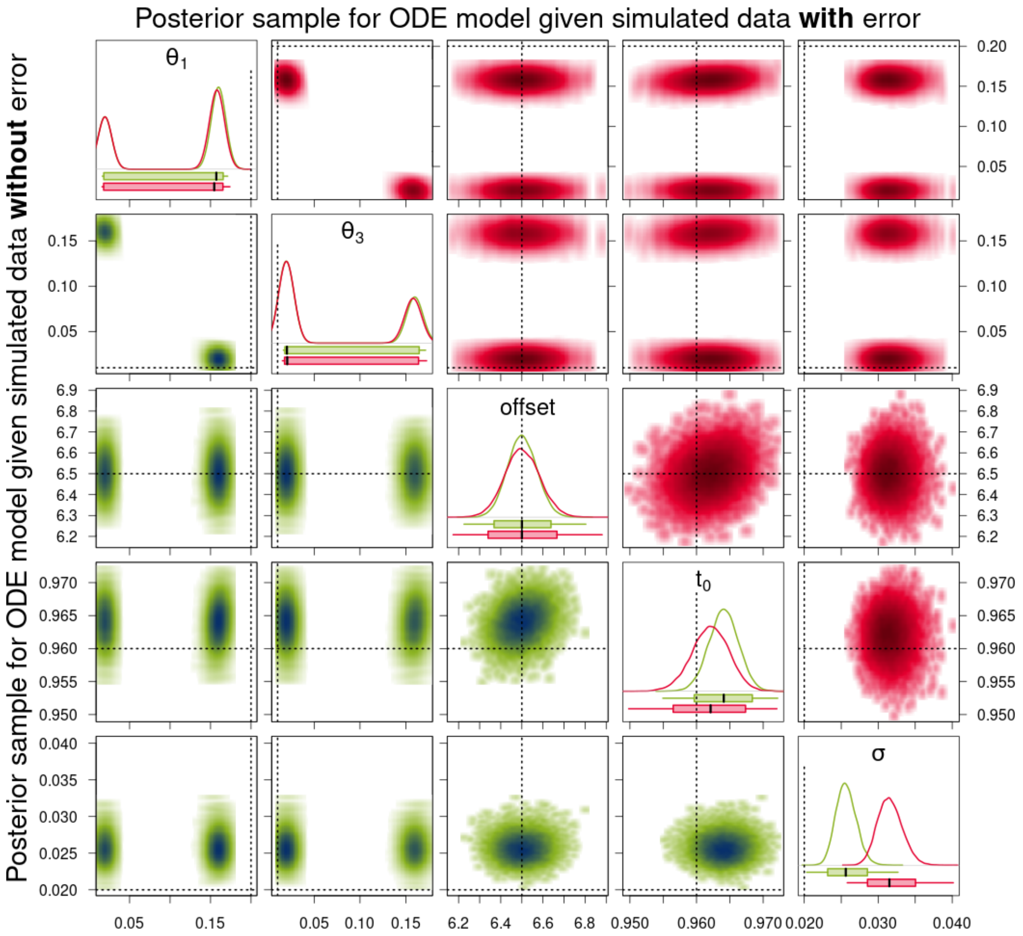


Figure 17: Density estimates of the posterior samples for parameters $\theta_1$ and $\theta_3$ for the ODE model given simulated data without (green, lower triangle) and with (red, upper triangle) measurement error. *Diagonal panels:* Marginal densities for the respective parameter and boxplots showing the 95% CI as box, the range of the sample as whiskers, and the median as thick black line. *Off-diagonal panels:* Smoothed scatter plots of the two-dimensional projections of the samples where darker hues signify higher density values. The dotted lines represent the true parameter values that were used to simulate the data. For the parameter $\sigma$, the dotted line only represents the true value for the data with measurement error. For the data without measurement error, $\sigma$ is equal to 0.

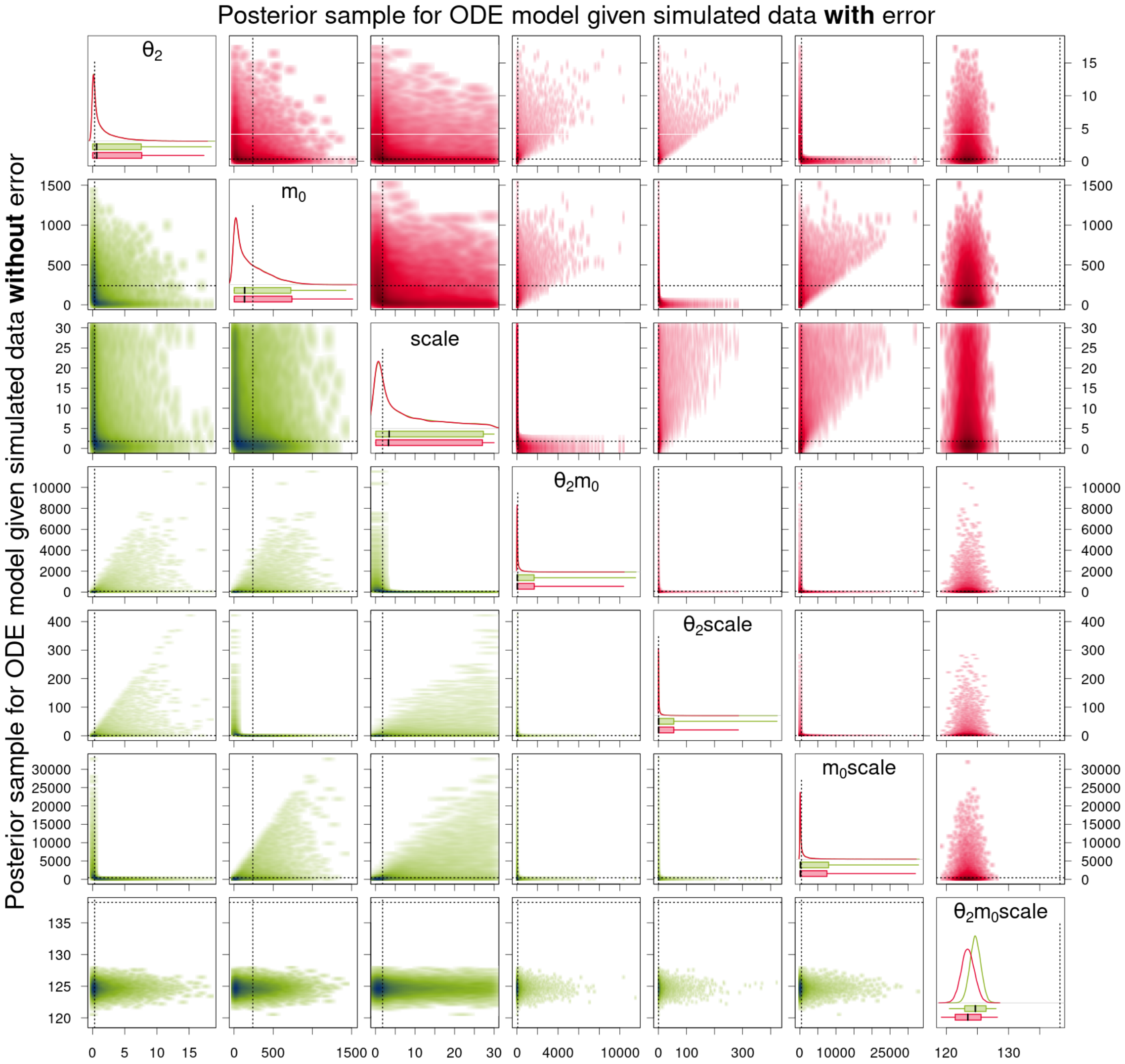


Figure 18: Density estimates of the posterior samples for parameters $\theta_2$, $m_0$, $\mathrm{scale}$, and their products for the ODE model given simulated data without (green, lower triangle) and with (red, upper triangle) measurement error. *Diagonal panels:* Marginal densities for the respective parameter and boxplots showing the 95% CI as box, the range of the sample as whiskers, and the median as thick black line. *Off-diagonal panels:* Smoothed scatter plots of the two-dimensional projections of the samples where darker hues signify higher density values. The dotted lines represent the true parameter values that were used to simulate the data.

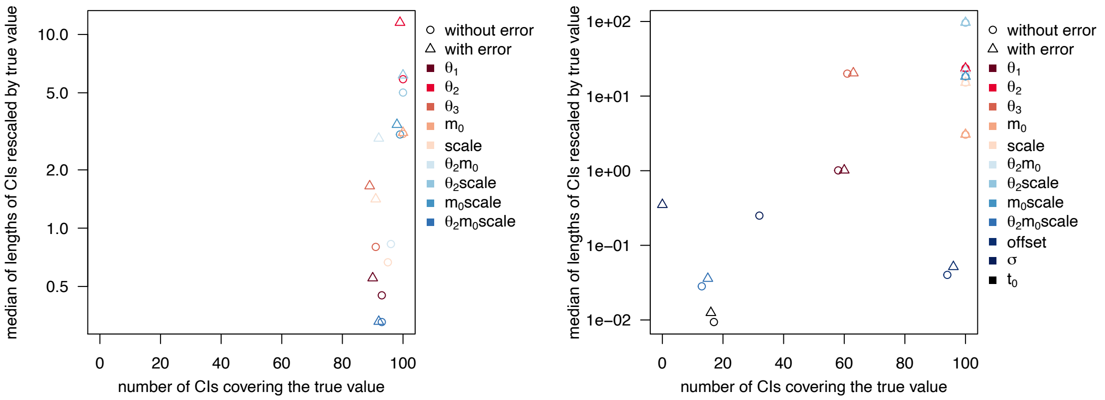


(a) SDE results

(b) ODE results

Figure 19: Statistics of posterior samples for the simulated data without and with measurement error aggregated over 100 simulated trajectories. The desirable region of value combinations is in the bottom right corner of each graph.

Figures 20 and 21 show the trace plots for the posteriors samples for the simulated data without and with measurement error that we have considered in detail in Section 6.2 of the main article.

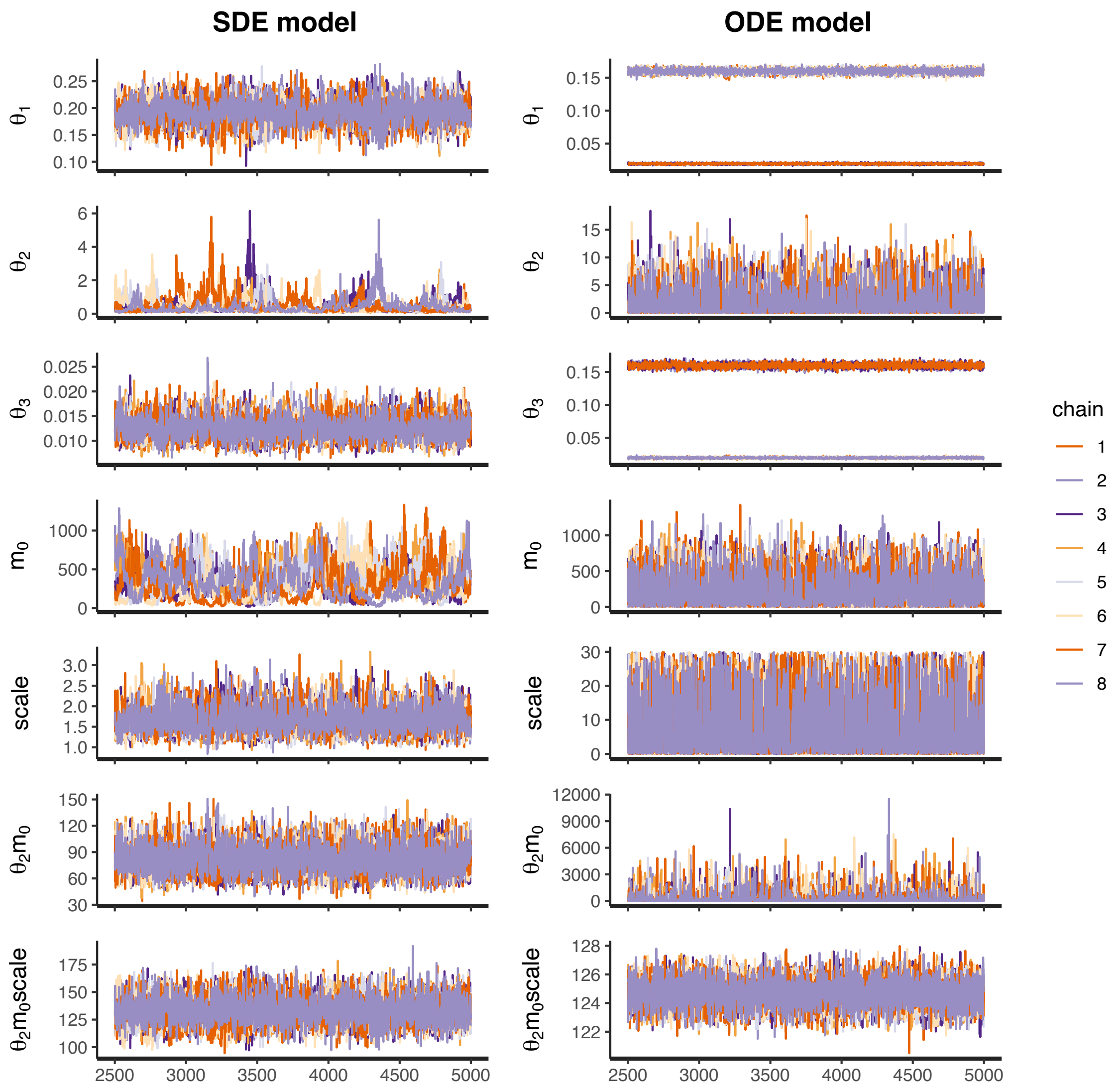


Figure 20: Trace plots of the posterior samples for the simulated data without error for the SDE model (left column) and the ODE model (right column). Both posterior samples consist of 8 chains with 2500 warm-up iterations and 2500 iterations after warm-up. Only the iterations after warm-up are displayed.

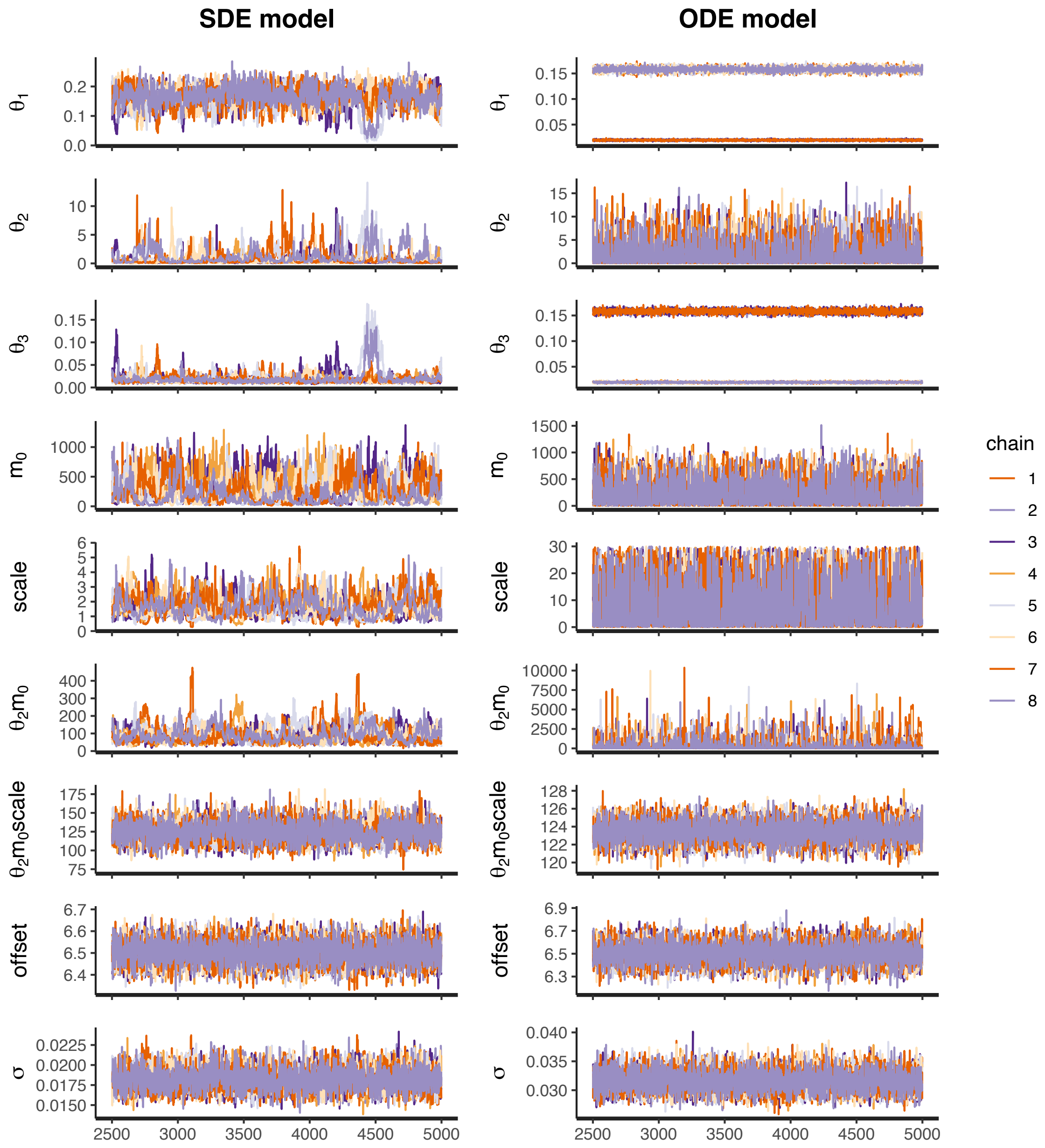


Figure 21: Trace plots of the posterior samples for the simulated data with error for the SDE model (left column) and the ODE model (right column). Both posterior samples consist of 8 chains with 2500 warm-up iterations and 2500 iterations after warm-up. Only the iterations after warm-up are displayed.

#### A.4.2 Additional sampling results for experimental dataset 1 (for eGFP)

Figure 22 shows the trace plots for the posteriors samples for the experimental data for eGFP that we have considered in detail in Section 6.3 of the main article.

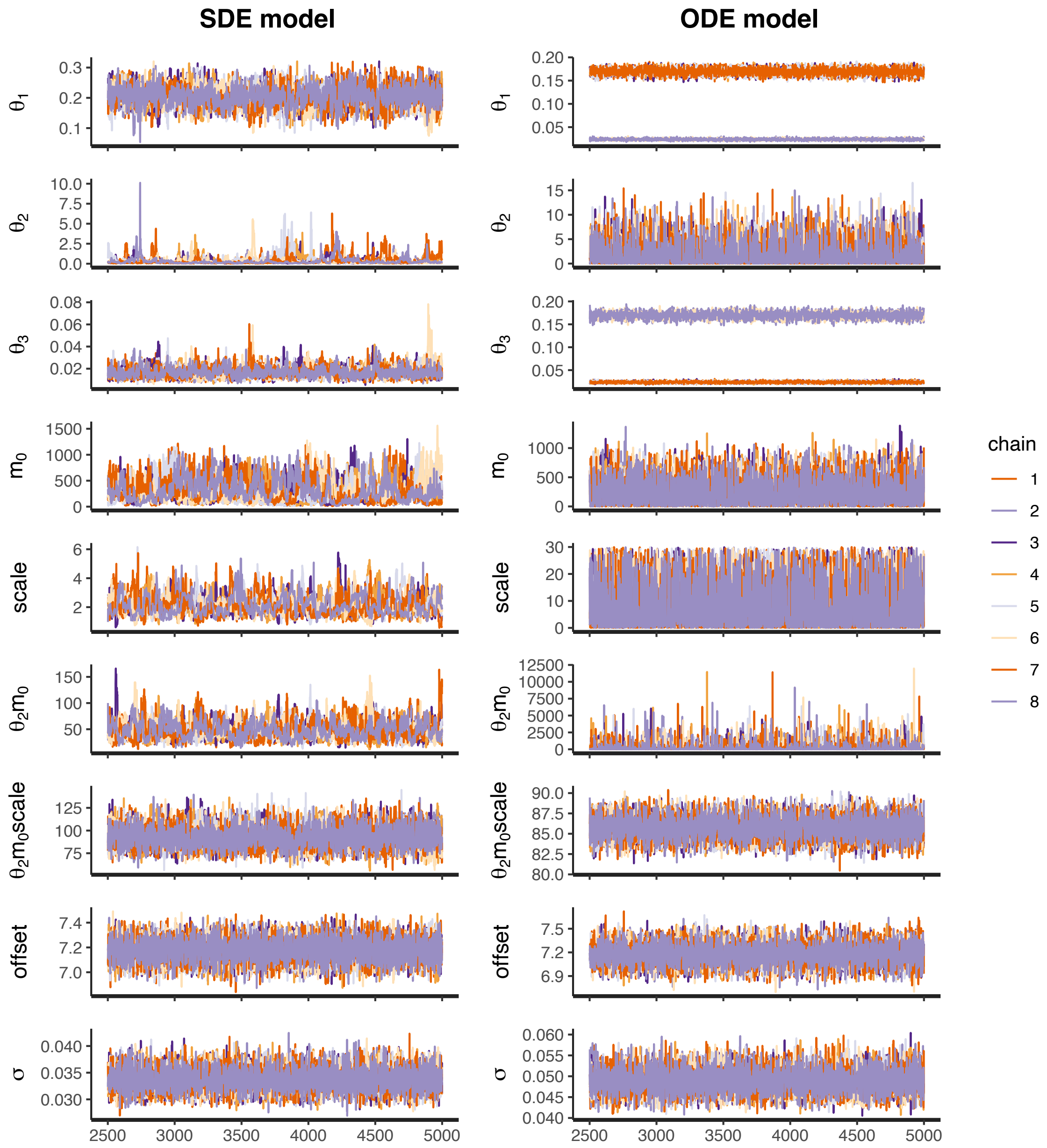


Figure 22: Trace plots of the posterior samples for the experimental data for eGFP for the SDE model (left column) and the ODE model (right column). Both posterior samples consist of 8 chains with 2500 warm-up iterations and 2500 iterations after warm-up. Only the iterations after warm-up are displayed.

#### A.4.3 Sampling results for experimental dataset 2 (for d2eGFP)

Tables 10 and 11 present a summary of the Stan output for the posterior sample of one observed trajectory for d2eGFP for the ODE and the SDE model, respectively, Figures 23 and 24 compare the density estimates of these two posterior samples, and Figure 25 shows the trace plots of these two posterior samples. Here, while of course still being symmetric, the posterior sample for the ODE model seems to be unimodal with respect to the parameters $\theta_1$ and $\theta_3$. This is due to the fact that the values of the two parameters are likely to be quite close to each other for this trajectory as can also be seen from the overlapping 95% CIs and the similar mean and median estimates for the SDE model. For the parameter $\mathrm{offset}$, the mean and median estimates from the posterior samples are very similar for the ODE and SDE model, but the 95% CI is a lot wider for the ODE model. For the measurement error parameter $\sigma$, the 95% CI for the SDE model is a lot narrower than that for the ODE model and the locations of the samples are quite far apart with a difference in the median estimates of 0.16.

Table 10: Summary of the Stan output for the ODE model given experimental data for d2eGFP. c.v. denotes the coefficient of variation and the columns headed by percentages contain the quantiles of the respective percentage value.

| | mean | c.v. | 2.5% | 50% | 97.5% | $n_{\text{eff}}$ | $\hat{R}$ |
|---|---|---|---|---|---|---|---|
| $\theta_1$ | 0.09 | 0.079 | 0.08 | 0.09 | 0.11 | 11585 | 1.00 |
| $\theta_2$ | 2.03 | 1.114 | 0.08 | 1.17 | 8.33 | 12371 | 1.00 |
| $\theta_3$ | 0.09 | 0.078 | 0.08 | 0.09 | 0.11 | 11200 | 1.00 |
| $m_0$ | 244.67 | 0.868 | 9.82 | 187.79 | 761.08 | 12127 | 1.00 |
| scale | 9.21 | 0.901 | 0.35 | 6.44 | 28.11 | 9845 | 1.00 |
| offset | 8.72 | 0.073 | 7.52 | 8.69 | 10.04 | 17557 | 1.00 |
| $t_0$ | 0.94 | 0.011 | 0.92 | 0.94 | 0.96 | 15806 | 1.00 |
| $\sigma$ | 0.17 | 0.053 | 0.15 | 0.17 | 0.18 | 16365 | 1.00 |
| $\theta_2 m_0$ | 367.06 | 1.893 | 27.97 | 121.84 | 2279.87 | 11547 | 1.00 |
| $\theta_2$scale | 12.37 | 1.907 | 1.03 | 4.19 | 80.33 | 7681 | 1.00 |
| $m_0$scale | 1756.24 | 1.569 | 94.74 | 672.69 | 10085.29 | 8815 | 1.00 |
| $\theta_2 m_0$scale | 786.93 | 0.026 | 746.72 | 786.79 | 828.01 | 22688 | 1.00 |

Table 11: Summary of the Stan output for the SDE model given experimental data for d2eGFP. The initial time point $t_0$ is not estimated here, but predetermined based on the mean estimate of the sample for the ODE model.

| | mean | c.v. | 2.5% | 50% | 97.5% | $n_{\text{eff}}$ | $\hat{R}$ |
|---|---|---|---|---|---|---|---|
| $\theta_1$ | 0.11 | 0.244 | 0.06 | 0.10 | 0.17 | 1494 | 1.01 |
| $\theta_2$ | 10.36 | 0.292 | 5.14 | 10.18 | 16.77 | 1226 | 1.01 |
| $\theta_3$ | 0.09 | 0.095 | 0.08 | 0.09 | 0.11 | 674 | 1.02 |
| $m_0$ | 13.45 | 0.317 | 7.06 | 12.73 | 23.72 | 954 | 1.01 |
| scale | 4.93 | 0.212 | 3.21 | 4.83 | 7.27 | 785 | 1.01 |
| offset | 8.65 | 0.005 | 8.57 | 8.65 | 8.74 | 22392 | 1.00 |
| $\sigma$ | 0.01 | 0.067 | 0.01 | 0.01 | 0.01 | 13124 | 1.00 |
| $\theta_2 m_0$ | 130.52 | 0.229 | 80.35 | 127.18 | 196.70 | 897 | 1.01 |
| $\theta_2$scale | 49.67 | 0.282 | 27.16 | 48.09 | 82.52 | 838 | 1.01 |
| $m_0$scale | 65.47 | 0.363 | 34.66 | 60.26 | 125.01 | 1343 | 1.01 |
| $\theta_2 m_0$scale | 615.77 | 0.092 | 509.06 | 614.02 | 733.51 | 17424 | 1.00 |

For the parameters $\theta_2$, $m_0$, $\mathrm{scale}$, and their products, the results look somewhat different from those for the eGFP trajectory and those for the simulated data. For the product $\theta_2 m_0 \mathrm{scale}$, the 95% CI for the SDE model is again a lot wider than for the ODE model, but here, the CIs do not overlap. For the parameters $\mathrm{scale}$ and $\theta_2 m_0$, the 95% CI for the SDE model are again a lot narrower than for the ODE model, and we consider them as practically identifiable for the SDE model but not the ODE model. But here, also for the parameters $m_0$, $\theta_2 m_0$, and $m_0 \mathrm{scale}$, the 95% CI for the SDE model are much narrower than for the ODE model, and the parameters seem to be practically identifiable. For parameter $\theta_2$ the 95% CI for the SDE model is slightly wider than for the ODE model, however, the distribution looks different.

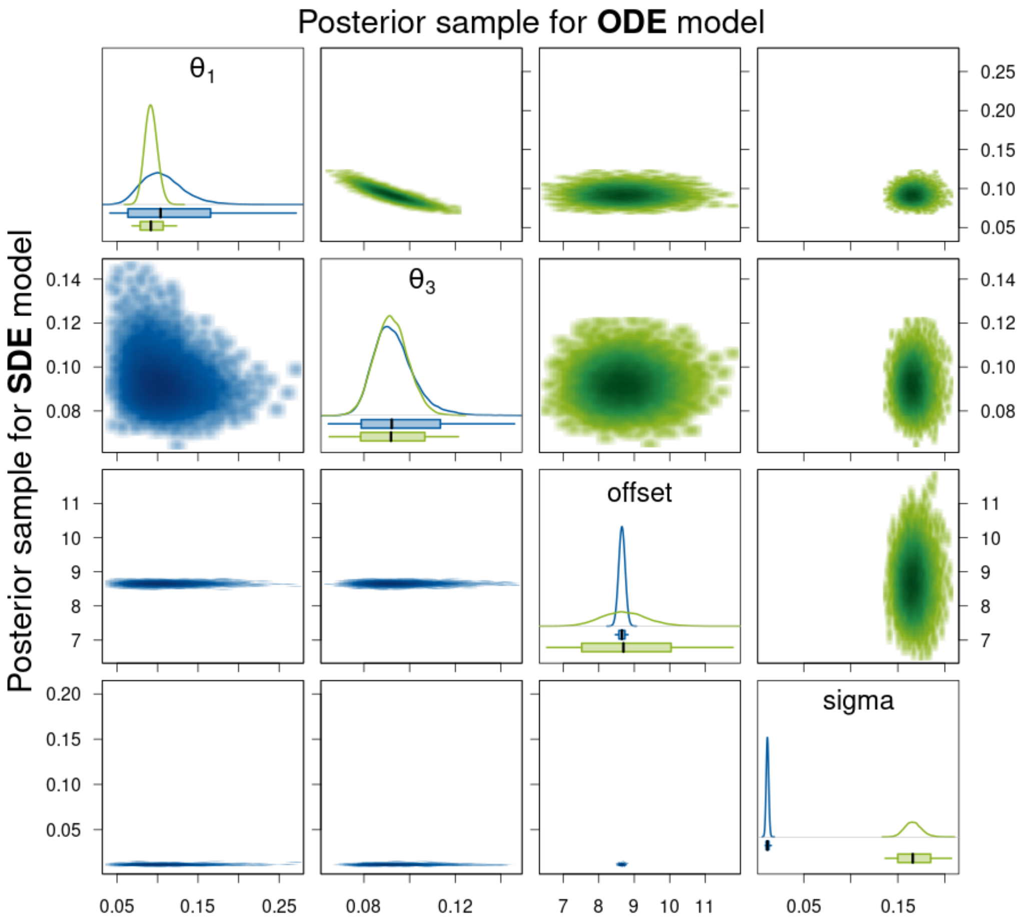


Figure 23: Density estimates of the posterior samples for parameters $\theta_1$, $\theta_3$, $\mathrm{offset}$, and $\sigma$ for the SDE (blue, lower triangle) and ODE (green, upper triangle) model given experimental data for d2eGFP. *Diagonal panels:* Marginal densities for the respective parameter and boxplots showing the 95% CI as box, the range of the sample as whiskers, and the median as thick black line. *Off-diagonal panels:* Smoothed scatter plots of the two-dimensional projections of the samples where darker hues signify higher density values.

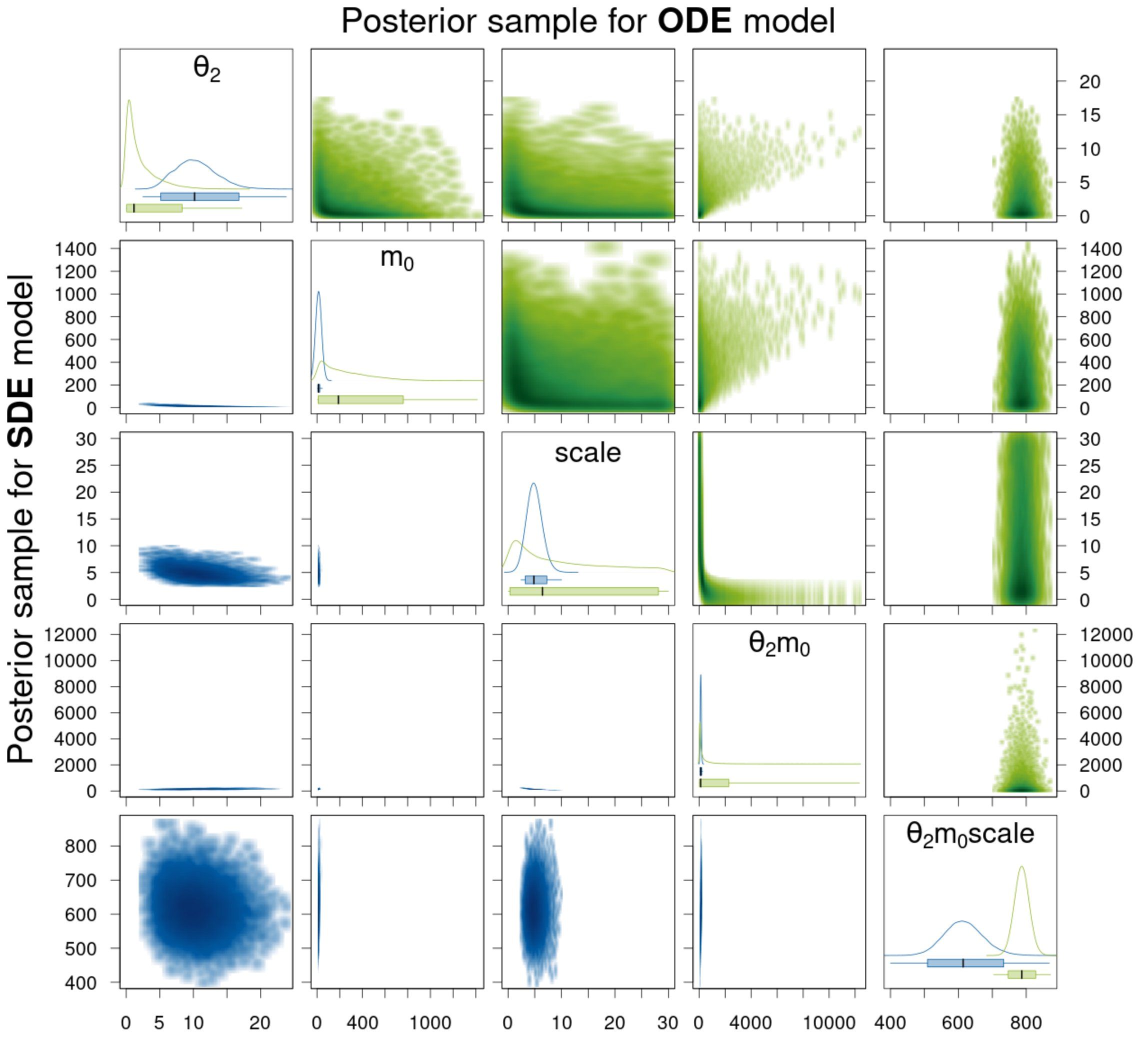


Figure 24: Density estimates of the posterior samples for parameters $\theta_2$, $m_0$, $\mathrm{scale}$, and their products for the SDE (blue, lower triangle) and ODE (green, upper triangle) model given experimental data for d2eGFP. For a detailed description of the figure's elements, see Figure 23.

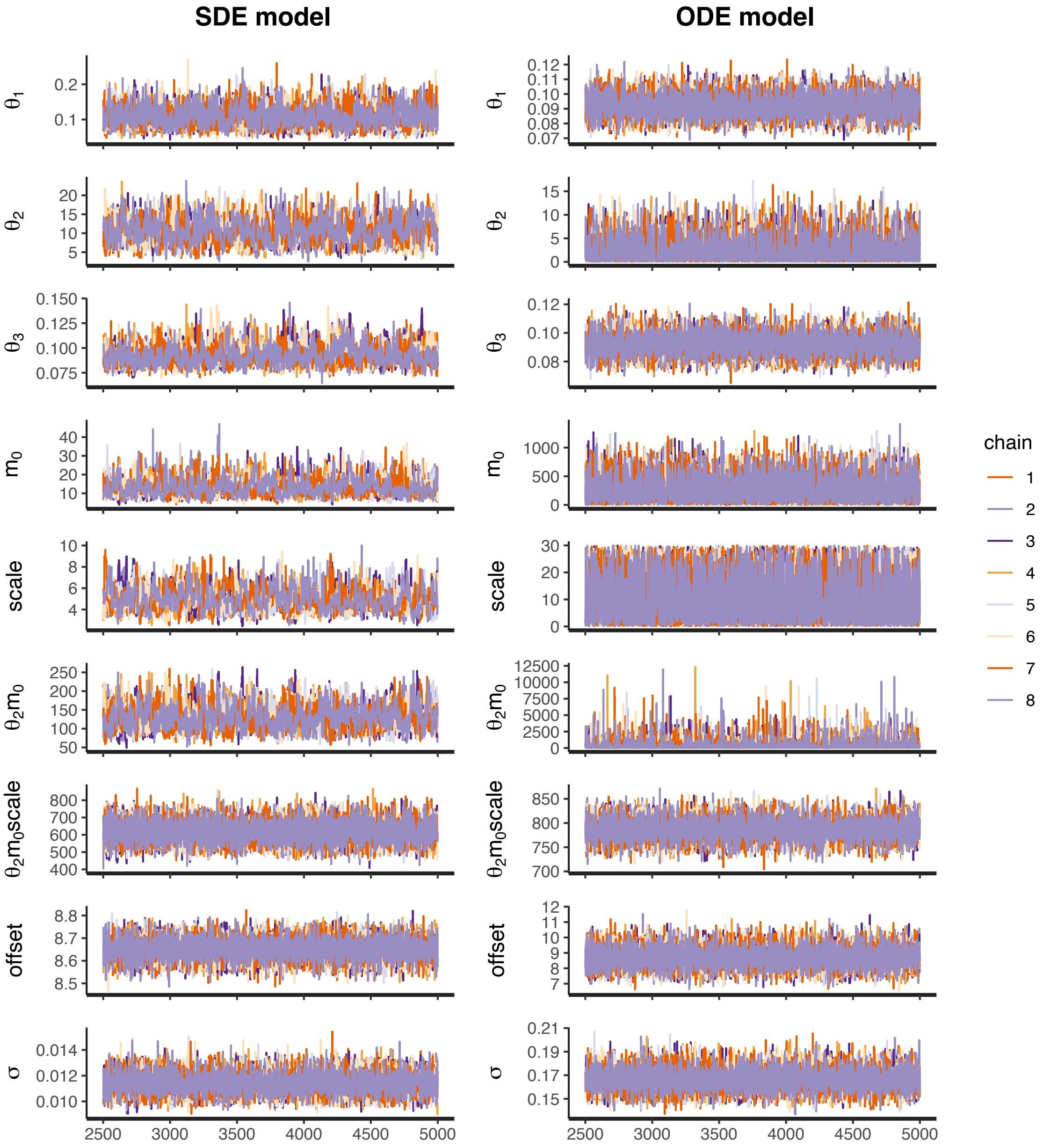


Figure 25: Trace plots of the posterior samples for the experimental data for d2eGFP for the SDE model (left column) and the ODE model (right column). Both posterior samples consist of 8 chains with 2500 warm-up iterations and 2500 iterations after warm-up. Only the iterations after warm-up are displayed.

The statistics of posterior samples aggregated for 100 experimental trajectories for d2eGFP in Table 12 are qualitatively very similar to those for eGFP in Table 9 in the main article. Therefore, we do not repeat the detailed description. We only point out that again unlike for the ODE model, the parameters $\mathrm{scale}$ and $\theta_2 m_0$ are identifiable for the SDE model which is indicated by the much narrower median length of the 95% CIs. We also want to mention that here, the median CI lengths for both degradation rate constants $\theta_1$ *and* $\theta_3$ are smaller for the ODE model than those for the SDE model. This is again due to the fact that for the majority of the observed trajectories the parameter values seem to be very close to each other; and therefore, the two modes of the ODE posterior distribution with respect to these parameters simply overlap. This leads to very narrow CIs which is consistent with our results for the simulated data if we consider the width of the individual modes there. However, we would like to remind the reader that the simulated data also showed that often neither of the modes (and sometimes not even the range of sampled values) covered the true parameter. So assuming that an MJP is the most appropriate description for the generating process of the experimental data, the low uncertainty suggested by narrow CIs for the ODE model might be misleading.

Table 12: Statistics of posterior samples aggregated for 100 experimental trajectories for d2eGFP.

| | | length of prior 95% center interval | median length of 95% CIs | c.v. of lengths of 95% CIs |
|---|---|---|---|---|
| $\theta_1$ | ODE | 11.05 | 0.03 | 0.061 |
| | SDE | 11.05 | 0.12 | 0.012 |
| $\theta_2$ | ODE | 11.05 | 7.88 | 0.006 |
| | SDE | 11.05 | 11.61 | 0.269 |
| $\theta_3$ | ODE | 11.05 | 0.03 | 0.062 |
| | SDE | 11.05 | 0.07 | 0.025 |
| $m_0$ | ODE | 884.82 | 749.92 | 0.181 |
| | SDE | 884.82 | 76.56 | 224.650 |
| $\mathrm{scale}$ | ODE | 28.50 | 27.55 | 0.001 |
| | SDE | 28.50 | 5.51 | 5.550 |
| $\theta_2 m_0$ | ODE | 6056.48 | 2048.33 | 15.537 |
| | SDE | 6056.48 | 172.02 | 173.641 |
| $\theta_2\mathrm{scale}$ | ODE | 228.08 | 67.95 | 0.843 |
| | SDE | 228.08 | 34.96 | 44.862 |
| $m_0\mathrm{scale}$ | ODE | 19271.13 | 9085.56 | 56.326 |
| | SDE | 19271.13 | 482.87 | 3408.541 |
| $\theta_2 m_0\mathrm{scale}$ | ODE | 113232.70 | 40.80 | 30.553 |
| | SDE | 113232.70 | 145.18 | 83.283 |
| $\mathrm{offset}$ | ODE | 28.50 | 2.02 | 1.751 |
| | SDE | 28.50 | 0.79 | 1.029 |
| $\sigma$ | ODE | 9.50 | 0.03 | 0.006 |
| | SDE | 9.50 | 0.01 | 0.005 |

### A.5 Stan specific diagnostics for the sampling output

Here, we summarize the Stan specific diagnostics described in A.3.3 for the HMC output from Sections 6.2 and 6.3. Tables 13 and 14 present the statistics of the number of divergent transition,

Tables 15 and 16 the statistics of the number of times that the user-specified maximal tree depth was exceeded, and Tables 17 and 18 that statistics of the BFMI.

Overall, all three diagnostics show poorer values for the sampling output for the SDE model than for the ODE model. This is not surprising as we sample from a much higher-dimensional distribution for the SDE model. We do not consider the poor diagnostics as a disadvantage of the procedure as they provide information that we do not even have for other MCMC algorithms and thus cannot compare to them.

Table 13: Statistics for the Stan diagnostic of the number of divergent transitions for the SDE model. The 100 sampling outputs per dataset are categorized by the number of divergent transitions that occurred after warm-up, i. e. during a total of 20,000 iterations. Hence, the values in columns 1 to 4 sum to 100. Column 5 gives the maximum number of divergent transitions that occurred after warm-up for one sampling output.

| dataset | none | $1-10$ | $11-100$ | $>100$ | maximum |
|---|---|---|---|---|---|
| simulated data without error | 37 | 10 | 25 | 28 | 1644 |
| simulated data with error | 88 | 4 | 5 | 3 | 568 |
| experimental data for eGFP | 93 | 4 | 3 | 0 | 39 |
| experimental data for d2eGFP | 90 | 3 | 6 | 1 | 540 |

Table 14: Statistics for the Stan diagnostic of the number of divergent transitions for the ODE model. See Table 13 for a detailed description.

| dataset | none | $1-10$ | $11-100$ | $>100$ | maximum |
|---|---|---|---|---|---|
| simulated data without error | 100 | 0 | 0 | 0 | 0 |
| simulated data with error | 100 | 0 | 0 | 0 | 0 |
| experimental data for eGFP | 99 | 1 | 0 | 0 | 1 |
| experimental data for d2eGFP | 92 | 8 | 0 | 0 | 2 |

Table 15: Statistics for the Stan diagnostic of the number of times that the maximal tree depth was exceeded for the SDE model. The user-defined maximal tree depth was set to a value of 15 prior to sampling. The 100 sampling outputs per dataset are categorized by the number of times that the maximal tree depth was exceeded after warm-up, i. e. during a total of 20,000 iterations. Hence, the values in columns 1 to 4 sum to 100. Column 5 gives the maximum number of times that the maximal tree depth was exceeded after warm-up for one sampling output.

| dataset | none | $1-10$ | $11-100$ | $>100$ | maximum |
|---|---|---|---|---|---|
| simulated data without error | 99 | 0 | 1 | 0 | 11 |
| simulated data with error | 10 | 26 | 21 | 43 | 7126 |
| experimental data for eGFP | 25 | 19 | 31 | 25 | 1976 |
| experimental data for d2eGFP | 95 | 2 | 3 | 0 | 59 |

Table 16: Statistics for the Stan diagnostic of the number of times that the maximal tree depth was exceeded for the ODE model. See Table 15 for a detailed description.

| dataset | none | $1-10$ | $11-100$ | $>100$ | maximum |
|---|---|---|---|---|---|
| simulated data without error | 91 | 6 | 0 | 3 | 2500 |
| simulated data with error | 96 | 0 | 0 | 4 | 2500 |
| experimental data for eGFP | 97 | 0 | 0 | 3 | 2500 |
| experimental data for d2eGFP | 100 | 0 | 0 | 0 | 0 |

Table 17: Statistics for the Stan diagnostic $\widehat{BFMI}$ for the SDE model. Each of the 100 sampling outputs per dataset consists of 8 HMC chains for each of which $\widehat{BFMI}$ is calculated. Then, we determine the minimum and the mean over the 8 chains. The table presents the mean and the standard deviation (s.d.) of these minima and means aggregated over the 100 sampling outputs per dataset.

| dataset | mean of minima | s.d. of minima | mean of means | s.d. of means |
|---|---|---|---|---|
| simulated data without error | 0.03 | 0.01 | 0.05 | 0.01 |
| simulated data with error | 0.05 | 0.02 | 0.07 | 0.01 |
| experimental data for eGFP | 0.05 | 0.04 | 0.08 | 0.04 |
| experimental data for d2eGFP | 0.07 | 0.05 | 0.09 | 0.05 |

Table 18: Statistics for the Stan diagnostic $\widehat{BFMI}$ for the ODE model. See Table 17 for a detailed description.

| dataset | mean of minima | s.d. of minima | mean of means | s.d. of means |
|---|---|---|---|---|
| simulated data without error | 0.95 | 0.19 | 1.03 | 0.06 |
| simulated data with error | 0.95 | 0.15 | 1.03 | 0.06 |
| experimental data for eGFP | 0.94 | 0.19 | 1.03 | 0.05 |
| experimental data for d2eGFP | 0.90 | 0.23 | 1.02 | 0.05 |
